# The Converging Effects of Different Categories of Antidepressants on the Brain: A Systematic Meta-Analysis of Public Transcriptional Profiling Data from the Hippocampus and Cortex

**DOI:** 10.1101/2025.04.21.648805

**Authors:** Eva M. Geoghegan, Megan H. Hagenauer, Erin Hernandez, Sophia Espinoza, Elizabeth Flandreau, Phi T. Nguyen, Adrienne N. Santiago, Mubashshir Ra’eed Bhuiyan, Sophie Mensch, Stanley Watson, Huda Akil, René Hen

**Author notes:** **Corresponding author:** Megan Hagenauer, Michigan Neuroscience Institute, University of Michigan, Ann Arbor, MI 48109. Joint first authorship: These authors have contributed equally to this work (Eva M. Geoghegan, Megan H. Hagenauer).

## Abstract

Depression can be treated with traditional pharmaceuticals targeting monoaminergic function, non-traditional drug classes and neuromodulatory interventions. To identify mechanisms of action shared across clinically-effective antidepressant treatment categories, we performed two systematic meta-analyses of public transcriptional profiling data from adult laboratory rodents (rats, mice). The outcome variable was gene expression, measured by microarray or RNA-Seq from bulk-dissected tissue from two depression-related brain regions (hippocampus, cortex). Relevant datasets were identified in the Gemma database of curated, reprocessed transcriptional profiling data using predefined search terms and inclusion/exclusion criteria (*hippocampus:* 6-24-2024, *cortex:* 7-10-2024). Differential expression results were extracted for all genes, minimizing bias. For each gene, a random effects meta-analysis model was fit to antidepressant vs. control effect sizes (Log2 Fold Changes) from each study for each brain region, with follow-up analyses exploring sources of effect heterogeneity. For the hippocampus, 15 relevant studies were identified, containing 22 antidepressant vs. control group comparisons (collective *n*=313 samples), with approximately half representing traditional versus non-traditional antidepressants. Of 16,439 analyzed genes, 58 were consistently differentially expressed (False Discovery Rate (FDR)<0.05) following treatment. Antidepressant effects were enriched in the dentate gyrus and in gene sets related to stress regulation, brain growth and plasticity, vasculature and glia, and immune function. Comparisons with single nucleus RNA-Seq confirmed effects on specific hippocampal cell types, including potential rejuvenation of dentate granule neurons. For the cortex, 13 studies were identified, containing 16 antidepressant vs. control group comparisons (collective *n*=233 samples). Of 15,583 analyzed genes, only one was consistently differentially expressed (FDR<0.05: *Atp6v1b2*), but overall expression patterns moderately resembled the hippocampus. These genes and pathways showing consistent differential expression across treatment categories may be promising targets for novel therapies. Future work should explore relevance to human clinical populations and potential heterogeneity introduced by sex and subregion.

**Key Points:**

- Depression can be treated with traditional antidepressants targeting monoaminergic function, as well as other drug classes and non-pharmaceutical interventions.
- Understanding the congruent effects of different types of antidepressant treatments on sensitive brain regions, such as the hippocampus and cortex, can highlight essential mechanisms of action.
- A meta-analysis of public transcriptional profiling datasets identified genes and functional gene sets that are differentially expressed across antidepressant categories.

**Plain Language Summary:** Major depressive disorder is characterized by persistent depressed mood and loss of interest and pleasure in life. Worldwide, an estimated 5% of adults suffer from depression, making it a leading cause of disability. The current standard of care for depressed individuals includes psychotherapy and antidepressant medications that enhance signaling by monoamine neurotransmitters, such as serotonin and norepinephrine. Other treatments include non-traditional antidepressants that function via alternative, often unknown, mechanisms. To identify mechanisms of action shared across different categories of antidepressants, we performed a meta-analysis using public datasets to characterize changes in gene expression (mRNA) following treatment with both traditional and non-traditional antidepressants. We focused on the hippocampus and cortex, which are two brain regions that are sensitive to both depression and antidepressant usage. We found 59 genes that had consistently higher or lower levels of expression (mRNA) across antidepressant categories. The functions associated with these genes were diverse, including regulation of stress response, the immune system, brain growth and adaptability. These genes are worth investigating further as potential linchpins for antidepressant efficacy or as targets for novel therapies.

**Graphical Abstract:** 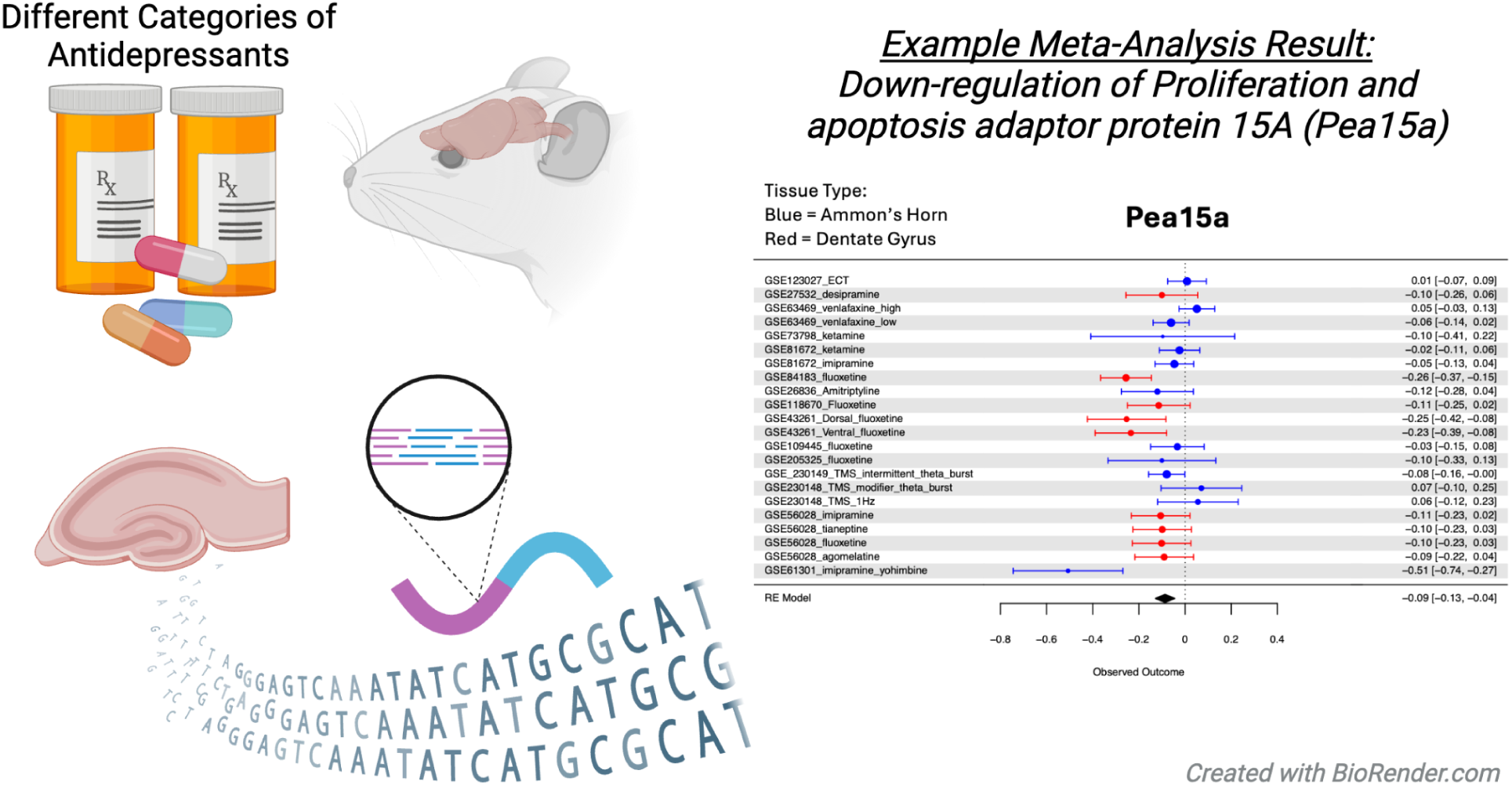

## Introduction

Major Depressive Disorder (MDD) is characterized by consistent depressed mood or markedly diminished pleasure in daily activities ^1^. Worldwide, an estimated 5% of adults suffer from depression, making it a leading cause of disability ^2,3^ and increased mortality ^4^. The current standard of care includes evidence-based psychotherapies and specific classes of drugs, which may be guided by blood biomarkers predictive of treatment outcomes ^5,6^. Although antidepressant treatments are effective for some patients, around 30-40% of patients fail to respond to first-line treatment ^7^.

To improve individualized treatment options, the mechanisms of action for different antidepressant treatments need to be investigated further to understand both their congruent and differing neurobiological effects. Many first-line antidepressants enhance monoamine neurotransmitter function, such as the serotonergic, noradrenergic, and dopaminergic systems. These include selective serotonin reuptake inhibitors (SSRIs), serotonin-norepinephrine reuptake inhibitors (SNRIs), tricyclic antidepressants (TCAs) and monoamine oxidase inhibitors (MAOIs) ^8–10^. However, monoamine potentiation may not be the direct mechanism of action of these drugs, as monoamine potentiation occurs rapidly and clinical improvement usually requires several weeks ^9^.

Other classes of pharmacological treatments have shown promise for treating depression. Ketamine, a N-Methyl-D-aspartate receptor (NMDAR) antagonist, has rapid and long-lasting antidepressant effects at subanesthetic doses ^11–13^. Agomelatine has synergistic effects as an agonist of melatonin and some serotonin receptors ^7,14,15^. Tianeptine is an atypical tricyclic believed to have antidepressant effects via μ-opioid receptor agonism ^16,17^. Other promising new classes include psychedelics (MDMA, psilocybin)^18^ and second generation antipsychotics (e.g., quetiapine)^19^. Finally, alpha-2 adrenergic antagonists may accelerate TCA treatment response times by reversing treatment-related alpha-2 adrenoceptor desensitization ^20,21^. Depression is also treated using neuromodulatory brain stimulation therapies, such as electroconvulsive therapy (ECT), deep brain stimulation (DBS) and transcranial magnetic stimulation (TMS) ^22–27^. Altogether, the diversity of neurobiological pathways targeted by these new treatments suggests that monaminergic effects may not be central to antidepressant response, or may lie upstream of other critical pathways.

In addition to diverse molecular targets, there are multiple brain regions that may mediate antidepressant response. Antidepressant treatments are often theorized to alleviate depressive symptoms via effects on the hippocampus. Hippocampal volume reduction is common in patients with depression ^28^. This atrophy may be due to prolonged exposure to stress hormones (glucocorticoids) and subsequent reduction in neurotrophic factors and adult hippocampal neurogenesis (AHN) in the dentate gyrus (DG) ^28^, both of which precede mood disturbance ^29–31^. Antidepressants reverse or prevent hippocampal atrophy in depression. Growing evidence suggests that both traditional, monoaminergic-targeting antidepressants (*e.g.,* SSRIs, TCAs) ^32,33^ and other antidepressant treatments (*e.g.,* ECT) ^34,35^ reverse hippocampal atrophy by stopping stress-induced dendritic retraction, increasing growth factor signaling (*e.g.,* brain-derived neurotrophic factor (BDNF) ^11,14,15,36^, and increasing hippocampal neuroplasticity and AHN ^32,37,38^. However, it is still unclear whether increases in hippocampal neuroplasticity and AHN are a central mechanism for all antidepressant classes.

Some antidepressants may also target the cortex. Depressed patients have documented cortical abnormalities, including thinner cortical gray matter ^39^ and reduced prefrontal cortex (PFC) volume ^40^. Rodent models of depression show similar structural and physiological changes, including dendritic shrinkage and reductions in myelination in the PFC ^41^. Antidepressant treatments may reverse these changes by directly modulating cortical function. For example, TMS is thought to facilitate synaptic plasticity within the target site, which is most commonly the dorsolateral prefrontal cortex (DLPFC) ^26^ , by increasing cortical activity ^42^, and antidepressant DBS targets cortical white matter tracts ^22^. Antidepressant pharmaceuticals can also modulate cortical activity. As an NMDA receptor antagonist, ketamine decreases cortical GABAergic interneuron activity, causing downstream disinhibition ^12^ and increasing synaptic plasticity ^11–13^. Similarly, monoaminergic antidepressants (SSRIs, SNRIs) have been shown to increase activity or metabolism in frontal cortical regions ^43^. However, whether cortical plasticity is a central mechanism for all antidepressant classes remains unknown.

In order to better understand the effects of antidepressant treatments on the hippocampus and cortex, we conducted two systematic meta-analyses of publicly available rodent transcriptional profiling data, as well as a variety of follow-up exploratory analyses. Transcriptional profiling technologies, such as ribonucleic acid-sequencing (RNA-Seq) and microarray, quantify gene expression in tissues. By conducting meta-analyses across studies using different classes of antidepressants, we hoped to identify patterns of gene expression common to a wide variety of antidepressant treatments while also increasing statistical power to provide stronger insights into gene expression patterns underlying functional changes.

## Methods

The meta-analysis project was conducted using a standardized pipeline for planned analyses (protocol: ^44^, validation: ^45^, *not pre-registered*). Additional exploratory analyses are denoted in **Fig 1** and the text, with all analysis code (R v.4.4.1 and RStudio v.2024.9.9.365 ^46,47^) available at: https://github.com/evageoghegan/AntidepressantMetaAnalysis.

**Figure 1.**
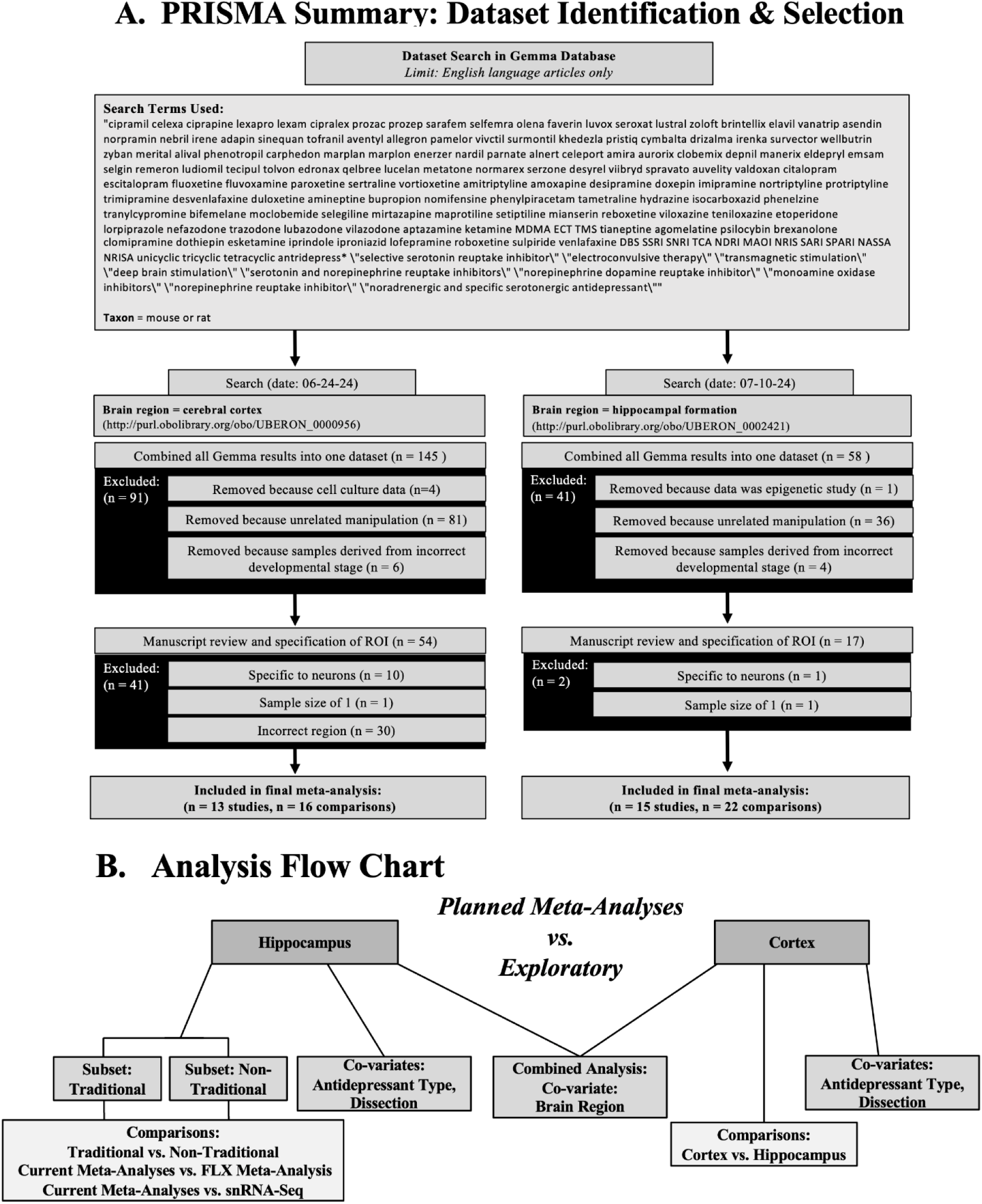
A PRISMA diagram overviewing hippocampal and cortical dataset search and selection. **A.** We identified transcriptional profiling datasets of laboratory rodents treated with antidepressants within the Gemma database using pre-specified keywords. The search encompassed tissue from the hippocampal formation. The titles, abstracts, and metadata for the datasets were initially scanned and filtered using pre-specified inclusion/exclusion criteria, including indications that the dataset was an epigenetic study, had a manipulation unrelated to antidepressants, or had samples originating from the incorrect developmental stage or cell culture. The secondary screening step included a detailed review of the metadata on Gemma and published methodology, followed by a final specification of the region of interest (ROI: hippocampus or cortex). Datasets were excluded in the secondary dataset filtering if they did not include a sufficient sample size, were cell type-specific or sampled from a different region. **B.** A flow chart overviewing the analyses, including planned meta-analyses examining the converging effects of antidepressants on the hippocampus and cortex, and exploratory analyses comparing effects within particular antidepressant categories (traditional monoaminergic-targeting vs. non-traditional mechanisms), dissections (dentate gyrus (DG) vs. hippocampus, anterior cingulate (ACG) vs. cortex, prefrontal cortex (PFC) vs. cortex), and brain regions (cortex vs. hippocampus)). Exploratory analyses also compared meta-analysis results to previous work, including a previous meta-analysis examining the effects of FLX on the hippocampus ^58^ and snRNA-Seq studies examining the effect of FLX and ECT on hippocampal cell types ^59,60^. <u>Abbreviations:</u> FLX: fluoxetine, ECT: Electroconvulsive therapy, DG: Dentate gyrus, ACG: Anterior cingulate cortex, PFC: Prefrontal cortex, ROI: region of interest, n=number of datasets

### Information source

The Gemma database was used to access curated and reprocessed public transcriptional profiling data ^48,49^. Gemma contains differential expression results for nearly 20,000 publicly available transcriptional profiling datasets, representing approximately 600,000 samples ^49,50^. We focused on this database to provide consistency in our meta-analysis input: Gemma adheres to a strict, standardized pipeline for dataset curation, preprocessing, and re-analysis ^48,49^. During preprocessing, Gemma realigns the datasets to an updated genome. Quality control measures include identification and removal of outlier samples, removal of genes (rows) with minimal variance in expression values (either zero variance or <70% distinct values), and manual curation for common issues such as batch effects ^48^. Differential expression is calculated using the *limma* or *limma-voom* pipeline, and statistical output is available for the full model (omnibus) and individual contrasts.

### Dataset identification and search

For our meta-analysis, we focused on treatments that specifically target depressive mood symptoms ^51^. A list of common traditional antidepressants, corresponding brand names and drug classes was compiled (**Fig. 1**). Non-traditional antidepressant treatments were added if there was approved clinical use or high-quality peer-reviewed clinical evidence available for potential antidepressant effects. These included ECT, TMS, DBS, ketamine, 3,4-methylenedioxymethamphetamine (MDMA), psilocybin, tianeptine, quetiapine, and agomelatine ^14,18,19,23,27,52–55^. The scope for the search included transcriptional profiling datasets from laboratory rats (*Rattus norvegicus*) or laboratory mice (*Mus musculus*) using bulk tissue dissections that targeted the full or large representative portion of the transcriptome. Search terms were initially generated by three of the authors (EMG, EH, and SE) and decisions reviewed and approved by co-authors (MHH, RH, SE, MRB, SM). The search terms were then structured with Boolean operators (PRISMA diagram: **Fig. 1**) and the search conducted using Gemma’s application programming interface (API, packages *gemma.R* (v.3.1.6), *plyr* (v.1.8.9) and *dplyr* (v.1.8.9) ^49,56,57^).

### Initial filtering

The metadata for the identified datasets was filtered by “organism part” using the formal hierarchical ontology term for the hippocampus (UBERON_0002421) or cerebral cortex (UBERON_0000956). Metadata records were then filtered down to publicly available datasets that were not labeled as problematic within the Gemma database. The identified datasets were scrutinized for inclusion/exclusion based on systematic criteria ^44^. Initial filtering for the hippocampal meta-analysis was performed by the first author (EMG), and then decisions were reviewed and approved by co-authors (MHH, RH). Forty-one studies were removed because either 1) treatment was administered during development, 2) brain collection was during development, 3) the focus was epigenetic, or 4) the manipulation was unrelated to antidepressant treatment. Initial filtering for the cortical meta-analysis was performed by two researchers (SE, EMG), and then decisions were reviewed and approved by co-authors (MHH, RH, MRB). Ninety-one studies were removed because either 1) the manipulation was unrelated to antidepressant treatment, 2) brain collection was during development, or 3) the studies were in cell culture.

### Secondary filtering

For the remaining studies, metadata for the differential expression results were downloaded and reviewed to ensure that there was an experimental manipulation relevant to antidepressant treatment and related statistical contrasts. The publications associated with the datasets were also referenced, focusing exclusively on methodological information to determine inclusion/exclusion. Following this review, two datasets were removed from the hippocampal meta-analysis: one due to being neuron-specific and another due to having a sample size of *n*=1 per subgroup. For the cortical meta-analysis, forty-one datasets were removed: ten due to being neuron-specific, thirty due to focus on non-cortical regions, and one due to having a sample size of *n*=1 per subgroup. At this stage, all inclusion/exclusion decisions were reviewed and finalized by the entire 2024 cohort of the *Brain Data Alchemy Project* (MHH, SE, EIF, SM, RB, LTC, AL, DMN, TQD) and laboratory principal investigator (RH).

### Reprocessing

Four studies required reprocessing: GSE118670, GSE26836, GSE84183 and GSE43261. For GSE118670, GSE26836, and GSE43261, the differential expression model used by Gemma included multiple tissue types and re-analysis was necessary to extract tissue-specific differential expression results. For GSE43261, the differential expression model also used treatment sensitivity phenotype as the variable of interest rather than the treatment itself. For GSE84183, the expression data appeared to have been log(2) transformed twice during Gemma’s preprocessing, showing a highly reduced range.

To re-analyze these datasets, we followed Gemma’s analysis pipeline as much as possible to maintain consistency. We used the *Gemma.R* (v.0.99.41) and *tidyverse* (v.2.0.0) ^48,50^ packages to import preprocessed log(2) expression data and sample metadata from the Gemma database (import date: 10/22/2024). Each dataset was subsetted to only our brain regions of interest, and the Gemma Diagnostics tab was consulted to determine whether outlier samples or batch effects were previously identified (*none found*). For GSE118670, adult and developing comparison mice were also excluded, as they did not receive antidepressant or control treatment conditions.

Genes (rows) were excluded that lacked log(2) expression data or had insufficient variability within the antidepressant or control groups. The control group for the antidepressant treatment was defined as the treatment factor reference level (intercept). Any additional experimental variables present in the metadata were included as co-variates (GSE84183: stress treatment, with a reference level of no stress, GSE118670, GSE43261, and GSE26836: no co-variates). The differential expression model was fit using the *limma* pipeline (package: *limma* v.3.17) with correction for heteroskedasticity (mean-variance trend) and an empirical Bayes correction ^61^.

### Result extraction & meta-analysis

Differential expression results for the antidepressant treatment vs. control statistical contrasts from each included study were extracted. The Log(2) Fold Change (Log2FC) was extracted for each gene from each study, and standard error (SE) and sampling variance calculated. If more than one result represented a gene, the Log2FC and SE were averaged. Results were then aligned across species using EntrezID and a version of the Jackson Labs Mouse Ortholog Database ^62^ that had been trimmed to one-to-one orthologs (*Meta-Analysis Input:* **Table S1**). For a gene to be included in the hippocampal or cortical meta-analysis, a minimum number of 11 results (contrasts) needed to be available. A simple (intercept-only) random effects meta-analysis was then fit to the Log2FC estimates from each contrast (package: *metafor* ^63^ (v.4.8.0)). A false discovery rate (FDR) correction using the Benjamin-Hochberg method was applied to the nominal meta-analysis (two-tailed) p-values (*multtest* v.1.32.0) 64. Differentially expressed genes (DEGs) were defined using a threshold of FDR<0.05, and visualized using forest plots (package: *metafor* 63 (v.4.8.0)).

### Assessment of meta-analysis result robustness and validity

To evaluate the robustness and validity of the meta-analysis results, we examined the potential impact of publication bias and influential outlier studies. Publication bias was assessed using Egger’s regression analysis (function *regtest()* in *metafor* ^63^ v.4.8.0), which detects funnel plot asymmetry^65^ . Nominal Egger test (two-tailed) p-values were corrected for FDR using the Benjamin-Hochberg method (*multtest* v.1.32.0) 64. Genes with Egger FDR<0.05 were considered to show evidence of potential publication bias. To assess the influence of individual studies on meta-analysis results, the meta-analysis was repeated while iteratively excluding each of the study contrasts (*leave1out()* function in *metafor* 63 v.4.8.0). If a previously-identified DEG was found to drop below a nominal significance threshold (*p*>0.05) following the exclusion of any of the study contrasts, the finding was considered to not be robust. To determine which of the study contrasts were disproportionately influential on the meta-analysis results, we outputted Cook’s Distances^66^ (Cook’s D) and Difference in Betas (DFBetas), as well as a general assessment of overall influence (TRUE/FALSE) as provided by the *influence()* function (*metafor* 63 v.4.8.0).

### Assessment or residual heterogeneity

To assess residual heterogeneity, we extracted statistics from the meta-analysis object (*metafor* 63 v.4.8.0) estimating the residual variance in the underlying true antidepressant effect sizes (Log2FCs) observed across study contrasts, either in absolute terms (Tau^2^ and associated SE) or as a percent (I^2^) or ratio (H^2^) of the total variability. To determine whether this residual heterogeneity exceeded what might be expected due to random sampling variability, we extracted the Cochran’s Q test statistic and associated nominal (two-tailed) p-value^67^, which was corrected for FDR using the Benjamin-Hochberg method (*multtest* v.1.32.0) ^64^. Genes with a Q-test FDR<0.05 were considered to show significant heterogeneity across studies.

### Exploratory follow-up analyses

In order to explore the effects of different treatment categories on the hippocampus, two subgroup meta-analyses were run using only the contrasts characterizing the effects of: 1) traditional, monoaminergic-targeting antidepressants, 2) non-traditional antidepressants (**Table 1**). Based on dataset availability, for a gene to be included in the traditional subgroup meta-analysis, differential expression results needed to be available from a minimum of 12 antidepressant vs. control comparisons, whereas for the non-traditional subgroup meta-analysis the minimum was 7 antidepressant vs. control comparisons. The effects of traditional versus non-traditional antidepressants were further compared using a hippocampal meta-regression that included antidepressant type (non-traditional vs. traditional) and dissection (dentate gyrus vs. hippocampus) as co-variates, with the general antidepressant effect (intercept) centered between the levels for each variable (*i.e.,* at the average of the cell means: *contrast.sum*). This meta-regression was designed to control for the correlation between antidepressant type and dissection method, as a larger percent (6 of 8: 75%) of the statistical contrasts for the non-traditional antidepressants used whole hippocampal tissue than for the traditional antidepressants (6 of 13: 46%). Two additional exploratory meta-regressions were run that included the variables of platform (microarray vs. RNA-seq) and depression model inclusion in the study (versus control-only studies), but both variables were found to have minimal impact in our design (*see Supplement*).

**Table 1.** Overview of datasets included in the meta-analyses. Color refers to datasets included in specific meta-analyses: hippocampal (yellow), hippocampal and cortical (green), cortical (blue). Experiment ID refers to the Gene Expression Omnibus (GEO) accession number for each dataset. The Author column refers to the first author of the publication where available ^68–88^. The Year column refers to the dataset release date listed on Gemma. All experimental details describe the samples used in the transcriptional profiling experiment (e.g., species, biological sex). The Total n indicates the number of tissue samples in the full dataset, whereas the HPC n and CTX n indicate the number of hippocampal and cortical samples included in the full (omnibus) statistical model for each brain region, respectively, with the sample size in parentheses indicating the number of samples represented in the antidepressant treatment-related statistical contrasts (if different from the full model). The Platform column indicates the transcriptional profiling technology used. The Tissue column indicates the specific region/subregion sampled. Model of Stress/Depression lists relevant subject characteristics or chronic stress conditions included in the experiment (acute stress due to behavioral assays or routine procedures not noted). Treatment column lists each treatment group (with # of samples) included in the treatment vs. control statistical contrasts used in the meta-analysis. Treatment categories included SSRIs (FLX, sertraline), TCAs (imipramine, desipramine, amitriptyline), SNRIs (venlafaxine, duloxetine), NMDA receptor antagonists (ketamine), Alpha-2 adrenergic antagonists combined with TCAs (yohimbine and imipramine), atypical antidepressants (tianeptine), MASSAs (agomelatine), atypical antipsychotics (quetiapine), serotonin–norepinephrine–dopamine releasing agent (SNDRA) (MDMA), and neuromodulatory brain stimulation (ECT, TMS). Within all hippocampal datasets (yellow and green), bolded treatments indicate inclusion in the traditional-only subanalysis and italicized treatments indicate inclusion in the non-traditional only subanalysis. <u>Abbreviations</u>: HPC: Hippocampus, DG: Dentate Gyrus, PFC: Prefrontal Cortex, ACG: Anterior Cingulate Cortex, FLX: fluoxetine, Ctrl: Control, CUMS: chronic unpredictable mild stress, CSDS: chronic social defeat stress, OBX: olfactory bulbectomy, LPS: lipopolysaccharide (LPS)-induced depression-like model, SIA: stress-induced analgesia, SSRI: selective serotonin reuptake inhibitor, NMDA: N-Methyl-D-aspartate, SNRI: serotonin-norepinephrine reuptake inhibitor, TCA: Tricyclic antidepressant, MASSAs: melatonin agonist and selective serotonin antagonist, SNDRA: serotonin–norepinephrine–dopamine releasing agent, MDMA: 3,4-Methylenedioxymethamphetamine, ECT: Electroconvulsive therapy, TMS: Transcranial magnetic stimulation

| Dataset ID | Total<br>*n* | HPC<br>*n* | CTX<br>*n* | Species | Sex | Authors | Year | Platform | Tissue | Model of Stress/<br>Depression | Treatment |
| --- | --- | --- | --- | --- | --- | --- | --- | --- | --- | --- | --- |
| GSE109445 | 12 | 12<br>(9) | NA | Rat | M | Wang *et al.*, | 2018 | Illumina<br>(RNA-Seq) | Ammon's horn | CUMS, Ctrl | **FLX (3), Ctrl (6)** |
| GSE123027 | 24 | 24 | NA | Mouse | M | Rimmerman *et al.*, | 2021 | Illumina<br>(RNA-Seq) | Ammon's horn | CUMS | *ECT (10), Ctrl (14)* |
| GSE205325 | 21 | 21 | NA | Rat | M | Demin *et al.*, | 2022 | Illumina<br>(RNA-Seq) | Ammon's horn | CUMS,<br>CUMS-LPS, Ctrl | **FLX (9), Ctrl (12)** |
| GSE230148 | 40 | 40 | NA | Rat | M | Weiler *et al.*, | 2023 | Agilent<br>(microarray) | Ammon's horn | Aged w/ or w/o<br>cog. impairment*,<br>Adult Ctrl | *TMS-mod (14), TMS-1 Hz (12),*<br>*Ctrl (14)* |
| GSE230149 | 56 | 28 | 28 | Rat | M | Weiler *et al.*, | 2023 | Agilent<br>(microarray) | Ammon's horn,<br>Parietal cortex | Aged w/ or w/o<br>without cog.<br>impairment*, Adult<br>Ctrl | <u>HPC</u>: TMS (13), Ctrl (15)<br><u>CTX</u>: TMS (14), Ctrl (14) |
| GSE27532 | 16 | 16 | NA | Mouse | M | Lisowski *et al.*, | 2011 | Illumina<br>(microarray) | DG | Bred for High/ Low<br>Stress Analgesia | **Desipramine (8), Ctrl (8)** |
| GSE56028 | 21 | 21 | NA | Rat | M | Patricio *et al.*, | 2015 | Agilent<br>(microarray) | DG | CUMS, Ctrl | **Imipramine (3), FLX (3),**<br>**Agomelatine (3),**<br>**Tianeptine (3), Ctrl (9)** |
| GSE61301 | 8 | 8 | NA | Rat | M | Husain *et al.*, | 2015 | Agilent<br>(microarray) | Ammon's horn | Ctrl | Imipramine+yohimbine (4), Ctrl<br>(4) |
| GSE63469 | 6 | 6 | NA | Mouse | M | Proft *et al.*, | 2019 | Affymetrix<br>(microarray) | Ammon's horn | Stress sensitive<br>(DBA/2) mice | **Venlafaxine- 100kg/d/kg (2),**<br>**Venlafaxine- 30kg/d/kg (2), Ctrl**<br>**(2)** |
| GSE73798 | 60 | 60<br>(24) | NA | Mouse | M | Ficek *et al.*, | 2019 | Illumina<br>(microarray) | Ammon's horn | Ctrl | *Ketamine (12), Ctrl (12)* |
| GSE81672 | 99 | 24 | 25 | Mouse | M | Bagot *et al.*, | 2018 | Agilent<br>(microarray) | Ammon's horn,<br>PFC | CSDS, Ctrl | <u>HPC</u>: Ketamine (6), Imipramine<br>(7), Ctrl (11)<br><u>CTX</u>: Ketamine (6), Imipramine<br>(7), Ctrl (12) |
| GSE43261 | 38 | 38 | NA | Mouse | M | Samuels *et al.*, | 2014 | Affymetrix<br>(microarray) | DG | CORT* | <u>HPC-ventral</u>: FLX (11),<br>Ctrl (8)<br><u>HPC-dorsal</u>: FLX (11), Ctrl (8) |
| GSE84183 | 64 | 32 | 32 | Mouse | M | Hervé *et al.*, | 2017 | Agilent<br>(microarray) | DG, ACG | CUMS*, Ctrl | <u>HPC</u>: FLX (16), Ctrl (16)<br><u>CTX</u>: FLX (16), Ctrl (16) |
| GSE118670 | 44 | 16 | 28<br>(16) | Mouse | M | Hagihara *et al.*, | 2019 | Affymetrix<br>(microarray) | DG, PFC | Ctrl | <u>HPC</u>: FLX (8), Ctrl (8)<br><u>CTX</u>: FLX (8), Ctrl (8) |
| GSE26836 | 12 | 6 | 6 | Mouse | M | Chadwick *et al.*, | 2015 | Illumina<br>(microarray) | Ammon's horn,<br>Frontal cortex | Aged Alzheimer's<br>genetic model* | <u>HPC</u>: Amitriptyline (3), Ctrl (3)<br><u>CTX</u>: Amitriptyline (3), Ctrl (3) |
| GSE28644 | 60 | NA | 60 | Mouse | M | Benton *et al.*, | 2012 | Affymetrix<br>(microarray) | Cerebral cortex | 30 inbred strains | FLX (30), Ctrl (3) |
| GSE93041 | 9 | NA | 9 (6) | Mouse | M | Orozco-Solis<br>*et al.*, | 2018 | Affymetrix<br>(microarray) | ACG | Ctrl | Ketamine (3), Ctrl (3) |
| GSE150264 | 36 | NA | 36 | Mouse | F | Rajkumar *et al.*, | 2020 | Agilent<br>(microarray) | ACG | Genetic depression<br>model, Ctrl | Imipramine (19), Ctrl (17) |
| GSE168172 | 15 | NA | 15 | Mouse | M | Funayama *et al.*, | 2022 | Illumina<br>(RNA-Seq) | PFC | Stress sensitive<br>(BALB) mice | Duloxetine (6),<br>Sertraline (6), Ctrl (3) |
| GSE138802 | 6 | NA | 6 | Mouse | M | Kokkinou *et al.*, | 2021 | Illumina<br>(RNA-Seq) | PFC | Ctrl | Ketamine (3), Ctrl (3) |
| GSE129359 | 24 | NA | 24<br>(12) | Mouse | M | Lukić *et al.*, | 2019 | Illumina<br>(RNA-Seq) | PFC | Ctrl | Duloxetine (6), Ctrl (6) |
| GSE45229 | 20 | NA | 10<br>(6) | Mouse | M | Kondo *et al.*, | 2013 | Affymetrix<br>(microarray) | Frontal cortex | Ctrl | Quetiapine-10 mg/kg (2),<br>Quetiapine-100 mg/kg (2), Ctrl (2) |
| GSE253280 | 36 | NA | 18<br>(12) | Rat | M | Warner-Schmidt, *et al.*, | 2024 | Illumina<br>(RNA-Seq) | Frontal cortex | Ctrl | MDMA (6), Ctrl (6) |

To further explore differences in antidepressant effects due to brain region, the contrasts from both hippocampus and cortex were included in a larger meta-regression model that included region (hippocampus vs. cortex) as a co-variate, with the general antidepressant effect (intercept) centered at the average of the cell means for the two regions (*contrast.sum*). For the cortical data, an exploratory meta-regression was also performed to examine the effects of antidepressant type (traditional vs. non-traditional) while controlling for dissection (either ACG vs. other cortex or PFC vs. other cortex), but the small number of available contrasts suggested that we were underpowered to properly address this question (*see Supplement*).

To compare the pattern of antidepressant differential expression identified in the different meta-analyses (*e.g.,* planned hippocampal and cortical meta-analyses, exploratory traditional and non-traditional antidepressant meta-analyses), Pearson’s product moment correlation was calculated and visualised using scatterplots, and ranked gene lists compared using Spearman’s rank correlation. To explore other potential sources of heterogeneity, hierarchically clustered heatmaps were created using the Log2FCs from each of the included studies for either all genes included in the hippocampal and cortical meta-analyses or the following ranked gene lists: planned hippocampal meta-analysis (all 58 significant DEGs), planned cortical meta-analysis and exploratory subgroup meta-analyses (top 50 genes, as ranked by p-value).

### Functional patterns

Gene set enrichment analysis was used to determine whether the hippocampal and cortical meta-analysis results, ranked by effect size (Log2FC), were enriched with down-regulation or upregulation in sets of genes associated with known biological pathways and previous experimental results. Both directional (Log2FC) and non-directional (abs(Log2FC)) versions of the analysis were carried out using the *fgsea* (v.1.32.0) package ^89^ (parameters: nperm=10000, min size=10, max size=1000) and a gene set database including both traditional gene ontology gene sets and custom brain-related gene sets (*Brain.GMT*: ^90^). P-values were corrected for FDR using the Benjamini-Hochberg method.

To explore cell type specificity, gene set enrichment analysis was also used to compare the results of the hippocampal meta-analyses (full, traditional and non-traditional antidepressant meta-analyses) to previously-documented effects of FLX on hippocampal cell types ^59^ or electroconvulsive shock (ECS) on hippocampal neurons ^60^ as measured by single nucleus RNA-Seq (snRNA-Seq). Gene sets were constructed representing all genes that were significantly upregulated or downregulated in response to FLX (5 days of treatment or 3 weeks of treatment) or 10 sessions of ECS (20 days of treatment) in each of the hippocampal cell type clusters (46 gene sets total). Two gene sets were also constructed representing the “developmental-like state” that was shown to increase in response to FLX and ECS, as measured using the top 100 genes upregulated and down-regulated in postnatal day 5 vs. postnatal day 132 DG granule cells ^91^. Enrichment within this custom gene set database was calculated using fGSEA using the procedure described above, but allowing for larger gene sets (max size=2500), followed by FDR correction.

### Comparison with previous FLX meta-analysis (Ibrahim et al. 2022)

The effects of the FLX on the hippocampus ^58^ were recently characterized using meta-analyses of transcriptional profiling results from either stressed (7 studies, n=75) or stress-naive rodents (7 studies, n=56). Due to differing inclusion/exclusion criteria, only 66% of the samples included in their meta-analyses were in our own meta-analysis (n=87 of 131), contributing 28% to our final sample size (n=87 of 313). Therefore, their results can provide insight that is partially independent from our own. Their publication provided results for the top DEGs identified using two methods of integrating findings across studies (integration method and portrait method) and consensus scores indicating whether those DEGs were predominantly upregulated or downregulated for the stressed (*Table 2 & 3 in* ^58^) and stress-naive animals (*Table 6 & Table 7 in* ^58^). For comparison with our results, consensus scores from these tables were converted to a categorical variable indicating “Up” or “Down” direction of effect, which was then tested as a predictor for effects within our meta-analyses (Log2FCs) using standard bivariate linear regression.

**Table 2.**
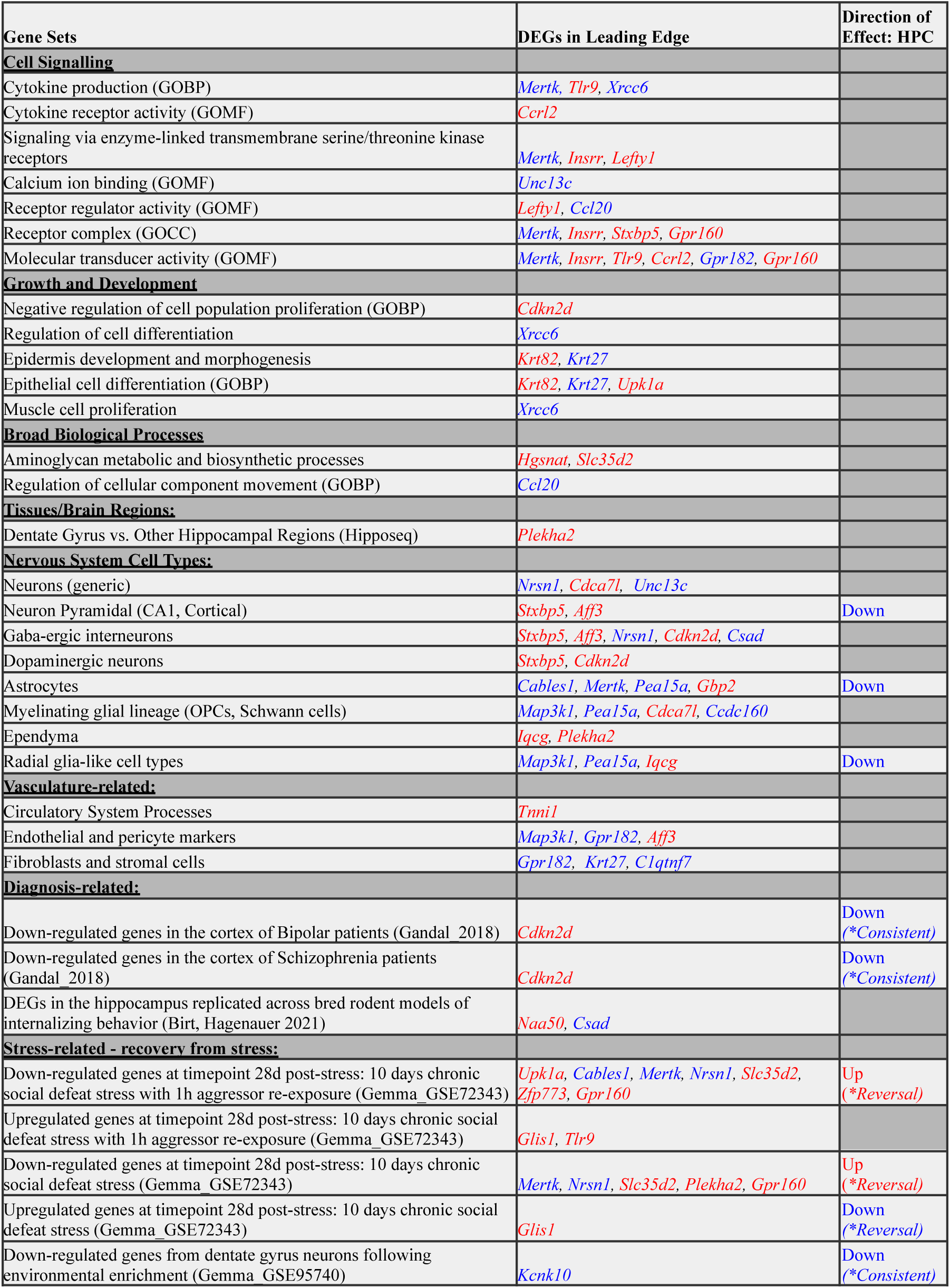
Convergent antidepressant effects on the hippocampus are enriched in many cell types and pathways, some of which have been previously linked to depression. Gene sets were identified as significantly enriched with antidepressant-related differential expression in the hippocampal meta-analysis via non-directional functional gene set enrichment analysis (fGSEA, FDR<0.05), and narrowed down to gene sets containing significant hippocampal antidepressant DEGs (FDR<0.05) amongst the leading genes driving the enrichment. The direction of effect for the gene set is provided if the directional hippocampal fGSEA analysis trended towards significance (FDR<0.10). Gene sets have been organized into groupings based on function, and filtered to reflect brain-relevant functions. Gene set names have been reformatted and simplified for easy viewing, removing highly redundant terms with similar leading genes. Color indicates direction of effect; red: upregulated, blue: downregulated. Full results can be found in **Table S5** and **Table S6**. <u>Abbreviations</u>: DEG: Differentially expressed gene, FDR: False Discovery Rate (q-value), HPC: Hippocampus, fGSEA: fast gene set enrichment analysis, GOBP: Gene Ontology biological process, GOCC: Gene Ontology cellular component, GOMF: Gene Ontology molecular function, CA1: Cornu Ammonus 1, OPC: Oligodendrocyte progenitor cell

## Results

### Hippocampus

#### Characteristics of the datasets included in the hippocampal meta-analysis

Fifty-eight datasets were identified using prespecified antidepressant search terms (**Fig. 1**). Of these, fifteen datasets survived inclusion/exclusion criteria, leaving a final collective sample size of *n*=313 hippocampal samples from antidepressant-treated and control subjects. These datasets included 22 antidepressant vs. control comparisons (*full extracted results:* **Table S1**), representing eleven types of treatments, including pharmacological treatments (SSRIs, TCAs, SNRIs, NMDAR antagonists, Alpha-2 adrenergic antagonists with TCAs, atypical antidepressants, and MASSAs) and non-pharmacological treatments (ECT, TMS) (**Table 1**), providing a diverse sample for examining the converging effects of different antidepressants on hippocampal gene expression.

The antidepressant treatments were tested using a variety of rodent depression models, including chronic unpredictable mild stress (CUMS), lipopolysaccharide-induced depression-like behavior, chronic corticosterone treatment, stress-susceptible strains, and chronic social defeat stress (CSDS) ^69,72,73,76–78,92^. Three datasets included aged subjects, some with transgenically-induced cognitive impairment ^71,80^. Three datasets utilized no depression model ^74,75,79^. For species, most datasets used *Mus musculus*, with six using *Rattus norvegicus*. All samples were male (**Table 1**).

Other potential sources of heterogeneity in the datasets included platform and dissection. Most datasets were collected using microarray technology; the remaining three were by RNA-seq. Two-thirds were derived from whole Ammon’s horn tissue, with the remaining specified as DG (**Table 1**).

#### Meta-analysis reveals consistent differential expression across antidepressant categories that was enriched in many cell types and pathways previously linked to depression

To be included in the meta-analysis, a gene needed to be present in a minimum of eleven antidepressant vs. control comparisons. 16,494 genes fulfilled this requirement, and 16,439 genes produced stable meta-analysis estimates (*full results:* **Table S2**). Fifty-eight genes were significantly differentially expressed (FDR<0.05, “DEGs”, **Table S3, Table S4**), of which 23 were consistently upregulated and 35 consistently downregulated following a variety of antidepressant treatments (*example forest plots:* **Fig. 2**) often in 19 or more comparisons (*heatmap:* **Fig. 3)**.

**Figure 2.**
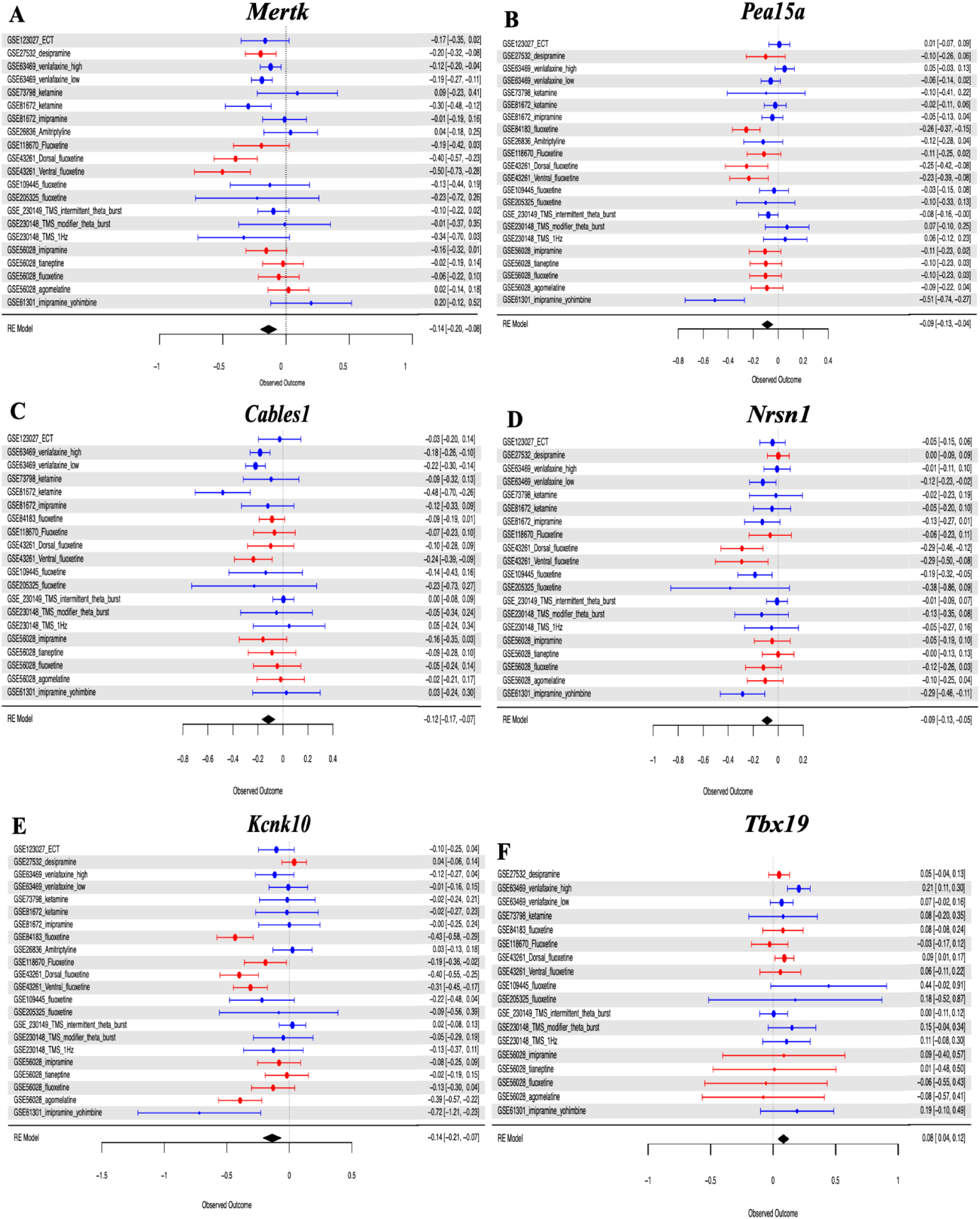
Hippocampal meta-analysis reveals consistent differential expression across antidepressant categories: Example forest plots for six of the top DEGs (FDR<0.05). Forest plots allow for visual inspection of the consistency and magnitude of antidepressant effects across all comparisons. Rows illustrate Log2FC (circles) with 95% confidence intervals (whiskers) for each of the antidepressant vs. control comparisons in each of the datasets and the meta-analysis random effects model (“RE Model”). Color indicates tissue type, with DG in red and Ammon’s horn in blue. The number of treatment vs. control comparisons that included measurements for the gene is determined following quality control. **A.** MER proto-oncogene tyrosine kinase (Mertk) was significantly downregulated across 21 comparisons. **B.** Proliferation and apoptosis adaptor protein 15A (Pea15a) was significantly downregulated across 22 comparisons. **C.** Cdk5 And Abl Enzyme Substrate (Cables1) was significantly downregulated across 20 comparisons. **D.** Neurensin 1 (Nrsn1) was significantly downregulated across 20 comparisons. **E.** Potassium Two Pore Domain Channel Subfamily K Member 10 (Kcnk10) was significantly downregulated across 22 comparisons. **F.** T-box Transcription Factor 19 (Tbx19) was significantly upregulated across 18 comparisons. <u>Abbreviations</u>: DEG: Differentially expressed gene, RE Model: Random effects model, DG: Dentate gyrus, Log2FC: Log2 Fold Change (Antidepressant treatment vs. Control), FDR: False Discovery Rate (q-value).

**Figure 3.**
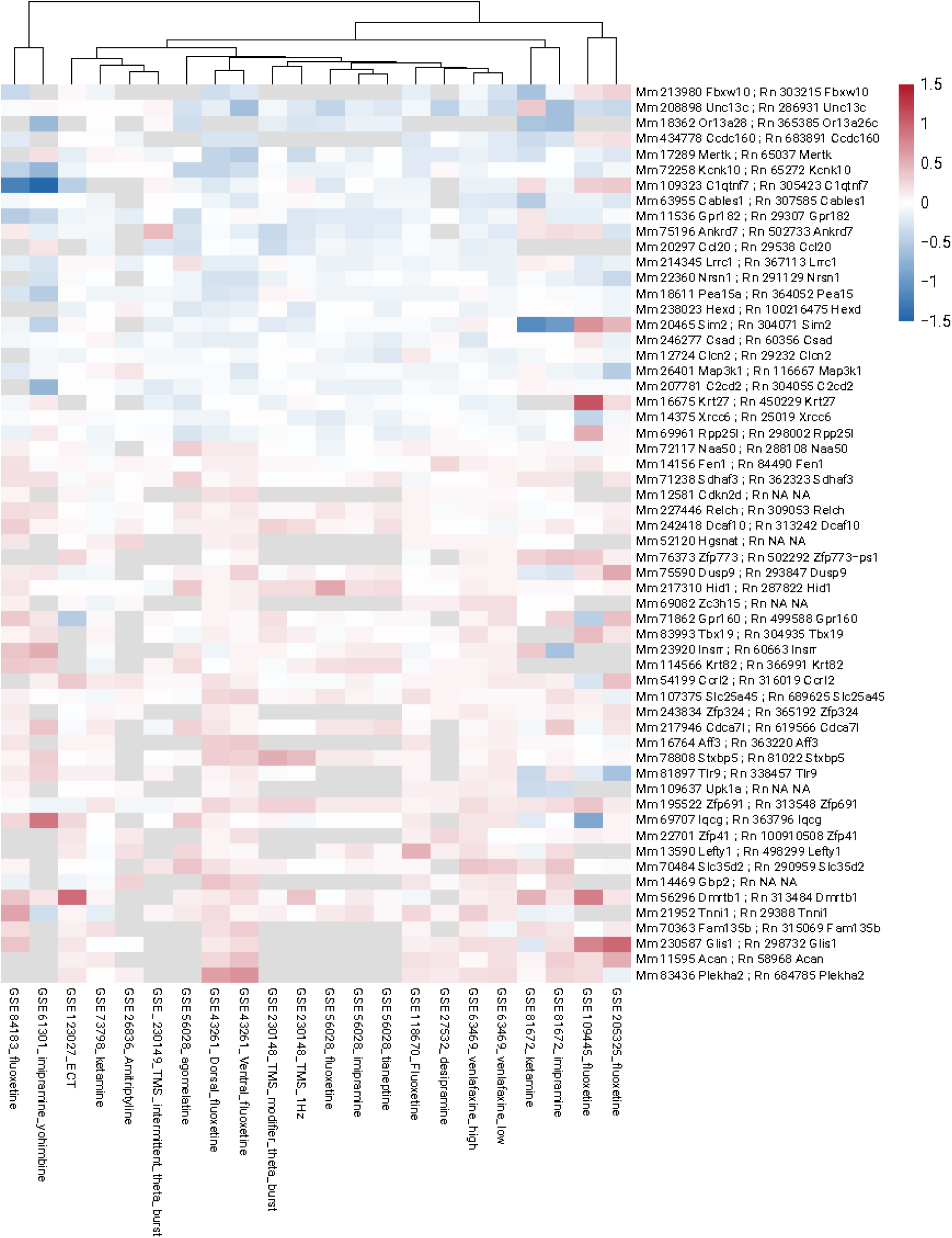
A hierarchically-clustered heatmap of antidepressant effects across hippocampal datasets shows the consistency of effects across treatment type, tissue type, and platform for the top DEGs identified in the meta-analysis. The heatmap allows for visual comparison of the expression patterns of the 58 hippocampal meta-analysis DEGs (FDR<0.05) across individual datasets. The color scale indicates standardized effect size (log₂ fold change), with red denoting upregulation and blue denoting downregulation relative to control samples. Each column represents results from an individual antidepressant treatment (vs. control) comparison from the included datasets, as identified using the Gene Expression Omnibus accession # (GSE…). Each row represents one gene, identified by mouse (Mm) Entrez ID and official mouse gene symbol and rat (Rn) Entrez ID and official rat gene symbol, with NA values indicating a lack of clear ortholog in the species. At the top of the figure the dendrogram groups datasets by similarity in their transcriptional profiles, with shorter connecting lines indicating greater similarity in gene expression changes across datasets. <u>Abbreviations</u>: DEG: Differentially expressed gene, Mm: mouse, Rn: rat, FDR: False Discovery Rate (q-value).

This convergent antidepressant differential expression was enriched within 114 gene sets (non-directional fGSEA analysis FDR<0.05, *full results:* **Table S5**), many of which were associated with factors modulating depression risk. Of these enriched gene sets, 62 contained at least one significant antidepressant DEG amongst the “leading edge” genes driving the enrichment, and thus are of particular interest. The enriched gene sets containing DEGs spanned several categories (summarized in **Table 2**), including cell signalling (*e.g.,* cytokine activity, calcium binding, enzyme-linked kinase receptors), growth and development (*e.g.,* cell proliferation and differentiation, morphogenesis, the DG), central nervous system cell types (*e.g.,* pyramidal, GABA-ergic, and dopaminergic neurons, astrocytes, myelinating cells, ependyma), and vasculature-related cell types (e.g., endothelial cells, pericytes, fibroblasts). There was also enrichment in gene sets derived from previous differential expression studies, including several included in our meta-analysis (2 gene sets, *not shown*), and studies of psychiatric disorders, depression, stress-related animal models, and environmental enrichment. The directional version of the analysis identified 36 gene sets showing a non-significant trend towards enrichment with antidepressant effects (FDR<0.1, *full results:* **Table S6**), twenty-three of which overlapped the non-directional results (**Table 2**). These pathways enriched with convergent antidepressant differential expression are worth investigating further as potential linchpins for antidepressant efficacy.

#### Assessment of Result Robustness and Validity

Overall, the risk of systematic bias within the datasets from the individual transcriptional profiling studies was deemed low for any particular gene-level result, as the original publications conducted full genome analyses and the data was re-analyzed using a standardized pipeline. That said, the bias in the literature against negative results may inflate effect sizes represented in publicly-released datasets, but in a manner that should be impartial to direction-of-effect. To double-check our assumptions, we conducted an Egger’s regression analysis to identify genes that might show evidence of funnel plot asymmetry, a commonly-used metric indicating potential publication bias. We found 58 genes that showed evidence of publication bias (Egger’s test FDR<0.05: **Table S7**), several of which were immediate early genes (*Jun, Egr4*) or genes involved in neurotransmission (*Tdo2, Grm7, Calb1, Npy*). This could suggest that antidepressant datasets were more likely to be published or released if they supported neurotransmission-related hypotheses. However, none of the genes with evidence of publication bias were amongst our antidepressant meta-analysis DEGs (FDR<0.05, *full results:* **Table S2**).

To assess the influence of individual studies and contrasts on meta-analysis results, we repeated the meta-analysis while iteratively excluding each of the study contrasts. We found that none of our meta-analysis DEGs dropped below nominal significance (*p*<0.05) following the removal of single study contrasts (maximum *p*=0.0091), suggesting that our results were robust (*full results:* **Table S2**). We did not find evidence that small sample size studies were adding disproportionate noise in our meta-analyses (**Fig. S1** & **Fig. S2**).

#### Assessment of Sources of Antidepressant Effect Heterogeneity

Our meta-analysis identified converging antidepressant effects across studies and treatments, but there was still evidence of residual heterogeneity. For example, although a hierarchically-clustered heatmap of the top 58 antidepressant DEGs in our meta-analysis did not show any obvious pattern in treatment type, platform or tissue type (**Fig. 3**), 14 of the DEGs still showed indications of significant true variation in antidepressant effects (Log2FC) across studies and contrasts as indicated by Cochran’s Q-test (FDR<0.05). This variation was greater when considering the full 16,494 genes included in the meta-analysis, which had a median I^2^ of >50%, in a manner that was similarly not elucidated using simple hierarchical clustering (**Fig. S1**). Therefore, we conducted a series of exploratory analyses to examine potential sources of effect heterogeneity.

#### Exploratory analyses: Traditional and non-traditional antidepressants have overlapping effects on the hippocampus

As an exploratory follow-up analysis, we compared the patterns of gene expression associated with traditional, mono-aminergic targeting antidepressants and non-traditional antidepressants, both pharmacological and neuromodulatory (**Table 1**, **Fig. 4A**). In the traditional antidepressant meta-analysis, ten datasets were included, with a collective sample size of *n*=177 samples used in thirteen antidepressant vs. control comparisons. In the non-traditional antidepressant meta-analysis, six datasets were included, with a collective sample size of *n*=148 samples used in eight antidepressant vs. control comparisons. For a gene to be included in the traditional subgroup meta-analysis, differential expression results needed to be available from a minimum of 12 antidepressant vs. control comparisons. Of the 9,628 genes fulfilling this requirement, 9,612 produced stable meta-analysis estimates. Sixty-five of these genes were significantly differentially expressed (FDR<0.05: 42 upregulated, 23 downregulated: **Fig. S3**, *full results*: **Table S8**). For the non-traditional subgroup meta-analysis, a gene needed to be present in a minimum of 7 antidepressant vs. control comparisons. Of the 12,203 genes fulfilling this requirement, 12,154 produced stable meta-analysis estimates. None of these genes were significantly differentially expressed (**Fig. S4**, *full results:* **Table S9**), but the pattern of non-traditional antidepressant-related expression moderately resembled that observed in the traditional subgroup meta-analysis when considering all genes present in both analyses (*df*=12,614, *Rho*=0.257, *p*<2.2e-16, **Fig S5**). As would be expected, both traditional and non-traditional antidepressants showed converging antidepressant effects for the top DEGs identified in our original meta-analysis (e.g., FDR<0.10 in original analysis: *df*=61, *Rho*=0.743, *p*<2.2e-16: **Fig. 4B**).

**Figure 4.**
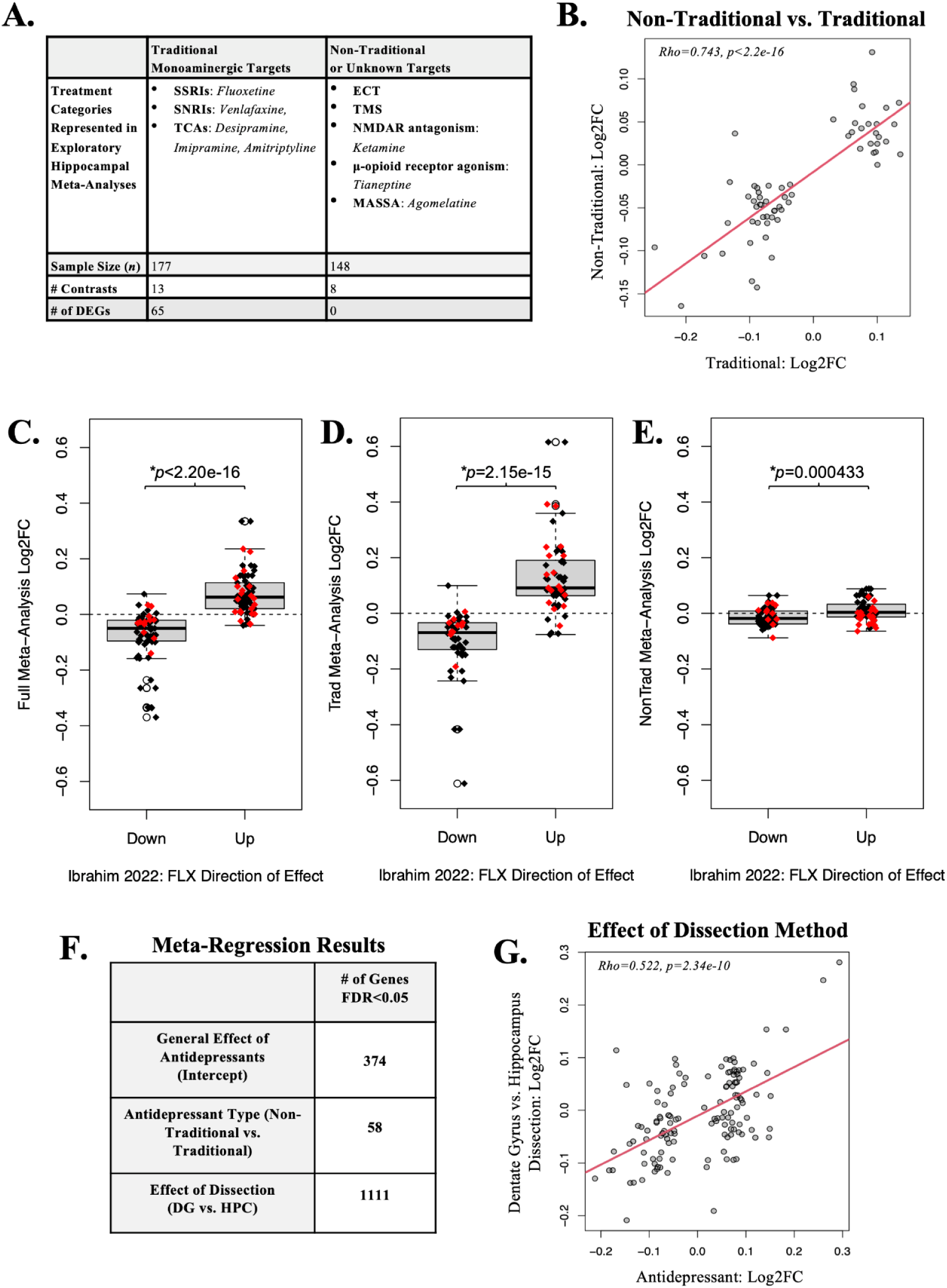
Exploratory analyses: Traditional and non-traditional antidepressants have overlapping effects on the hippocampus. **A.** Table summarizing the types of antidepressant treatments, sample sizes, and number of antidepressant vs. control contrasts included for the exploratory meta-analyses examining the effects of traditional (n=177, 13 contrasts) and non-traditional (n=148, 8 contrasts) antidepressants on the hippocampus. **B.** Scatterplot comparing log₂ fold changes from the traditional and non-traditional antidepressant treated hippocampus shows converging effects for the top antidepressant genes identified in our original meta-analysis (FDR<0.10: Rho=0.743, p<2.2e-16). Convergence is also seen when comparing the effects of traditional and non-traditional antidepressants across all genes present in both analyses (Rho=0.257, p<2.2e-16, **Fig S5**). **C-E.** Boxplots with overlaid jittered data points illustrating that our meta-analyses replicated the direction of effect observed within previous meta-analyses of the effects of fluoxetine (FLX) on the hippocampus ^58^. The previous meta-analyses used a sample that only partially overlapped with our own (28%) and thus provide semi-independent insight. The y-axis shows the Log2FC estimated within our antidepressant meta-analyses (**C:** full meta-analysis, **D:** traditional antidepressant meta-analysis, **E:** non-traditional antidepressant meta-analysis) for each of the top genes identified within the previous FLX meta-analyses of transcriptional profiling results from stressed (red) and stress-naive (black) samples using both reported methods (“portrait” or “integration”). The x-axis indicates the direction of effect identified by the consensus score in the FLX meta-analyses. Boxes = first quartile, median, and third quartile; whiskers = range or 1.5× the interquartile range; open dot = outlier datapoint falling beyond the whiskers of the boxplot. **F.** An exploratory meta-regression controlling for heterogeneity introduced by antidepressant type (non-traditional vs. traditional) and dissection (dentate gyrus (DG) vs. whole hippocampus (HPC)) revealed more genes showing general antidepressant effects (FDR<0.05) than our original hippocampal meta-analysis, as well as many genes showing effects modulated by antidepressant type (FDR<0.10) and dissection (FDR<0.10). **G.** The modulating effect of dissection (Log2FC) identified by the meta-regression correlated with the overall effect of antidepressants (Log2FC) both for the top genes identified in our original meta-analysis (scatterplot: genes with original FDR<0.10: Rho=0.522, p=2.34e-10) and for all genes in the analysis (**Fig. S8**: Rho=0.498, p<2.2e-16) in a manner suggesting that antidepressant effects were larger in DG dissections than in the whole hippocampus. <u>Abbreviations</u>: DEG: Differentially expressed gene, FDR: False Discovery Rate (q-value), Rho: Spearman rank correlation, p: nominal p-value, n: sample size, DG: Dentate gyrus, HPC: Hippocampus, Log2FC: Log(2) Fold Change, FLX: Fluoxetine, SSRI: selective serotonin reuptake inhibitor, NMDA: N-Methyl-D-aspartate, SNRI: serotonin-norepinephrine reuptake inhibitor, TCA: Tricyclic Antidepressant, MASSAs: melatonin agonist and selective serotonin antagonist, ECT: Electroconvulsive therapy, TMS: Transcranial magnetic stimulation, Trad: Traditional mono-aminergic targeting antidepressants, NonTrad: Non-traditional antidepressants

A meta-regression that included antidepressant type (non-traditional vs. traditional) and dissection (DG vs. whole hippocampus) as co-variates showed even stronger findings (**Fig. S6**). The general antidepressant effects detected within the meta-regression with co-variates were very similar to the antidepressant effects detected in our original meta-analysis (**Fig. S7A**: *df*=16,401, *Rho*=0.900, *p*<2.2e-16), and the predicted effects of traditional and non-traditional antidepressants paralleled our previous subsetted meta-analysis findings (**Fig. S7B:** traditional: *df*=9,594, *Rho*=0.954, *p*<2.2e-16, **Fig. S7C:** non-traditional: *df*=12,123, *Rho*=0.779, *p*<2.2e-16). However, more genes now showed a significant overall effect of antidepressants (374 genes *FDR*<0.05) due to better controlling for heterogeneity, with 58 genes showed a modulating effect of antidepressant type (*FDR*<0.05) and 1111 genes showed a modulating effect of dissection (*FDR*<0.05, **Fig 4F**, *full results:* **Table S10**). Notably, the modulating effect of dissection correlated with both our original antidepressant meta-analysis estimates (**Fig. S8A**: all genes: *df*=16,401, *Rho*=0.362, *p*<2.2e-16, original top DEGs (*FDR*<0.10): *df*=128, *Rho*=0.343, *p*=6.98e-05) and current antidepressant meta-regression estimates (**Fig. S8B:** all genes: *df*=16,440, *Rho*=0.498, *p*<2.2e-16, original top DEGs (*FDR*<0.10): *df*=128, *Rho*=0.522, *p*=2.34e-10, **Fig 4G**) in a manner suggesting that antidepressant effects were larger in DG dissections than in the whole hippocampus.

We attempted exploratory meta-regression analyses to examine other potential sources of effect heterogeneity (platform, inclusion of a depression model) but found little evidence of impact, most likely because these variables were not well represented in our design (**Fig S9,** *full results:* **Table S11** & **Table S12**).

#### Exploratory analysis: The pattern of effects detected in previous FLX meta-analyses appear partially generalizable to other antidepressants

The effects of FLX on hippocampal gene expression in stressed and stress-naive animals were recently characterized using meta-analyses of public transcriptional profiling studies ^58^. When comparing our meta-analysis findings to their results, we found reasonable convergence, despite only partially overlapping samples (28%). Overall, we tended to observe the same direction of effect (**Table S13**), with more positive Log2FCs observed in our meta-analyses for genes previously found to be upregulated with FLX vs. down-regulated with FLX (**Fig. 4D**, *full meta-analysis*: *R*^2^=0.438, *β+/-SE*=0.149+/-0.0146, *T*(135)=10.25, *p*<2.2e-16). This similarity was stronger with the results of our meta-analysis using only traditional antidepressants (**Fig. 4E**, *R^2^*=0.447, *β*+/-SE=0.237+/-0.0254, *T*(105)=9.311, *p*<2.15e-15), which had an overlapping sample, but still present - although weaker - in our meta-analysis using non-traditional antidepressants (**Fig. 4F**, *R^2^*=0.0969, *β+/-SE*=0.0227+/-0.00627, *T*(122)=3.618, *p*=0.000433). There was no overlap with our top DEGs, but 22 of the FLX genes showed the same direction of effect and nominal significance (p<0.05) in our full hippocampal meta-analysis, and only one showed the opposite direction of effect and nominal significance, providing suggestive support.

#### Exploratory analysis: Antidepressant effects in the current hippocampal meta-analyses reproduce effects detected in individual cell types using snRNA-seq

To follow up on the enrichment of antidepressant effects within gene sets associated with particular hippocampal cell types, we compared our meta-analysis results to the effects of antidepressant treatments observed in two snRNA-Seq studies: the effects of FLX (5 days or 3 weeks of treatment) on all hippocampal cell types ^59^ and the effects of ECS (10 sessions over 20 days of treatment) on hippocampal neuronal cell types ^60^. We found that differential expression within the antidepressant meta-analyses was enriched within sets of genes that responded to three weeks of FLX and ECS treatment in multiple hippocampal cell type clusters (**Table S14**), including DG granule neurons, astrocytes, and other hippocampal neuron types (CA1, CA3, Mossy, GABA-ergic), and within sets of genes responding to 5 days of FLX treatment in microglia and oligodendrocytes. Furthermore, the enrichment of upregulation or downregulation within the meta-analyses results always paralleled the direction of effect observed in the snRNA-Seq experiments for treatments with demonstrable antidepressant efficacy (3 weeks of FLX and ECS). There was also an enrichment of antidepressant effects within two gene sets previously identified as signifying the re-activation of a “developmental-like state” in DG granule neurons^60,93^, which was theorized to mediate the antidepressant effects of FLX ^59^ and ECS ^60^ (**Fig. 5**).

**Figure 5.**
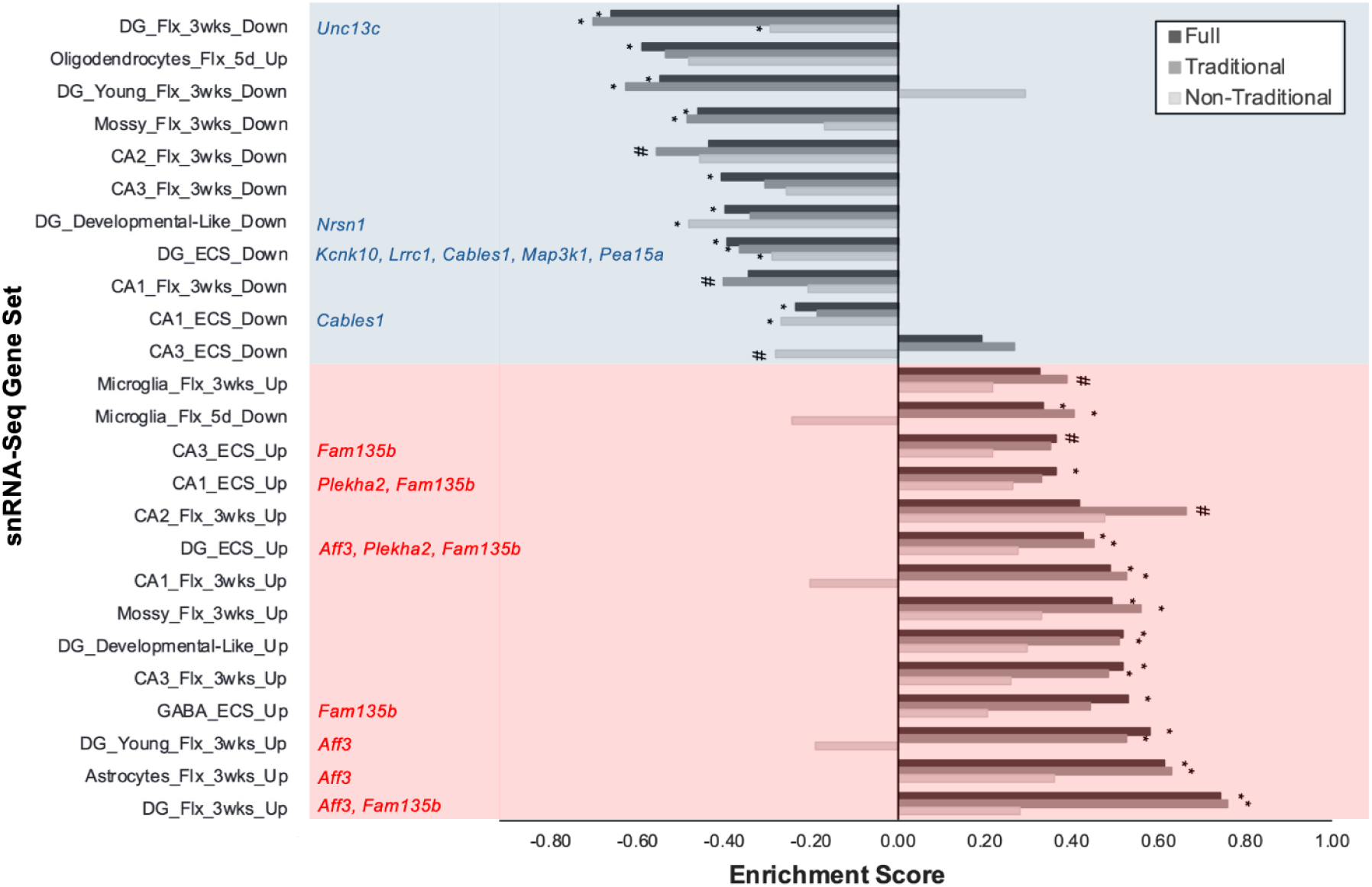
Exploratory analysis: Meta-analyses of antidepressant effects identified in bulk hippocampal tissue uncover similar patterns of differential expression as those previously identified in specific hippocampal cell types using snRNA-Seq. Sets of genes that were found to be differentially expressed in hippocampal cell types in response to fluoxetine (FLX: 5 days (5d) or 3 weeks (3wks) of treatment) or electroconvulsive shock (ECS, 10 days) as measured by snRNA-Seq were enriched with similar differential expression in our hippocampal antidepressant meta-analyses (full meta-analysis, exploratory meta-analysis focused on traditional antidepressants, exploratory meta-analysis focused on non-traditional antidepressants). For ease of visualization, only gene sets with at least nominal enrichment (#p<0.05) of differential expression in at least one of the meta-analyses were included in the figure, with asterisks indicating enrichment surviving false discovery rate correction (*FDR<0.05). In general, the direction of effect for the enrichment in the antidepressant meta-analyses tended to mirror the direction of effect observed in the original snRNA-Seq results. <u>Abbreviations</u>: DG: Dentate Gyrus (granule cells), Mossy: Mossy Cells, CA: Cornu Ammonis (neurons), GABA: GABAergic interneurons, wks: Weeks, d: Days

There was evidence of treatment heterogeneity: in general, the traditional antidepressant meta-analysis results tended to more closely resemble the FLX snRNA-Seq results than the non-traditional meta-analysis results (**Fig. 5, Table S14**). This might be expected due to FLX targeting traditional monoaminergic systems (SSRI). However, a similar bias was not observed for the ECS-derived gene sets, and the full meta-analysis results were strongly enriched with differential expression in gene sets derived from both FLX and ECS snRNA-Seq experiments. Notably, genes that were down-regulated in DG granule neurons following 3 weeks of FLX or ECS showed enrichment within all three of our hippocampal meta-analyses (full, traditional, and non-traditional).

### Cortex

#### Characteristics of the datasets included in the cortical meta-analysis

Using prespecified antidepressant search terms, 147 datasets were identified (**Fig. 1**). Of these datasets, thirteen survived inclusion/exclusion criteria, leaving a final collective sample size of *n*=233 cortical samples from antidepressant-treated and control subjects. These thirteen datasets included 16 antidepressant vs. control comparisons (*full extracted results:* **Table S1**) featuring a variety of antidepressants including pharmacological treatments (SSRIs, SNRIs, TCAs), atypical antipsychotics, NMDA receptor antagonists, serotonin–norepinephrine–dopamine releasing agent (SNDRA) and non-pharmacological treatments (TMS) (**Table 1**), providing a diverse sample for examining the converging effects of different antidepressants on cortical gene expression.

Unlike the hippocampal studies, most cortical studies (representing 10 of 16 contrasts) tested the effects of antidepressants without using depression models ^71,79–88^, but sometimes used samples representing other characteristics including aged transgenic mice ^80^ and inbred strains^81^. The remainder used CSDS and CUMS ^76,78^, a genetic depression model (*Brd1*+/- mice)^83^, and stress-sensitive strain^84^. For species, all but two of the datasets used *Mus musculus*; the remainder used *Rattus norvegicus.* With one exception, the datasets used only males (**Table 1**).

Other potential sources of heterogeneity in the datasets included platform and dissection. Most datasets were collected using microarray technology, with four using RNA-Seq. Most datasets focused on the frontal cortex (11), but dissection varied considerably: four datasets focused on the PFC, one on the prelimbic subregion, and three on the anterior cingulate cortex (ACG). There was also a dataset from the parietal cortex, and “cerebral cortex,” broadly defined (**Table 1**).

#### Cortical meta-analysis reveals one gene consistently upregulated by antidepressants (Atp6v1b2)

To be included in the cortical meta-analysis, a gene needed to be present in a minimum of eleven antidepressant vs. control comparisons. 15,583 genes fulfilled this requirement, and 15,454 genes produced stable meta-analysis estimates (**Fig. 6B**, *full results:* **Table S15**). Only one gene, *ATPase H+ transporting V1 subunit B2* (*Atp6v1b2*) was significantly differentially expressed (*FDR*<0.05), showing upregulation across sixteen comparisons (**Fig. 6D**, *heatmap:* **Fig. S10**). There were no gene sets that were significantly enriched with antidepressant differential expression in the directional or non-directional fGSEA analysis (*FDR*<0.05, *full results:* **Table S16 & Table S17**).

**Figure 6.**
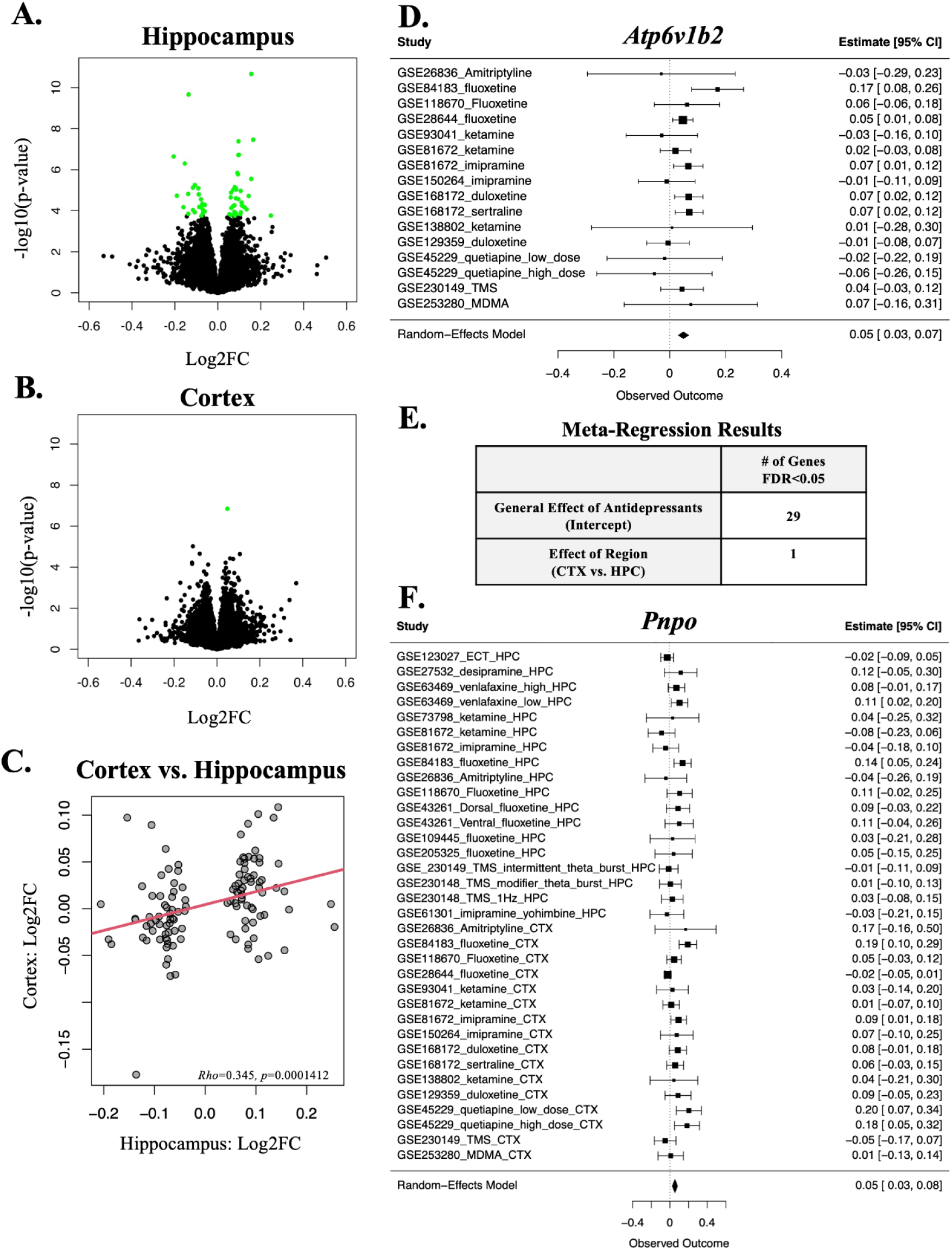
The cortex had fewer significant antidepressant effects, but showed some parallels with the hippocampus. **A–B.** There were more antidepressant DEGs identified in the hippocampus than in the cortical meta-analysis. Volcano plots display differential expression results for hippocampal (A) and cortical (B) meta-analyses, with log₂ fold change plotted against –log₁₀(p-value). Genes reaching significance at FDR<0.05 are highlighted in green. **C.** A scatterplot illustrates the positive correlation between the antidepressant effects (log₂ fold changes or Log2FC) derived from the hippocampal and cortical meta-analyses. This is true when considering either the top-ranking antidepressant genes identified in the hippocampal meta-analysis (depicted above: FDR<0.10 in the original analysis: Rho=0.345, p=0.0001412) or all 14,436 genes included in both analyses (Rho=0.292, p<2e-16, **Fig. S16**). **D.** Forest plot illustrating the antidepressant effects for Atp6v1b2, the one gene that reached significance in the cortical meta-analysis (FDR<0.05), which was upregulated across 16 comparisons. Rows illustrate Log2FC (circles) with 95% confidence intervals (whiskers) for each of the datasets and the meta-analysis random effects model (“RE Model”). **E.** A table showing the results from an exploratory meta-regression using all 38 antidepressant vs. control contrasts derived from both the hippocampus and cortex, including brain region as a co-variate. Out of the 12,857 genes included in the meta-regression, 29 showed an overall effect of antidepressants and 1 showed a modulating effect of brain region. **F.** A forest plot illustrating the small but consistent upregulation of Pyridoxamine 5’-phosphate oxidase (Pnpo) following antidepressant treatment in datasets derived from both the hippocampus (HPC) and cortex (CTX) in the cross-regional meta-regression (FDR<0.05). The enzyme encoded by Pnpo catalyzes the terminal, rate-limiting step in the synthesis of vitamin B6. <u>Abbreviations:</u> HPC: hippocampus, CTX: cortex, Log2FC: Log(2) Fold Change, DEG: Differentially expressed gene, FDR: False discovery rate (q-value), Rho: Spearman rank correlation, p: nominal p-value, RE Model: random-effects model, TMS: Transmagnetic stimulation, MDMA: 3,4-methylenedioxymethamphetamine, ECT: Electroconvulsive therapy

#### Assessment of Result Robustness and Validity

Similar to the hippocampus, the overall risk of systematic bias within the cortical datasets from the individual transcriptional profiling studies was deemed low for any particular gene-level result, as the results were derived from full genome analyses. That said, an Egger’s regression analysis identified 7 genes with evidence of funnel plot asymmetry (FDR<0.05: **Table S18**), providing weak evidence suggesting potential publication bias, but perhaps only in favor of datasets with overall larger effect sizes, as none of the 7 genes had clear theoretical connections to antidepressant mechanisms. *Atp6v1b2* was not included amongst these genes (*full results:* **Table S15**).

To assess the influence of individual studies and contrasts on meta-analysis results, we repeated the meta-analysis while iteratively excluding each of the study contrasts. We found that *Atp6v1b2* did not drop below nominal significance (*p*<0.05) following the removal of single study contrasts (maximum *p*=0.000127), suggesting that the result is robust (*full results:* **Table S15)**. We did not find evidence that small sample size studies were adding disproportionate noise in our meta-analysis (**Fig. S11** & **Fig. S12**).

#### Assessment of Sources of Antidepressant Effect Heterogeneity

The relative lack of converging antidepressant effects across cortical studies and treatments might be due to residual heterogeneity. However, the median I^2^ for the cortical meta-analysis was less than what we observed in the hippocampus (38% vs. 50%, **Fig. S11**), especially when considering the top antidepressant-related genes as ranked by p-value (*e.g.,* the top 58 genes in the hippocampus meta-analysis (∼20%) vs. cortical meta-analysis (0.02%)). Likewise, a hierarchically-clustered heatmap of either the top antidepressant genes (**Fig. S10**) or all 15,583 included in the cortical meta-analysis (**Fig. S11**) showed no obvious pattern related to treatment type, platform or tissue type. We attempted exploratory meta-regression analyses to examine potential sources of effect heterogeneity (antidepressant type, dissection) akin to what was conducted for the hippocampus, but found little evidence of impact (antidepressant type, PFC dissection, **Fig. S13,** *full results:* **Table S19**) or found potential impact but based on minimal observations (ACG dissection, **Fig. S14, Fig. S15,** *full results:* **Table S20**).

#### Antidepressant effects in the cortex show some resemblance to the hippocampus

Despite the relative lack of significant antidepressant DEGs in the cortical meta-analysis, there was a positive correlation between the hippocampal and cortical meta-analysis results when considering either the top antidepressant genes identified in the hippocampal meta-analysis (*FDR*<0.10 in the original analysis: *df*=116, *Rho*=0.345, *p*=0.0001412, **Fig 6C**) or all genes included in both analyses (*df*=14,434, *Rho*=0.290, *p*<2e-16, **Fig S16A**). To explore this similarity further, we conducted a meta-regression using all 38 antidepressant vs. control contrasts derived from both the hippocampus and cortex, including brain region as a co-variate. Out of the 12,857 genes included in the meta-regression, 29 showed an overall effect of antidepressants (*FDR*<0.05, **Fig 6D**, *full results:* **Table S21**), seven of which had been previously identified in our hippocampal meta-analysis. In general, these antidepressant effect sizes tended to be small, with all but 3 of the DEGs showing an *abs*(Log2FC)<0.10 (example forest plot: **Fig 6E**). Only a single gene showed a significant modulating effect of brain region (*FDR*<0.05) (*Sema4d: Semaphorin-4D*), although 20 of the top antidepressant DEGs identified in the previous hippocampal meta-analysis showed a nominal modulating effect of region in the meta-regression (*p*<0.05), suggesting effects were smaller in the cortex. Altogether, these findings imply that broadly-defined brain region (hippocampus vs. cortex) is not a primary source of antidepressant effect heterogeneity in these bulk dissections, whereas hippocampal subregion (DG vs. hippocampus) appears more impactful.

## Discussion

Our meta-analyses provide a view into the shared mechanisms of action of diverse antidepressant treatments in the hippocampus and cortex. A wide variety of treatments were represented, including SSRIs, SNRIs, TCAs, NMDA receptor agonists, alpha-2 adrenergic antagonists combined with TCAs, MASSAs, ECT and TMS. This aggregation of data produced enhanced statistical power, with a collective sample size of *n*=313 (hippocampus) and *n*=233 (cortex). The hippocampal meta-analysis identified 58 DEGs, many of which were associated with gene sets mediating factors known to modulate depression risk, such as stress, environmental enrichment, and other psychiatric disorders, as well as pathways previously linked to antidepressant efficacy, including DG granule neurons, cell proliferation, morphogenesis, vasculature, glia, synaptic plasticity, and immune signaling. Exploratory analyses further highlighted the importance of the DG in antidepressant effects. Our hippocampal meta-analysis also confirmed patterns of gene expression identified in previous antidepressant studies, including a meta-analysis of the effects of the FLX on the hippocampus ^58^ and snRNA-Seq studies characterizing the effects of FLX and ECS on hippocampal cell types ^59,60^, demonstrating that these effects can be generalized to a broad swath of antidepressants.

In the cortex, antidepressant effects resembled the hippocampus in some ways, but there was only one significant DEG, *Atp6v1b2*. When an exploratory meta-regression was performed using both cortical and hippocampal datasets, there were many antidepressant effects that were consistent across both brain regions, with weak evidence of broad regional variation. Altogether, our results indicate that different antidepressant treatments converge on a variety of genes and pathways, with many effects shared across the hippocampus and cortex, but varying at the level of subregion and cell type.

### Hippocampus: Differential expression related to the neurogenic niche in the DG

Neuroimaging studies have consistently shown hippocampal volume reduction in patients with depression, with stress playing a key role ^28,32^, and antidepressants producing a reversal ^32,37,38^. Several theories of antidepressant mechanism center on the promotion of growth-related processes and AHN ^32,33^. Therefore, it is interesting that multiple genes and cell types related to the neurogenic niche in the DG were highlighted in our meta-analysis results, and our exploratory meta-regression suggested that antidepressant effects were larger in DG dissections. Antidepressant treatment upregulated gene expression specific to the DG, including our most upregulated DEG (*Pleckstrin Homology Domain Containing A2 (Plekha2)*). Antidepressant treatment also altered expression specific to a variety of support cell types that play a critical role in the neurogenic niche, including vasculature-related cells (endothelial cells, pericytes, astrocytes) that secrete factors that promote cell proliferation and differentiation ^94–96^ and regulate neural stem cell exposure to blood-borne substances ^96–98^ , and radial glia-like cells, which can serve as quiescent and active neural stem cells ^99,100^.

As our meta-analysis results reflect bulk dissections, these findings could imply functional changes within those cell types or altered cell type balance. To address this question, we conducted an exploratory follow-up analysis comparing our meta-analysis findings to snRNA-Seq results characterizing the effects of FLX treatment ^59^ and ECS ^60^ on hippocampal cell types. Our findings showed a strong enrichment of antidepressant-related expression amongst sets of genes previously shown to be differentially expressed in response to FLX and ECS in DG granule cells. Moreover, our meta-analysis replicated previous results indicating that antidepressants (FLX, ECS) induce a transcriptional remodelling in DG granule neurons resembling a return to a developmental-like state ^59,60^. To do this, antidepressants were theorized to reopen critical developmental periods by removing specific forms of the extracellular matrix (ECM), thereby allowing synaptic growth and plasticity ^59^. Notably, we also observed differential expression of ECM-related genes (e.g., *Aggrecan* (*Acan*) and *Heparan-alpha-glucosaminide N-acetyltransferase* (*Hgsnat*)) previously shown to modulate anxiety and exploratory activity ^101,102^, and an enrichment of antidepressant-related effects in gene sets specific to fibroblasts, which secrete ECM components (collagen, proteoglycans) ^103^. These findings support the utility of triangulating between highly-powered bulk analyses with diverse samples and smaller, but highly-specific, single-cell or spatial analyses.

### Hippocampus: Differential expression related to neurodevelopment and morphogenesis

Many genes and pathways related to morphogenesis and neurodevelopment were also upregulated following antidepressant exposure, both within our hippocampal meta-analyses and previous snRNA-Seq studies ^59,60^. For example, *Plekha2* was the most strongly upregulated DEG in our meta-analysis, and previously found to be down-regulated following chronic stress and upregulated in the DG following antidepressant interventions (FLX and exercise), correlating with increased AHN and dendritic spine density ^104^. *Plekha2* was also upregulated in DG granule neurons following ECS ^60^, and encodes a protein (TAPP2) which promotes cell adhesion to the ECM ^105,106^, suggesting a mechanism by which antidepressants might modulate ECM regulation of “immature-like” and “mature-like” states in DG granule cells. This protein also modulates a signaling axis (PI3K/AKT: ^107–109^) proffered as a new antidepressant target ^110^ because it mediates neurogenic and growth factor (BDNF: ^111^) effects in depression models ^112–115^. For these reasons, *Plekha2* is a promising candidate for follow-up studies.

Several top DEGs have been directly tied to hippocampal morphology. For example, *Proliferation and apoptosis adaptor protein 15A* (*Pea15a*) was down-regulated by antidepressants in our meta-analysis and in DG granule neurons in response to ECS ^60^ and encodes an abundant phosphoprotein modulating apoptosis and cell proliferation signaling ^116,117^ that correlates with ECT-induced hippocampal grey matter volume increases in depressed patients ^118^. *ALF transcription elongation factor 3 (Aff3)* was upregulated in our meta-analysis and in multiple hippocampal cell types (DG neurons, astrocytes) following FLX or ECS treatment ^59,60^. It encodes a component of the transcriptional super elongation complex (AFF3/LAF4) important for the migration of neural progenitor cells ^119^. As loss of *Aff3* produces hippocampal atrophy and enlarged ventricles ^120^, its upregulation could be important for antidepressant reversal of hippocampal atrophy. *Family With Sequence Similarity 135 Member B (Fam135b)* was also upregulated in our meta-analysis and in hippocampal neurons following FLX or ECS treatment ^59,60^, and can promote neurite extension and neuron survival ^121^, suggesting another mechanism for reversing depression-related atrophy.

### Hippocampus: Differential expression related to neuronal signaling and neuroplasticity

Multiple genes responsible for neuronal signaling were downregulated by antidepressants in our meta-analysis and in snRNA-Seq datasets. For example, *Neurensin 1 (Nrsn1)* is a brain-specific gene potentially important for neural plasticity and nerve signal transduction ^122,123^ that is sensitive to chronic stress. *Nrsn1* was part of the enriched gene set signalling the return of DG granule neurons to a “developmental-like” state theorized to be important for antidepressant effects ^59,60^. *Potassium Two Pore Domain Channel Subfamily K Member 10* (*Kcnk10*) was down-regulated in our meta-analysis, in DG granule neurons in response to ECS ^60^, and following environmental enrichment, which has antidepressant-like effects. It encodes potassium channels that can be stimulated by arachidonic acid, which is implicated in the development of depression ^124,125^. *Unc-13 homolog C* (*Unc13c*) was down-regulated in our meta-analysis and in DG granule neurons in response to FLX ^59^ and mediates excitatory neurotransmission ^126^, whereas another component of hippocampal glutamatergic synapses, *Syntaxin Binding Protein 5* (*Stxbp5*) ^127,128^ was upregulated. Collectively, these findings suggest molecular targets that may mediate antidepressant effects on neuroplasticity.

### Hippocampus: Differential expression related to stress responses

Stress and its effects on the hippocampus are well-documented in the pathophysiology of depression, with the hippocampus acting as both a target and regulator of stress ^129^. Several antidepressant DEGs regulate stress responses. For example, *T-box Transcription Factor 19* (*Tbx19*) was upregulated by antidepressants and is an important regulator of the hypothalamus-pituitary (HPA) stress-response axis, correlating with neurotic traits ^130^. Moreover, sets of genes previously shown to respond to chronic stress were enriched with antidepressant differential expression, typically in the opposing direction. Individual antidepressant DEGs also showed stress-opposing effects (*e.g., Plekha2, Uroplakin 1A (Upk1a)*), whereas other DEGs showed antidepressant effects mimicking chronic stress (*e.g., Nrsn1, GLIS family zinc finger 1* (*Glis1*), and corticosteroid-sensitive *Cdk5 And Abl Enzyme Substrate 1* (*Cables1*) ^60,131^ and *MER proto-oncogene tyrosine kinase* (*Mertk*) ^132^). As some of these genes are tied to brain and behavior abnormalities following stress ^132^, these results may represent an activation of natural compensatory mechanisms. Our findings also suggest possible pathways for antidepressants to reverse stress-effects on growth-related processes, as *Cables1* is responsible for cell cycle progression ^131^ and *Glis1* can promote the generation of pluripotent stem cells ^133^.

### Hippocampus: Differential expression related to immune function

Immune dysregulation is a prominent theory in depression ^134^. Environmental stressors produce an increased immune response in the brain, including activation of hippocampal microglia ^135,136^. Several classes of antidepressants, including SSRIs and SNRIs, modulate inflammatory mechanisms and oxidative stress ^137^. We found that antidepressants upregulated many genes involved in immune cell recruitment, such as *C-C motif chemokine receptor-like 2* (*Ccrl2*), *Plekha2* ^109^, and *Guanylate Binding Protein 2 (Gbp2)* ^138^. Many immune-related cell types in the brain can release neurotrophic factors and anti-inflammatory cytokines as well as pro-inflammatory cytokines ^139,140^, but the observed effects didn’t clearly suggest a reduction in inflammation in favor of growth-related processes. For example, *Gbp2* was upregulated by antidepressants, but promotes M1 (pro-inflammatory) microglial polarization ^141^ and stress-sensitive *Mertk* was down-regulated and mediates anti-inflammatory phagocytosis ^132,142,143^. Similarly, the enrichment of antidepressant effects in cytokine-related gene sets could suggest pro-inflammatory or anti-inflammatory effects. *Toll-like receptor 9* (*Tlr9*) was upregulated and can promote both, encouraging immune homeostasis ^144,145^.

### Cortex: Differential expression of a gene related to microglial function and neuroplasticity

In the cortex, *Atp6v1b2* was the only significant DEG, upregulated across a variety of antidepressant treatments. However, it is a strong candidate for mediating antidepressant effects, as it contains a risk allele for depression ^146,147^, and is differentially methylated following antidepressant treatment ^148^. *Atp6v1b2* encodes a component of vacuolar ATPase (V-ATPase), a proton pump that acidifies intracellular organelles ^149^ in a manner important for protein degradation, autophagy, small molecule transport, and receptor-mediated endocytosis ^149,150^. In microglia, *Atp6v1b2* is likely to promote phagocytosis and M2 anti-inflammatory phenotype ^151–153^. This could mediate antidepressant function, as microglial activation in the frontal cortex correlates with depression severity ^92^. V-ATPase also plays an important role in neurons, generating the electrochemical gradient that facilitates neurotransmitter loading into vesicles ^147,154–158^, suggesting antidepressants could increase the efficiency of synaptic transmission. Furthermore, V-ATPase can activate the mTOR pathway ^159^, which may mediate rapid antidepressant effects ^160^. Altogether, these findings suggest that *Atp6v1b2* is a promising candidate for follow-up, along with other members of the V-ATPase family ^161^.

## Limitations and future directions

### Regional specificity

There was a positive correlation between the antidepressant effects identified in the hippocampus and cortex. Despite that overlap, there was a notable difference in the number of identified DEGs. This may indicate higher sensitivity and responsiveness in the hippocampus to antidepressants, perhaps conferred by the sensitivity of the DG and neurogenic niche, but could also reflect differences in statistical power (n=313 vs. n=233), and heterogeneity in the included subjects and tissue dissections. Notably, an exploratory meta-regression combining data from both the hippocampus and cortex found minimal evidence that broadly-defined brain region (hippocampus vs. cortex) was a primary source of antidepressant effect heterogeneity. However, variation in cortical region and cell type balance may still introduce heterogeneity in these bulk dissections. Within our cortical meta-analysis, we included samples from multiple cortical regions, including the PFC and ACG ^41^. These regions are likely to serve different roles in the manifestation of depression symptoms and treatment response ^162,163^. Exploratory analyses suggested that antidepressant effects in the ACG may differ from other cortical dissections, but it was difficult to draw strong conclusions because there were only 3 or fewer studies representing this region in the meta-analysis for any particular gene. Future well-powered studies will be needed to assess regional heterogeneity.

### Relevance to human clinical populations

Future work should also explore the relevance of our findings to more diverse human clinical populations. Controlled rodent studies provide insight into causality, as treatment effects in clinical populations are often confounded by disease severity, but there are important species differences in brain physiology and morphology ^164^. Also, unlike clinical populations, our samples were almost all male, despite well-documented sex differences in the neurobiological effects of depression in pre-clinical and clinical studies ^165^.

Moreover, the scope of our search included datasets not derived from depression models, and antidepressants may affect neural structures and cognition in nondepressed subjects differently than depressed subjects ^166,167^, including in stress-based animal models ^58^. Notably, most cortical datasets used no depression model, which could account for weaker cortical antidepressant effects. To address this issue, we ran a meta-regression that explored whether the inclusion of a depression model in a study (vs. only controls) modulated antidepressant effects. We found little impact, but our meta-analysis project was not well-designed to answer this question, as antidepressant effects were extracted from each study using the full sample and not subsetted by subject characteristics. Furthermore, the depression models represented in the meta-analyses encompassed highly divergent pathways to depressive-like behavior, including chronic stress, inflammation, and olfactory bulbectomy, which may be optimally reversed by different forms of treatment. Therefore, future studies are needed to better delineate differential treatment effects across animal depression models and subject demographics.

## Conclusion

Altogether, our meta-analyses identified 59 genes that were consistently differentially expressed following treatment with a variety of antidepressants, many of which were associated with functions previously implicated in depression. These genes and pathways are worth investigating as potential linchpins for antidepressant efficacy or targets for novel therapies, with triangulation of results across species to infer translatability. For future work, we plan to compare our results with environmental factors that alleviate depression, like environmental enrichment and exercise. Better-powered studies should also follow up on our exploratory analyses examining how categories of antidepressants differ - instead of converge - in their effects, to elucidate treatment resistance and identify predictors of individual response to different treatment categories.

## Supporting information

Supplemental Methods, Results, Small Tables, Figures, and Legends

## Acknowledgements

This work was completed as part of the *Brain Data Alchemy Project* and supported by the Hope for Depression Research Foundation (HDRF: HA, RH), University of Michigan Undergraduate Research Opportunities Program (SE), Grinnell College Center for Careers, Life, and Service (EH & TQD), National Institute on Drug Abuse (NIDA U01 DA043098: HA & SJW), and the Pritzker Neuropsychiatric Disorders Research Foundation (HA & SJW). Funders and sponsors had no active role in the review. In addition, a thank you to all the other members of the Brain Data Alchemy Project 2024 cohort who helped with critical discussions, ideas and feedback: Lakshmi Chennupati, Ashlee Lewis, Anna Drozman, Brandon Hughes, Duy Nguyen and Toni Duan. We would also like to thank Reviewer #2 for their careful, detailed feedback and guidance.

## CRediT statement

EMG - Conceptualization, Methodology, Software, Formal analysis, Investigation, Data Curation, Writing - Original Draft, Writing - Review & Editing, Visualization

MHH: Conceptualization, Methodology, Software, Formal Analysis, Writing - Review & Editing, Visualization, Validation, Funding Acquisition, Supervision, Project administration

EH: Conceptualization, Methodology, Software, Formal analysis, Investigation, Data Curation, Visualization, Writing - Original Draft, Writing - Review & Editing,

SE: Conceptualization, Methodology, Software, Formal analysis, Investigation, Data Curation, Visualization, Writing - Original Draft, Writing - Review & Editing

EIF: Conceptualization, Methodology, Writing - Review & Editing

PTN: Writing - Review & Editing, Data Curation

AS: Writing - Review & Editing, Data Curation

MRB: Conceptualization, Methodology, Writing - Review & Editing

SM: Conceptualization, Methodology, Writing - Review & Editing

SJW: Writing - Review & Editing, Funding acquisition

HA: Writing - Review & Editing, Supervision, Funding acquisition

RH: Conceptualization, Methodology, Writing - Review & Editing, Supervision, Funding acquisition

## Data Availability Statement

This paper centers on secondary data analysis using publicly available datasets (see Table 1 for full list of accession numbers).

*\*All genes are identified using official gene symbols\**

## Abbreviations

ACG: anterior cingulate cortex
AHN: adult hippocampal neurogenesis
API: application programming interface
β: regression coefficient
BDNF: brain-derived neurotrophic factor
CA: cornu ammonis
CA1, CA2, CA3: cornu ammonis regions 1, 2, and 3
CI_lb: confidence interval lower bound
CI_ub: confidence interval upper bound
Cook’s D: Cook’s distance
CSDS: chronic social defeat stress
Ctrl: control
CTX: cortex
CUMS: chronic unpredictable mild stress
*d*: day
DBS: deep brain stimulation
DEG: differentially expressed gene
DFBetas: differences in betas (indicator of influential observations)
DLPFC: dorsolateral prefrontal cortex
*df*: degrees of freedom
DG: dentate gyrus
DNA: deoxyribonucleic acid
ECS: electroconvulsive shock
ECT: electroconvulsive therapy
FDR: false discovery rate
fGSEA: fast gene set enrichment analysis
Fig: Figure
FLX: fluoxetine
GABA: Gamma-Aminobutyric Acid
*H^2^*: estimated residual variance in effect size, as a ratio compared to total variability
HPC: Hippocampus
*I^2^*: estimated residual variance in effect size, as a percent of total variability
Log2FC: log(2) fold change
LPS: lipopolysaccharide (LPS)-induced depression-like model
MAOI: monoamine oxidase inhibitor
MASSA: melatonin agonist and selective serotonin antagonist
MDD: Major Depressive Disorder
MDMA: 3,4-Methylenedioxymethamphetamine
mRNA: messenger RNA
*n*: sample size
NMDAR: N-Methyl-D-aspartate receptor
OBX: olfactory bulbectomy
*p*: p-value
PFC: prefrontal cortex
PRISMA: Preferred Reporting Items for Systematic reviews and Meta-Analyses
*R*: Pearson correlation coefficient
RE Model: random effects model
*Rho*: Spearman correlation coefficient
RNA: ribonucleic acid
RNA-Seq: RNA-Sequencing
ROI: region of interest
*SE*: standard error
SIA: stress-induced analgesia
SNDRA: serotonin–norepinephrine–dopamine releasing agent
SNRI: serotonin-norepinephrine reuptake inhibitor
snRNA-Seq: single nuclei RNA-Sequencing
SSRI: selective serotonin reuptake inhibitor
*T*: T-statistic
Tau^2^: estimated residual variance in effect size
TCA: tricyclic antidepressant
TMS: transcranial magnetic stimulation
*v*: version
*wk*: week

## References

1. American Psychiatric Association. Diagnostic and Statistical Manual of Mental Disorders. (American Psychiatric Association, 2013). doi:10.1176/appi.books.9780890425596.

2. World Health Organization. Depressive Disorder (Depression). https://www.who.int/news-room/fact-sheets/detail/depression (2023).

3. Friedrich, M. J. Depression Is the Leading Cause of Disability Around the World. JAMA 317, 1517 (2017).

4. Kessler, R. C. & Bromet, E. J. The epidemiology of depression across cultures. Annu. Rev. Public Health 34, 119–138 (2013).

5. Cohen, Z. D. & DeRubeis, R. J. Treatment Selection in Depression. Annu. Rev. Clin. Psychol. 14, 209–236 (2018).

6. Uher, R., Tansey, K. E., Dew, T., Maier, W., Mors, O., Hauser, J., Dernovsek, M. Z., Henigsberg, N., Souery, D., Farmer, A. & McGuffin, P. An Inflammatory Biomarker as a Differential Predictor of Outcome of Depression Treatment With Escitalopram and Nortriptyline. Am. J. Psychiatry 171, 1278–1286 (2014).

7. Norman, T. R. & Olver, J. S. Agomelatine for depression: expanding the horizons? Expert Opin. Pharmacother. 20, 647–656 (2019).

8. Gautam, S., Jain, A., Gautam, M., Vahia, V. & Grover, S. Clinical Practice Guidelines for the management of Depression. Indian J. Psychiatry 59, 34 (2017).

9. Harmer, C. J., Duman, R. S. & Cowen, P. J. How do antidepressants work? New perspectives for refining future treatment approaches. Lancet Psychiatry 4, 409–418 (2017).

10. Artigas, F., Nutt, D. J. & Shelton, R. Mechanism of action of antidepressants. Psychopharmacol. Bull. 36 Suppl 2, 123–132 (2002).

11. Zanos, P. & Gould, T. D. Mechanisms of ketamine action as an antidepressant. Mol. Psychiatry 23, 801–811 (2018).

12. Homayoun, H. & Moghaddam, B. NMDA Receptor Hypofunction Produces Opposite Effects on Prefrontal Cortex Interneurons and Pyramidal Neurons. J. Neurosci. 27, 11496–11500 (2007).

13. Carreno, F. R., Donegan, J. J., Boley, A. M., Shah, A., DeGuzman, M., Frazer, A. & Lodge, D. J. Activation of a ventral hippocampus–medial prefrontal cortex pathway is both necessary and sufficient for an antidepressant response to ketamine. Mol. Psychiatry 21, 1298–1308 (2016).

14. Guardiola-Lemaitre, B., De Bodinat, C., Delagrange, P., Millan, M. J., Munoz, C. & Mocaër, E. Agomelatine: mechanism of action and pharmacological profile in relation to antidepressant properties. Br. J. Pharmacol. 171, 3604–3619 (2014).

15. Racagni, G., Riva, M. A., Molteni, R., Musazzi, L., Calabrese, F., Popoli, M. & Tardito, D. Mode of action of agomelatine: Synergy between melatonergic and 5-HT_2C_ receptors. World J. Biol. Psychiatry 12, 574–587 (2011).

16. Samuels, B. A., Nautiyal, K. M., Kruegel, A. C., Levinstein, M. R., Magalong, V. M., Gassaway, M. M., Grinnell, S. G., Han, J., Ansonoff, M. A., Pintar, J. E., Javitch, J. A., Sames, D. & Hen, R. The Behavioral Effects of the Antidepressant Tianeptine Require the Mu-Opioid Receptor. Neuropsychopharmacology 42, 2052–2063 (2017).

17. Han, J., Andreu, V., Langreck, C., Pekarskaya, E. A., Grinnell, S. G., Allain, F., Magalong, V., Pintar, J., Kieffer, B. L., Harris, A. Z., Javitch, J. A., Hen, R. & Nautiyal, K. M. Mu opioid receptors on hippocampal GABAergic interneurons are critical for the antidepressant effects of tianeptine. Neuropsychopharmacology 47, 1387–1397 (2022).

18. Gill, H., Gill, B., Chen-Li, D., El-Halabi, S., Rodrigues, N. B., Cha, D. S., Lipsitz, O., Lee, Y., Rosenblat, J. D., Majeed, A., Mansur, R. B., Nasri, F., Ho, R. & McIntyre, R. S. The emerging role of psilocybin and MDMA in the treatment of mental illness. Expert Rev. Neurother. 20, 1263–1273 (2020).

19. Tran, K. & Argáez, C. Quetiapine for Major Depressive Disorder: A Review of Clinical Effectiveness, Cost-Effectiveness, and Guidelines. (Canadian Agency for Drugs and Technologies in Health, Ottawa (ON), 2020).

20. Invernizzi, R. W. & Garattini, S. Role of presynaptic α2-adrenoceptors in antidepressant action: recent findings from microdialysis studies. Prog. Neuropsychopharmacol. Biol. Psychiatry 28, 819–827 (2004).

21. Smith, C. B. & Hollingsworth, P. J. α2-Adrenoceptors and antidepressant treatments. in IUPHAR 9th International Congress of Pharmacology London 1984 (eds Paton, W., Mitchell, J., Turner, P., Padgham, C. & Ashcroft, E.) 131–136 (Palgrave Macmillan UK, London, 1984). doi:10.1007/978-1-349-17615-1_20.

22. Figee, M., Riva-Posse, P., Choi, K. S., Bederson, L., Mayberg, H. S. & Kopell, B. H. Deep Brain Stimulation for Depression. Neurotherapeutics 19, 1229–1245 (2022).

23. Kisely, S., Li, A., Warren, N. & Siskind, D. A systematic review and meta-analysis of deep brain stimulation for depression. Depress. Anxiety 35, 468–480 (2018).

24. Singh, A. & Kar, S. K. How Electroconvulsive Therapy Works?: Understanding the Neurobiological Mechanisms. Clin. Psychopharmacol. Neurosci. Off. Sci. J. Korean Coll. Neuropsychopharmacol. 15, 210–221 (2017).

25. Deng, Z.-D., Robins, P. L., Regenold, W., Rohde, P., Dannhauer, M. & Lisanby, S. H. How electroconvulsive therapy works in the treatment of depression: is it the seizure, the electricity, or both? Neuropsychopharmacology 49, 150–162 (2024).

26. Downar, J., Siddiqi, S. H., Mitra, A., Williams, N. & Liston, C. Mechanisms of Action of TMS in the Treatment of Depression. in Emerging Neurobiology of Antidepressant Treatments (eds Browning, M., Cowen, P. J. & Sharp, T.) vol. 66 233–277 (Springer International Publishing, Cham, 2024).

27. Sonmez, A. I., Camsari, D. D., Nandakumar, A. L., Voort, J. L. V., Kung, S., Lewis, C. P. & Croarkin, P. E. Accelerated TMS for Depression: A systematic review and meta-analysis. Psychiatry Res. 273, 770–781 (2019).

28. Belleau, E. L., Treadway, M. T. & Pizzagalli, D. A. The Impact of Stress and Major Depressive Disorder on Hippocampal and Medial Prefrontal Cortex Morphology. Biol. Psychiatry 85, 443–453 (2019).

29. Roddy, D. W., Farrell, C., Doolin, K., Roman, E., Tozzi, L., Frodl, T., O’Keane, V. & O’Hanlon, E. The Hippocampus in Depression: More Than the Sum of Its Parts? Advanced Hippocampal Substructure Segmentation in Depression. Biol. Psychiatry 85, 487–497 (2019).

30. Sheline, Y. I. Depression and the Hippocampus: Cause or Effect? Biol. Psychiatry 70, 308–309 (2011).

31. Charney, D. S. & Manji, H. K. Life Stress, Genes, and Depression: Multiple Pathways Lead to Increased Risk and New Opportunities for Intervention. Sci. STKE 2004, (2004).

32. Sahay, A. & Hen, R. Adult hippocampal neurogenesis in depression. Nat. Neurosci. 10, 1110–1115 (2007).

33. Santarelli, L., Saxe, M., Gross, C., Surget, A., Battaglia, F., Dulawa, S., Weisstaub, N., Lee, J., Duman, R., Arancio, O., Belzung, C. & Hen, R. Requirement of Hippocampal Neurogenesis for the Behavioral Effects of Antidepressants. Science 301, 805–809 (2003).

34. Madsen, T. M., Treschow, A., Bengzon, J., Bolwig, T. G., Lindvall, O. & Tingström, A. Increased neurogenesis in a model of electroconvulsive therapy. Biol. Psychiatry 47, 1043–1049 (2000).

35. Loef, D., Tendolkar, I., Van Eijndhoven, P. F. P., Hoozemans, J. J. M., Oudega, M. L., Rozemuller, A. J. M., Lucassen, P. J., Dols, A. & Dijkstra, A. A. Electroconvulsive therapy is associated with increased immunoreactivity of neuroplasticity markers in the hippocampus of depressed patients. Transl. Psychiatry 13, 355 (2023).

36. Duman, R. S., Deyama, S. & Fogaça, M. V. Role of BDNF in the pathophysiology and treatment of depression: Activity-dependent effects distinguish rapid-acting antidepressants. Eur. J. Neurosci. 53, 126–139 (2021).

37. Sapolsky, R. M. Depression, antidepressants, and the shrinking hippocampus. Proc. Natl. Acad. Sci. 98, 12320–12322 (2001).

38. Czéh, B., Michaelis, T., Watanabe, T., Frahm, J., De Biurrun, G., Van Kampen, M., Bartolomucci, A. & Fuchs, E. Stress-induced changes in cerebral metabolites, hippocampal volume, and cell proliferation are prevented by antidepressant treatment with tianeptine. Proc. Natl. Acad. Sci. 98, 12796–12801 (2001).

39. Schmaal, L., Hibar, D. P., Sämann, P. G., Hall, G. B., Baune, B. T., Jahanshad, N., Cheung, J. W., Van Erp, T. G. M., Bos, D., Ikram, M. A., Vernooij, M. W., Niessen, W. J., Tiemeier, H., Hofman, A., Wittfeld, K., Grabe, H. J., Janowitz, D., Bülow, R., Selonke, M., Völzke, H., Grotegerd, D., Dannlowski, U., Arolt, V., Opel, N., Heindel, W., Kugel, H., Hoehn, D., Czisch, M., Couvy-Duchesne, B., Rentería, M. E., Strike, L. T., Wright, M. J., Mills, N. T., De Zubicaray, G. I., McMahon, K. L., Medland, S. E., Martin, N. G., Gillespie, N. A., Goya-Maldonado, R., Gruber, O., Krämer, B., Hatton, S. N., Lagopoulos, J., Hickie, I. B., Frodl, T., Carballedo, A., Frey, E. M., Van Velzen, L. S., Penninx, B. W. J. H., Van Tol, M.-J., Van Der Wee, N. J., Davey, C. G., Harrison, B. J., Mwangi, B., Cao, B., Soares, J. C., Veer, I. M., Walter, H., Schoepf, D., Zurowski, B., Konrad, C., Schramm, E., Normann, C., Schnell, K., Sacchet, M. D., Gotlib, I. H., MacQueen, G. M., Godlewska, B. R., Nickson, T., McIntosh, A. M., Papmeyer, M., Whalley, H. C., Hall, J., Sussmann, J. E., Li, M., Walter, M., Aftanas, L., Brack, I., Bokhan, N. A., Thompson, P. M. & Veltman, D. J. Cortical abnormalities in adults and adolescents with major depression based on brain scans from 20 cohorts worldwide in the ENIGMA Major Depressive Disorder Working Group. Mol. Psychiatry 22, 900–909 (2017).

40. Krishnan, V. & Nestler, E. J. Animal Models of Depression: Molecular Perspectives. in Molecular and Functional Models in Neuropsychiatry (ed. Hagan, J. J.) vol. 7 121–147 (Springer Berlin Heidelberg, Berlin, Heidelberg, 2011).

41. Pizzagalli, D. A. & Roberts, A. C. Prefrontal cortex and depression. Neuropsychopharmacology 47, 225–246 (2022).

42. Rizvi, S. & Khan, A. M. Use of Transcranial Magnetic Stimulation for Depression. Cureus 10.7759/cureus.4736 (2019) doi:10.7759/cureus.4736.

43. Bellani, M., Dusi, N., Yeh, P.-H., Soares, J. C. & Brambilla, P. The effects of antidepressants on human brain as detected by imaging studies. Focus on major depression. Prog. Neuropsychopharmacol. Biol. Psychiatry 35, 1544–1552 (2011).

44. Hagenauer, M., Manh Nguyen, D., Flandreau, E., Geoghegan, E., Mensch, S., Ra’eed Bhuiyan, M., Thirupatamma Chennupati, L., Espinoza, S., Lewis, A., Drozman, A. & W. Hughes, M.S., Ph.D., B. Brain Data Alchemy Project: Meta-Analysis of Re-Analyzed Public Transcriptional Profiling Data in the Gemma Database (v.2024) v1. Preprint at 10.17504/protocols.io.yxmvmejkng3p/v1 (2024).

45. Rhoads, C. A., Hagenauer, M. H., Xiong, J., Hernandez, E., Nguyen, D. M., Saffron, A., Flandreau, E. I., Watson, S. & Akil, H. A Meta-Analysis of the Effects of Acute Sleep Deprivation on the Cortical Transcriptome in Rodent Models. Preprint at 10.1101/2025.04.21.648791 (2025).

46. R Core Team. _R: A Language and Environment for Statistical Computing_. R Foundation for Statistical Computing (2024).

47. RStudio Team. RStudio: Integrated Development for R. PBC (2024).

48. Lim, N., Tesar, S., Belmadani, M., Poirier-Morency, G., Mancarci, B. O., Sicherman, J., Jacobson, M., Leong, J., Tan, P. & Pavlidis, P. Curation of over 10 000 transcriptomic studies to enable data reuse. Database 2021, baab006 (2021).

49. University of British Columbia. Gemma. (2024).

50. Wickham, H., Averick, M., Bryan, J., Chang, W., McGowan, L., François, R., Grolemund, G., Hayes, A., Henry, L., Hester, J., Kuhn, M., Pedersen, T., Miller, E., Bache, S., Müller, K., Ooms, J., Robinson, D., Seidel, D., Spinu, V., Takahashi, K., Vaughan, D., Wilke, C., Woo, K. & Yutani, H. Welcome to the Tidyverse. J. Open Source Softw. 4, 1686 (2019).

51. Bains, N. & Abdijadid, S. Major Depressive Disorder StatPearls [Internet]. (StatPearls Publishing, Treasure Island (FL)).

52. Gassaway, M. M., Rives, M.-L., Kruegel, A. C., Javitch, J. A. & Sames, D. The atypical antidepressant and neurorestorative agent tianeptine is a μ-opioid receptor agonist. Transl. Psychiatry 4, e411–e411 (2014).

53. Bahji, A., Vazquez, G. H. & Zarate, C. A. Comparative efficacy of racemic ketamine and esketamine for depression: A systematic review and meta-analysis. J. Affect. Disord. 278, 542–555 (2021).

54. Pagnin, D., De Queiroz, V., Pini, S. & Cassano, G. B. Efficacy of ECT in Depression: A Meta-Analytic Review. Focus 6, 155–162 (2008).

55. Patel, R. & Titheradge, D. MDMA for the treatment of mood disorder: all talk no substance? Ther. Adv. Psychopharmacol. 5, 179–188 (2015).

56. dplyr: A Grammar of Data Manipulation. R package version 1.1.4, https://github.com/tidyverse/dplyr, https://dplyr.tidyverse.org. (2023).

57. Wickham, H. plyr: Tools for Splitting, Applying and Combining Data. (2023).

58. Ibrahim, E. C., Gorgievski, V., Ortiz-Teba, P., Belzeaux, R., Turecki, G., Sibille, E., Charbonnier, G. & Tzavara, E. T. Transcriptomic Studies of Antidepressant Action in Rodent Models of Depression: A First Meta-Analysis. Int. J. Mol. Sci. 23, 13543 (2022).

59. Nguyen, P. T., Tamura, S., Sun, E., Shi, Y., Xiao, Y., Lacefield, C., Turi, G. & Hen, R. Antidepressants promote developmental-like plasticity through remodeling of extracellular matrix. Preprint at 10.1101/2025.01.03.631260 (2025).

60. Santiago, A. N., Castello-Saval, J., Nguyen, P., Chung, H. M., Khan, S., Luna, V. M., Hen, R. & Chang, W.-L. Effects of electroconvulsive shock on the function, circuitry, and transcriptome of dentate gyrus granule neurons. Neuropsychopharmacology https://doi.org/10.1038/s41386-026-02345-x (2026) 10.1038/s41386-026-02345-x.

61. Ritchie, M. E., Phipson, B., Wu, D., Hu, Y., Law, C. W., Shi, W. & Smyth, G. K. limma powers differential expression analyses for RNA-sequencing and microarray studies. Nucleic Acids Res. 43, e47 (2015).

62. Baldarelli, R. M., Smith, C. L., Ringwald, M., Richardson, J. E., Bult, C. J., Mouse Genome Informatics Group, Anagnostopoulos, A., Begley, D. A., Bello, S. M., Christie, K., Finger, J. H., Hale, P., Hayamizu, T. F., Hill, D. P., Knowlton, M. N., Krupke, D. M., McAndrews, M., Law, M., McCright, I. J., Ni, L., Onda, H., Sitnikov, D., Smith, C. M., Tomczuk, M., Wilming, L., Xu, J., Zhu, Y., Blodgett, O., Campbell, J. W., Corbani, L. E., Frost, P., Giannatto, S. C., Miers, D. B., Motenko, H., Neuhauser, S. B., Shaw, D. R., Butler, N. E. & Ormsby, J. E. Mouse Genome Informatics: an integrated knowledgebase system for the laboratory mouse. GENETICS 227, iyae031 (2024).

63. Viechtbauer, W. Conducting Meta-Analyses in *R* with the metafor Package. J. Stat. Softw. 36, (2010).

64. Pollard, K. S., Dudoit, S. & Laan, M. J. van der. Multiple Testing Procedures: the multtest Package and Applications to Genomics. in Bioinformatics and Computational Biology Solutions Using R and Bioconductor 249–271 (Springer, New York, NY, 2005). doi:10.1007/0-387-29362-0_15.

65. Egger, M., Davey Smith, G., Schneider, M. & Minder, C. Bias in meta-analysis detected by a simple, graphical test. BMJ 315, 629–634 (1997).

66. Cook, R. D. Detection of Influential Observation in Linear Regression. Technometrics 19, 15–18 (1977).

67. Cochran, W. G. The Combination of Estimates from Different Experiments. Biometrics 10, 101–129 (1954).

68. Wang, Q.-S., Yan, K., Li, K.-D., Gao, L.-N., Wang, X., Liu, H., Zhang, Z., Li, K. & Cui, Y.-L. Targeting hippocampal phospholipid and tryptophan metabolism for antidepressant-like effects of albiflorin. Phytomedicine Int. J. Phytother. Phytopharm. 92, 153735 (2021).

69. Rimmerman, N., Verdiger, H., Goldenberg, H., Naggan, L., Robinson, E., Kozela, E., Gelb, S., Reshef, R., Ryan, K. M., Ayoun, L., Refaeli, R., Ashkenazi, E., Schottlender, N., Ben Hemo-Cohen, L., Pienica, C., Aharonian, M., Dinur, E., Lazar, K., McLoughlin, D. M., Zvi, A. B. & Yirmiya, R. Microglia and their LAG3 checkpoint underlie the antidepressant and neurogenesis-enhancing effects of electroconvulsive stimulation. Mol. Psychiatry 27, 1120–1135 (2022).

70. Demin, K. A., Krotova, N. A., Ilyin, N. P., Galstyan, D. S., Kolesnikova, T. O., Strekalova, T., de Abreu, M. S., Petersen, E. V., Zabegalov, K. N. & Kalueff, A. V. Evolutionarily conserved gene expression patterns for affective disorders revealed using cross-species brain transcriptomic analyses in humans, rats and zebrafish. Sci. Rep. 12, 20836 (2022).

71. Weiler, M., Stieger, K. C., Shroff, K., Klein, J. P., Wood, W. H., Zhang, Y., Chandrasekaran, P., Lehrmann, E., Camandola, S., Long, J. M., Mattson, M. P., Becker, K. G. & Rapp, P. R. Transcriptional changes in the rat brain induced by repetitive transcranial magnetic stimulation. Front. Hum. Neurosci. 17, 1215291 (2023).

72. Lisowski, P., Juszczak, G. R., Goscik, J., Stankiewicz, A. M., Wieczorek, M., Zwierzchowski, L. & Swiergiel, A. H. Stress susceptibility-specific phenotype associated with different hippocampal transcriptomic responses to chronic tricyclic antidepressant treatment in mice. BMC Neurosci. 14, 144 (2013).

73. Patrício, P., Mateus-Pinheiro, A., Irmler, M., Alves, N. D., Machado-Santos, A. R., Morais, M., Correia, J. S., Korostynski, M., Piechota, M., Stoffel, R., Beckers, J., Bessa, J. M., Almeida, O. F., Sousa, N. & Pinto, L. Differential and Converging Molecular Mechanisms of Antidepressants’ Action in the Hippocampal Dentate Gyrus. Neuropsychopharmacology 40, 338–349 (2015).

74. Husain, B. F. A., Nanavaty, I. N., Marathe, S. V., Rajendran, R. & Vaidya, V. A. Hippocampal transcriptional and neurogenic changes evoked by combination yohimbine and imipramine treatment. Prog. Neuropsychopharmacol. Biol. Psychiatry 61, 1–9 (2015).

75. Ficek, J., Zygmunt, M., Piechota, M., Hoinkis, D., Rodriguez Parkitna, J., Przewlocki, R. & Korostynski, M. Molecular profile of dissociative drug ketamine in relation to its rapid antidepressant action. BMC Genomics 17, 362 (2016).

76. Bagot, R. C., Cates, H. M., Purushothaman, I., Vialou, V., Heller, E. A., Yieh, L., LaBonté, B., Peña, C. J., Shen, L., Wittenberg, G. M. & Nestler, E. J. Ketamine and Imipramine Reverse Transcriptional Signatures of Susceptibility and Induce Resilience-Specific Gene Expression Profiles. Biol. Psychiatry 81, 285–295 (2017).

77. Samuels, B. A., Leonardo, E. D., Dranovsky, A., Williams, A., Wong, E., Nesbitt, A. M. I., McCurdy, R. D., Hen, R. & Alter, M. Global State Measures of the Dentate Gyrus Gene Expression System Predict Antidepressant-Sensitive Behaviors. PLoS ONE 9, e85136 (2014).

78. Hervé, M., Bergon, A., Le Guisquet, A.-M., Leman, S., Consoloni, J.-L., Fernandez-Nunez, N., Lefebvre, M.-N., El-Hage, W., Belzeaux, R., Belzung, C. & Ibrahim, E. C. Translational Identification of Transcriptional Signatures of Major Depression and Antidepressant Response. Front. Mol. Neurosci. 10, 248 (2017).

79. Hagihara, H., Ohira, K. & Miyakawa, T. Transcriptomic evidence for immaturity induced by antidepressant fluoxetine in the hippocampus and prefrontal cortex. Neuropsychopharmacol. Rep. 39, 78–89 (2019).

80. Chadwick, W., Mitchell, N., Caroll, J., Zhou, Y., Park, S.-S., Wang, L., Becker, K. G., Zhang, Y., Lehrmann, E., Wood, W. H., Martin, B. & Maudsley, S. Amitriptyline-Mediated Cognitive Enhancement in Aged 3×Tg Alzheimer’s Disease Mice Is Associated with Neurogenesis and Neurotrophic Activity. PLoS ONE 6, e21660 (2011).

81. Benton, C. S., Miller, B. H., Skwerer, S., Suzuki, O., Schultz, L. E., Cameron, M. D., Marron, J. S., Pletcher, M. T. & Wiltshire, T. Evaluating genetic markers and neurobiochemical analytes for fluoxetine response using a panel of mouse inbred strains. Psychopharmacology (Berl.) 221, 297–315 (2012).

82. Orozco-Solis, R., Montellier, E., Aguilar-Arnal, L., Sato, S., Vawter, M. P., Bunney, B. G., Bunney, W. E. & Sassone-Corsi, P. A Circadian Genomic Signature Common to Ketamine and Sleep Deprivation in the Anterior Cingulate Cortex. Biol. Psychiatry 82, 351–360 (2017).

83. Rajkumar, A. P., Qvist, P., Donskov, J. G., Lazarus, R., Pallesen, J., Nava, N., Winther, G., Liebenberg, N., Cour, S. H. L., Paternoster, V., Fryland, T., Palmfeldt, J., Fejgin, K., Mørk, A., Nyegaard, M., Pakkenberg, B., Didriksen, M., Nyengaard, J. R., Wegener, G., Mors, O., Christensen, J. H. & Børglum, A. D. Reduced Brd1 expression leads to reversible depression-like behaviors and gene-expression changes in female mice. Transl. Psychiatry 10, 239 (2020).

84. Funayama, Y., Li, H., Ishimori, E., Kawatake-Kuno, A., Inaba, H., Yamagata, H., Seki, T., Nakagawa, S., Watanabe, Y., Murai, T., Oishi, N. & Uchida, S. Antidepressant Response and Stress Resilience Are Promoted by CART Peptides in GABAergic Neurons of the Anterior Cingulate Cortex. Biol. Psychiatry Glob. Open Sci. 3, 87–98 (2023).

85. Kokkinou, M., Irvine, E. E., Bonsall, D. R., Natesan, S., Wells, L. A., Smith, M., Glegola, J., Paul, E. J., Tossell, K., Veronese, M., Khadayate, S., Dedic, N., Hopkins, S. C., Ungless, M. A., Withers, D. J. & Howes, O. D. Reproducing the dopamine pathophysiology of schizophrenia and approaches to ameliorate it: a translational imaging study with ketamine. Mol. Psychiatry 26, 2562–2576 (2021).

86. Lukić, I., Getselter, D., Ziv, O., Oron, O., Reuveni, E., Koren, O. & Elliott, E. Antidepressants affect gut microbiota and Ruminococcus flavefaciens is able to abolish their effects on depressive-like behavior. Transl. Psychiatry 9, 133 (2019).

87. Kondo, M. A., Tajinda, K., Colantuoni, C., Hiyama, H., Seshadri, S., Huang, B., Pou, S., Furukori, K., Hookway, C., Jaaro-Peled, H., Kano, S., Matsuoka, N., Harada, K., Ni, K., Pevsner, J. & Sawa, A. Unique pharmacological actions of atypical neuroleptic quetiapine: possible role in cell cycle/fate control. Transl. Psychiatry 3, e243–e243 (2013).

88. Warner-Schmidt, J., Stogniew, M., Mandell, B., Rowland, R. S., Schmidt, E. F. & Kelmendi, B. Methylone is a rapid-acting neuroplastogen with less off-target activity than MDMA. Front. Neurosci. 18, 1353131 (2024).

89. Korotkevich, G., Sukhov, V., Budin, N., Shpak, B., Artyomov, M. N. & Sergushichev, A. Fast gene set enrichment analysis. Preprint at 10.1101/060012 (2016).

90. Megan H. Hagenauer, Yusra Sannah, Elaine K. Hebda-Bauer, Cosette Rhoads, Angela M. O’Connor, Stanley J. Watson Jr., & Huda Akil. Resource: A Curated Database of Brain-Related Functional Gene Sets (Brain.GMT). 10.1101/2024.04.05.588301.

91. Hochgerner, H., Zeisel, A., Lönnerberg, P. & Linnarsson, S. Conserved properties of dentate gyrus neurogenesis across postnatal development revealed by single-cell RNA sequencing. Nat. Neurosci. 21, 290–299 (2018).

92. Wang, H., He, Y., Sun, Z., Ren, S., Liu, M., Wang, G. & Yang, J. Microglia in depression: an overview of microglia in the pathogenesis and treatment of depression. J. Neuroinflammation 19, 132 (2022).

93. Nguyen, P. T., Sun, E., Shi, Y., Tamura, S., Xiao, Y., Lacefield, C., Turi, G. F. & Hen, R. Antidepressants reactivate developmental plasticity through remodeling of extracellular matrix. bioRxiv 2025.01.03.631260 (2025) doi:10.1101/2025.01.03.631260.

94. Ehret, F., Vogler, S. & Kempermann, G. A co-culture model of the hippocampal neurogenic niche reveals differential effects of astrocytes, endothelial cells and pericytes on proliferation and differentiation of adult murine precursor cells. Stem Cell Res. 15, 514–521 (2015).

95. Palmer, T. D., Willhoite, A. R. & Gage, F. H. Vascular niche for adult hippocampal neurogenesis. J. Comp. Neurol. 425, 479–494 (2000).

96. Chen, L., Li, Z., Wang, W., Zhou, Y., Li, W. & Wang, Y. Adult hippocampal neurogenesis: New avenues for treatment of brain disorders. Stem Cell Rep. 20, 102600 (2025).

97. Karakatsani, A., Álvarez-Vergara, M. I. & Ruiz De Almodóvar, C. The vasculature of neurogenic niches: Properties and function. Cells Dev. 174, 203841 (2023).

98. Licht, T. & Keshet, E. The vascular niche in adult neurogenesis. Mech. Dev. 138, 56–62 (2015).

99. Llorente, V., Velarde, P., Desco, M. & Gómez-Gaviro, M. V. Current Understanding of the Neural Stem Cell Niches. Cells 11, 3002 (2022).

100. Miranda-Negrón, Y. & García-Arrarás, J. E. Radial glia and radial glia-like cells: Their role in neurogenesis and regeneration. Front. Neurosci. 16, 1006037 (2022).

101. Martins, C., Hůlková, H., Dridi, L., Dormoy-Raclet, V., Grigoryeva, L., Choi, Y., Langford-Smith, A., Wilkinson, F. L., Ohmi, K., DiCristo, G., Hamel, E., Ausseil, J., Cheillan, D., Moreau, A., Svobodová, E., Hájková, Z., Tesařová, M., Hansíková, H., Bigger, B. W., Hrebícek, M. & Pshezhetsky, A. V. Neuroinflammation, mitochondrial defects and neurodegeneration in mucopolysaccharidosis III type C mouse model. Brain J. Neurol. 138, 336–355 (2015).

102. Marcó, S., Pujol, A., Roca, C., Motas, S., Ribera, A., Garcia, M., Molas, M., Villacampa, P., Melia, C. S., Sánchez, V., Sánchez, X., Bertolin, J., Ruberte, J., Haurigot, V. & Bosch, F. Progressive neurologic and somatic disease in a novel mouse model of human mucopolysaccharidosis type IIIC. Dis. Model. Mech. 9, 999–1013 (2016).

103. Dick, M. K., Miao, J. H. & Limaiem, F. Histology, Fibroblast. in StatPearls (StatPearls Publishing, Treasure Island (FL), 2025).

104. Huang, G.-J., Ben-David, E., Tort Piella, A., Edwards, A., Flint, J. & Shifman, S. Neurogenomic Evidence for a Shared Mechanism of the Antidepressant Effects of Exercise and Chronic Fluoxetine in Mice. PLoS ONE 7, e35901 (2012).

105. Rouillard, A. D., Gundersen, G. W., Fernandez, N. F., Wang, Z., Monteiro, C. D., McDermott, M. G. & Ma’ayan, A. The harmonizome: a collection of processed datasets gathered to serve and mine knowledge about genes and proteins. Database 2016, baw100 (2016).

106. Diamant, I., Clarke, D. J. B., Evangelista, J. E., Lingam, N. & Ma’ayan, A. Harmonizome 3.0: integrated knowledge about genes and proteins from diverse multi-omics resources. Nucleic Acids Res. 53, D1016–D1028 (2025).

107. Li, H. & Marshall, A. J. Phosphatidylinositol (3,4) bisphosphate-specific phosphatases and effector proteins: A distinct branch of PI3K signaling. Cell. Signal. 27, 1789–1798 (2015).

108. Wullschleger, S., Wasserman, D. H., Gray, A., Sakamoto, K. & Alessi, D. R. Role of TAPP1 and TAPP2 adaptor binding to PtdIns(3,4)P2 in regulating insulin sensitivity defined by knock-in analysis. Biochem. J. 434, 265–274 (2011).

109. Landego, I., Jayachandran, N., Wullschleger, S., Zhang, T., Gibson, I. W., Miller, A., Alessi, D. R. & Marshall, A. J. Interaction of TAPP adapter proteins with phosphatidylinositol (3,4)-bisphosphate regulates B-cell activation and autoantibody production. Eur. J. Immunol. 42, 2760–2770 (2012).

110. Chen, Y., Guan, W., Wang, M.-L. & Lin, X.-Y. PI3K-AKT/mTOR Signaling in Psychiatric Disorders: A Valuable Target to Stimulate or Suppress? Int. J. Neuropsychopharmacol. 27, pyae010 (2024).

111. Yang, T., Nie, Z., Shu, H., Kuang, Y., Chen, X., Cheng, J., Yu, S. & Liu, H. The Role of BDNF on Neural Plasticity in Depression. Front. Cell. Neurosci. 14, 82 (2020).

112. Wu, H., Lu, D., Jiang, H., Xiong, Y., Qu, C., Li, B., Mahmood, A., Zhou, D. & Chopp, M. Simvastatin-Mediated Upregulation of VEGF and BDNF, Activation of the PI3K/Akt Pathway, and Increase of Neurogenesis Are Associated with Therapeutic Improvement after Traumatic Brain Injury. J. Neurotrauma 25, 130–139 (2008).

113. Jiang, W., Tang, Y.-Y., Zhu, W.-W., Li, C., Zhang, P., Li, R.-Q., Chen, Y.-J., Zou, W. & Tang, X.-Q. PI3K/AKT pathway mediates the antidepressant- and anxiolytic-like roles of hydrogen sulfide in streptozotocin-induced diabetic rats via promoting hippocampal neurogenesis. NeuroToxicology 85, 201–208 (2021).

114. Zheng, R., Zhang, Z.-H., Chen, C., Chen, Y., Jia, S.-Z., Liu, Q., Ni, J.-Z. & Song, G.-L. Selenomethionine promoted hippocampal neurogenesis via the PI3K-Akt-GSK3β-Wnt pathway in a mouse model of Alzheimer’s disease. Biochem. Biophys. Res. Commun. 485, 6–15 (2017).

115. Bruel-Jungerman, E., Veyrac, A., Dufour, F., Horwood, J., Laroche, S. & Davis, S. Inhibition of PI3K-Akt Signaling Blocks Exercise-Mediated Enhancement of Adult Neurogenesis and Synaptic Plasticity in the Dentate Gyrus. PLoS ONE 4, e7901 (2009).

116. Park, J.-Y. & Kang, T.-C. The differential roles of PEA15 phosphorylations in reactive astrogliosis and astroglial apoptosis following status epilepticus. Neurosci. Res. 137, 11–22 (2018).

117. Araujo, H., Danziger, N., Cordier, J., Glowinski, J. & Chneiweiss, H. Characterization of PEA-15, a major substrate for protein kinase C in astrocytes. J. Biol. Chem. 268, 5911–5920 (1993).

118. Sun, H., Bai, T., Zhang, X., Fan, X., Zhang, K., Zhang, J., Hu, Q., Xu, J., Tian, Y. & Wang, K. Molecular mechanisms underlying structural plasticity of electroconvulsive therapy in major depressive disorder. Brain Imaging Behav. 18, 930–941 (2024).

119. Moore, J. M., Oliver, P. L., Finelli, M. J., Lee, S., Lickiss, T., Molnár, Z. & Davies, K. E. Laf4/Aff3, a gene involved in intellectual disability, is required for cellular migration in the mouse cerebral cortex. PloS One 9, e105933 (2014).

120. Voisin, N., Schnur, R. E., Douzgou, S., Hiatt, S. M., Rustad, C. F., Brown, N. J., Earl, D. L., Keren, B., Levchenko, O., Geuer, S., Verheyen, S., Johnson, D., Zarate, Y. A., Hančárová, M., Amor, D. J., Bebin, E. M., Blatterer, J., Brusco, A., Cappuccio, G., Charrow, J., Chatron, N., Cooper, G. M., Courtin, T., Dadali, E., Delafontaine, J., Del Giudice, E., Doco, M., Douglas, G., Eisenkölbl, A., Funari, T., Giannuzzi, G., Gruber-Sedlmayr, U., Guex, N., Heron, D., Holla, Ø. L., Hurst, A. C. E., Juusola, J., Kronn, D., Lavrov, A., Lee, C., Lorrain, S., Merckoll, E., Mikhaleva, A., Norman, J., Pradervand, S., Prchalová, D., Rhodes, L., Sanders, V. R., Sedláček, Z., Seebacher, H. A., Sellars, E. A., Sirchia, F., Takenouchi, T., Tanaka, A. J., Taska-Tench, H., Tønne, E., Tveten, K., Vitiello, G., Vlčková, M., Uehara, T., Nava, C., Yalcin, B., Kosaki, K., Donnai, D., Mundlos, S., Brunetti-Pierri, N., Chung, W. K. & Reymond, A. Variants in the degron of AFF3 are associated with intellectual disability, mesomelic dysplasia, horseshoe kidney, and epileptic encephalopathy. Am. J. Hum. Genet. 108, 857–873 (2021).

121. Sheila, M., Narayanan, G., Ma, S., Tam, W. L., Chai, J. & Stanton, L. W. Phenotypic and molecular features underlying neurodegeneration of motor neurons derived from spinal and bulbar muscular atrophy patients. Neurobiol. Dis. 124, 1–13 (2019).

122. Lencer, R., Mills, L. J., Alliey-Rodriguez, N., Shafee, R., Lee, A. M., Reilly, J. L., Sprenger, A., McDowell, J. E., McCarroll, S. A., Keshavan, M. S., Pearlson, G. D., Tamminga, C. A., Clementz, B. A., Gershon, E. S., Sweeney, J. A. & Bishop, J. R. Genome-wide association studies of smooth pursuit and antisaccade eye movements in psychotic disorders: findings from the B-SNIP study. Transl. Psychiatry 7, e1249–e1249 (2017).

123. Cho, J., Yu, N.-K., Choi, J.-H., Sim, S.-E., Kang, S. J., Kwak, C., Lee, S.-W., Kim, J., Choi, D. I., Kim, V. N. & Kaang, B.-K. Multiple repressive mechanisms in the hippocampus during memory formation. Science 350, 82–87 (2015).

124. Regulska, M., Szuster-Głuszczak, M., Trojan, E., Leśkiewicz, M. & Basta-Kaim, A. The Emerging Role of the Double-Edged Impact of Arachidonic Acid- Derived Eicosanoids in the Neuroinflammatory Background of Depression. Curr. Neuropharmacol. 19, 278–293 (2020).

125. Dong, Y. Y., Pike, A. C. W., Mackenzie, A., McClenaghan, C., Aryal, P., Dong, L., Quigley, A., Grieben, M., Goubin, S., Mukhopadhyay, S., Ruda, G. F., Clausen, M. V., Cao, L., Brennan, P. E., Burgess-Brown, N. A., Sansom, M. S. P., Tucker, S. J. & Carpenter, E. P. K2P channel gating mechanisms revealed by structures of TREK-2 and a complex with Prozac. Science 347, 1256–1259 (2015).

126. Ansari, U., Chen, V., Sedighi, R., Syed, B., Muttalib, Z., Ansari, K., Ansari, F., Nadora, D., Razick, D. & Lui, F. Role of the UNC13 family in human diseases: A literature review. AIMS Neurosci. 10, 388–400 (2023).

127. Barak, B., Williams, A., Bielopolski, N., Gottfried, I., Okun, E., Brown, M. A., Matti, U., Rettig, J., Stuenkel, E. L. & Ashery, U. Tomosyn expression pattern in the mouse hippocampus suggests both presynaptic and postsynaptic functions. Front. Neuroanat. 4, 149 (2010).

128. Batten, S. R., Matveeva, E. A., Whiteheart, S. W., Vanaman, T. C., Gerhardt, G. A. & Slevin, J. T. Linking kindling to increased glutamate release in the dentate gyrus of the hippocampus through the STXBP5/tomosyn-1 gene. Brain Behav. 7, e00795 (2017).

129. Dranovsky, A. & Hen, R. Hippocampal Neurogenesis: Regulation by Stress and Antidepressants. Biol. Psychiatry 59, 1136–1143 (2006).

130. Wasserman, D., Geijer, T., Sokolowski, M., Rozanov, V. & Wasserman, J. Genetic variation in the hypothalamic–pituitary–adrenocortical axis regulatory factor, T-box 19, and the angry/hostility personality trait. Genes Brain Behav. 6, 321–328 (2007).

131. Gulyaeva, N. V. Functional Neurochemistry of the Ventral and Dorsal Hippocampus: Stress, Depression, Dementia and Remote Hippocampal Damage. Neurochem. Res. 44, 1306–1322 (2019).

132. Byun, Y. G., Kim, N.-S., Kim, G., Jeon, Y.-S., Choi, J. B., Park, C.-W., Kim, K., Jang, H., Kim, J., Kim, E., Han, Y.-M., Yoon, K.-J., Lee, S.-H. & Chung, W.-S. Stress induces behavioral abnormalities by increasing expression of phagocytic receptor MERTK in astrocytes to promote synapse phagocytosis. Immunity 56, 2105–2120.e13 (2023).

133. Scoville, D. W., Kang, H. S. & Jetten, A. M. GLIS1-3: emerging roles in reprogramming, stem and progenitor cell differentiation and maintenance. Stem Cell Investig. 4, 80 (2017).

134. Miller, A. H. & Raison, C. L. The role of inflammation in depression: from evolutionary imperative to modern treatment target. Nat. Rev. Immunol. 16, 22–34 (2016).

135. Price, R. B. & Duman, R. Neuroplasticity in cognitive and psychological mechanisms of depression: an integrative model. Mol. Psychiatry 25, 530–543 (2020).

136. Calcia, M. A., Bonsall, D. R., Bloomfield, P. S., Selvaraj, S., Barichello, T. & Howes, O. D. Stress and neuroinflammation: a systematic review of the effects of stress on microglia and the implications for mental illness. Psychopharmacology (Berl.) 233, 1637–1650 (2016).

137. Mariani, N., Everson, J., Pariante, C. M. & Borsini, A. Modulation of microglial activation by antidepressants. J. Psychopharmacol. (Oxf.) 36, 131–150 (2022).

138. Zhang, Z., Guo, Z., Jin, P., Yang, H., Hu, M., Zhang, Y., Tu, Z. & Hou, S. Transcriptome Profiling of Hippocampus After Cerebral Hypoperfusion in Mice. J. Mol. Neurosci. 73, 423–436 (2023).

139. Hu, X., Leak, R. K., Shi, Y., Suenaga, J., Gao, Y., Zheng, P. & Chen, J. Microglial and macrophage polarization—new prospects for brain repair. Nat. Rev. Neurol. 11, 56–64 (2015).

140. Cope, E. C. & Gould, E. Adult Neurogenesis, Glia, and the Extracellular Matrix. Cell Stem Cell 24, 690–705 (2019).

141. You, J.-E., Kim, E.-J., Kim, H. W., Kim, J.-S., Kim, K. & Kim, P.-H. Exploring the Role of Guanylate-Binding Protein-2 in Activated Microglia-Mediated Neuroinflammation and Neuronal Damage. Biomedicines 12, 1130 (2024).

142. Scott, R. S., McMahon, E. J., Pop, S. M., Reap, E. A., Caricchio, R., Cohen, P. L., Earp, H. S. & Matsushima, G. K. Phagocytosis and clearance of apoptotic cells is mediated by MER. Nature 411, 207–211 (2001).

143. Nguyen, L. T., Aprico, A., Nwoke, E., Walsh, A. D., Blades, F., Avneri, R., Martin, E., Zalc, B., Kilpatrick, T. J. & Binder, M. D. Mertk-expressing microglia influence oligodendrogenesis and myelin modelling in the CNS. J. Neuroinflammation 20, 253 (2023).

144. Matsuda, T., Murao, N., Katano, Y., Juliandi, B., Kohyama, J., Akira, S., Kawai, T. & Nakashima, K. TLR9 signalling in microglia attenuates seizure-induced aberrant neurogenesis in the adult hippocampus. Nat. Commun. 6, 6514 (2015).

145. Hung, Y.-Y., Huang, K.-W., Kang, H.-Y., Huang, G. Y.-L. & Huang, T.-L. Antidepressants normalize elevated Toll-like receptor profile in major depressive disorder. Psychopharmacology (Berl.) 233, 1707–1714 (2016).

146. Shyn, S. I., Shi, J., Kraft, J. B., Potash, J. B., Knowles, J. A., Weissman, M. M., Garriock, H. A., Yokoyama, J. S., McGrath, P. J., Peters, E. J., Scheftner, W. A., Coryell, W., Lawson, W. B., Jancic, D., Gejman, P. V., Sanders, A. R., Holmans, P., Slager, S. L., Levinson, D. F. & Hamilton, S. P. Novel loci for major depression identified by genome-wide association study of Sequenced Treatment Alternatives to Relieve Depression and meta-analysis of three studies. Mol. Psychiatry 16, 202–215 (2011).

147. Gonda, X., Eszlari, N., Anderson, I. M., Deakin, J. F. W., Bagdy, G. & Juhasz, G. Association of ATP6V1B2 rs1106634 with lifetime risk of depression and hippocampal neurocognitive deficits: possible novel mechanisms in the etiopathology of depression. Transl. Psychiatry 6, e945–e945 (2016).

148. Barbu, M. C., Huider, F., Campbell, A., Amador, C., Adams, M. J., Lynall, M.-E., Howard, D. M., Walker, R. M., Morris, S. W., Van Dongen, J., Porteous, D. J., Evans, K. L., Bullmore, E., Willemsen, G., Boomsma, D. I., Whalley, H. C. & McIntosh, A. M. Methylome-wide association study of antidepressant use in Generation Scotland and the Netherlands Twin Register implicates the innate immune system. Mol. Psychiatry 27, 1647–1657 (2022).

149. Nishi, T. & Forgac, M. The vacuolar (H+)-ATPases — nature’s most versatile proton pumps. Nat. Rev. Mol. Cell Biol. 3, 94–103 (2002).

150. Collins, M. P. & Forgac, M. Regulation and function of V-ATPases in physiology and disease. Biochim. Biophys. Acta Biomembr. 1862, 183341 (2020).

151. Li, Y., Dai, Y. & Chu, L. V-ATPase B2 promotes microglial phagocytosis of myelin debris by inactivating the MAPK signaling pathway. Neuropeptides 106, 102436 (2024).

152. Fairley, L. H., Lai, K. O., Wong, J. H., Chong, W. J., Vincent, A. S., D’Agostino, G., Wu, X., Naik, R. R., Jayaraman, A., Langley, S. R., Ruedl, C. & Barron, A. M. Mitochondrial control of microglial phagocytosis by the translocator protein and hexokinase 2 in Alzheimer’s disease. Proc. Natl. Acad. Sci. 120, e2209177120 (2023).

153. Sharma, K., Schmitt, S., Bergner, C. G., Tyanova, S., Kannaiyan, N., Manrique-Hoyos, N., Kongi, K., Cantuti, L., Hanisch, U.-K., Philips, M.-A., Rossner, M. J., Mann, M. & Simons, M. Cell type– and brain region–resolved mouse brain proteome. Nat. Neurosci. 18, 1819–1831 (2015).

154. Moriyama, Y. & Futai, M. H+-ATPase, a primary pump for accumulation of neurotransmitters, is a major constituent of brain synaptic vesicles. Biochem. Biophys. Res. Commun. 173, 443–448 (1990).

155. Eaton, A. F., Merkulova, M. & Brown, D. The H^+^ -ATPase (V-ATPase): from proton pump to signaling complex in health and disease. Am. J. Physiol.-Cell Physiol. 320, C392–C414 (2021).

156. Hinton, A., Bond, S. & Forgac, M. V-ATPase functions in normal and disease processes. Pflüg. Arch. - Eur. J. Physiol. 457, 589–598 (2009).

157. Egashira, Y., Takase, M. & Takamori, S. Monitoring of Vacuolar-Type H^+^ ATPase-Mediated Proton Influx into Synaptic Vesicles. J. Neurosci. 35, 3701–3710 (2015).

158. Abbas, Y. M., Wu, D., Bueler, S. A., Robinson, C. V. & Rubinstein, J. L. Structure of V-ATPase from the mammalian brain. Science 367, 1240–1246 (2020).

159. Pamarthy, S., Kulshrestha, A., Katara, G. K. & Beaman, K. D. The curious case of vacuolar ATPase: regulation of signaling pathways. Mol. Cancer 17, 41 (2018).

160. Li, N., Lee, B., Liu, R.-J., Banasr, M., Dwyer, J. M., Iwata, M., Li, X.-Y., Aghajanian, G. & Duman, R. S. mTOR-Dependent Synapse Formation Underlies the Rapid Antidepressant Effects of NMDA Antagonists. Science 329, 959–964 (2010).

161. Filipović, D., Costina, V., Findeisen, P. & Inta, D. Fluoxetine Enhances Synaptic Vesicle Trafficking and Energy Metabolism in the Hippocampus of Socially Isolated Rats. Int. J. Mol. Sci. 23, 15351 (2022).

162. Barazany, D. & Assaf, Y. Visualization of Cortical Lamination Patterns with Magnetic Resonance Imaging. Cereb. Cortex 22, 2016–2023 (2012).

163. Jorstad, N. L., Close, J., Johansen, N., Yanny, A. M., Barkan, E. R., Travaglini, K. J., Bertagnolli, D., Campos, J., Casper, T., Crichton, K., Dee, N., Ding, S.-L., Gelfand, E., Goldy, J., Hirschstein, D., Kiick, K., Kroll, M., Kunst, M., Lathia, K., Long, B., Martin, N., McMillen, D., Pham, T., Rimorin, C., Ruiz, A., Shapovalova, N., Shehata, S., Siletti, K., Somasundaram, S., Sulc, J., Tieu, M., Torkelson, A., Tung, H., Callaway, E. M., Hof, P. R., Keene, C. D., Levi, B. P., Linnarsson, S., Mitra, P. P., Smith, K., Hodge, R. D., Bakken, T. E. & Lein, E. S. Transcriptomic cytoarchitecture reveals principles of human neocortex organization. Science 382, eadf6812 (2023).

164. Preuss, T. M. & Wise, S. P. Evolution of prefrontal cortex. Neuropsychopharmacology 47, 3–19 (2022).

165. Eid, R. S., Gobinath, A. R. & Galea, L. A. M. Sex differences in depression: Insights from clinical and preclinical studies. Prog. Neurobiol. 176, 86–102 (2019).

166. Willard, S. L., Uberseder, B., Clark, A., Daunais, J. B., Johnston, W. D., Neely, D., Massey, A., Williamson, J. D., Kraft, R. A., Bourland, J. D., Jones, S. R. & Shively, C. A. Long term sertraline effects on neural structures in depressed and nondepressed adult female nonhuman primates. Neuropharmacology 99, 369–378 (2015).

167. Prado, C. E., Watt, S. & Crowe, S. F. A meta-analysis of the effects of antidepressants on cognitive functioning in depressed and non-depressed samples. Neuropsychol. Rev. 28, 32–72 (2018).

