## Supplemental Methods, Results, Small Tables, Figures, and Legends for "The Converging Effects of Different Categories of Antidepressants on the Brain: A Systematic Meta-Analysis of Public Transcriptional Profiling Data from the Hippocampus and Cortex"

**<u>Supplemental Tables, Figures, and Legends for:</u>**

**Corresponding author:**

Megan Hagenauer

#### Supplemental Table Legends

***Table S1. The input used for both the hippocampal and cortical meta-analyses.*** *This .xlsx file includes 5 worksheets: 1) The worksheet "HPC_Log2FoldChanges" provides the log(2) fold changes for each gene (identified by mouse and rat Entrez id: 31,493 rows) for each of the 22 antidepressant treatment vs. control comparisons included in the hippocampal meta-analysis (identified by gene expression omnibus accession # and treatment name). 2) The worksheet "HPC_SamplingVariances" provides the sampling variances for each gene to accompany the log(2) fold changes in worksheet #1. 3) The worksheet "CTX_Log2FoldChanges" provides the log(2) fold changes for each gene (identified by mouse and rat Entrez id: 36,110 rows) for each of the 16 antidepressant treatment vs. control comparisons included in the cortical meta-analysis (identified by gene expression omnibus accession # and treatment name). 4) The worksheet "CTX_SamplingVariances" provides the sampling variances for each gene to accompany the log(2) fold changes in worksheet #3. 5) The worksheet "Gene_Annotation_Orthology" contains a version of the Jackson Labs Mouse Ortholog Database that has been trimmed to one-to-one rat-mouse orthologs with accompanying annotation (gene Entrez id, gene symbol, genetic location, gene name).*

***Table S2. The full hippocampal antidepressant meta-analysis results (16,494 genes, 16,439 stable meta-analysis estimates).*** *This .xlsx file includes two worksheets: 1) The worksheet "metaOutputByPval" provides the full meta-analysis results, with each row representing the results for one gene, and each column providing either gene annotation or meta-analysis statistical output. The results are ordered by p-value, so that the top rows in the worksheet are the genes with the smallest p-values. 2) The worksheet "Column Definitions" provides the definitions for the variables present in each column in "metaOutputByPval".*

***Table S3. Hippocampal meta-analysis reveals genes that are consistently upregulated following antidepressant treatment (DEGs FDR<0.05).*** *The rows are ordered by decreasing Log2FC (highest to lowest). Positive Log2FC value denotes that a gene's expression is higher in the experimental group compared to the control group with increasingly positive values signifying stronger effect. Definitions: Log2FC = Log2 Fold Change (Antidepressant treatment vs. Control), SE = Standard Error, CI_lb=confidence interval lower bound, CI_ub=confidence interval upper bound, p-value = nominal p-value, FDR = False Discovery Rate (q-value). The #Comparisons represents the number of treatments that included measurements for the gene following quality control. Full results for all genes can be found in* ***Table S2****.*

| Gene Symbol | Mouse Entrez Gene ID | Log2FC Estimate | SE | CI_lb | CI_ub | # Contrasts | Pval | FDR |
|---|---|---|---|---|---|---|---|---|
| ***Plekha2*** | 83436 | 0.248 | 0.0659 | 0.118 | 0.377 | 13 | 1.72E-04 | 4.99E-02 |
| ***Acan*** | 11595 | 0.165 | 0.0299 | 0.106 | 0.224 | 12 | 3.47E-08 | 1.72E-04 |
| ***Glis1*** | 230587 | 0.156 | 0.0233 | 0.111 | 0.202 | 14 | 2.18E-11 | 3.60E-07 |
| ***Fam135b*** | 70363 | 0.156 | 0.0333 | 0.091 | 0.221 | 12 | 2.80E-06 | 4.20E-03 |
| ***Tnni1*** | 21952 | 0.144 | 0.0336 | 0.078 | 0.210 | 19 | 1.90E-05 | 1.31E-02 |
| ***Dmrtb1*** | 56296 | 0.135 | 0.0343 | 0.068 | 0.202 | 19 | 8.65E-05 | 3.48E-02 |
| ***Gbp2*** | 14469 | 0.124 | 0.0311 | 0.063 | 0.185 | 12 | 6.72E-05 | 2.94E-02 |
| ***Slc35d2*** | 70484 | 0.113 | 0.0279 | 0.058 | 0.167 | 19 | 5.55E-05 | 2.69E-02 |
| ***Lefty1*** | 13590 | 0.111 | 0.0272 | 0.058 | 0.165 | 15 | 4.04E-05 | 2.30E-02 |

| | | | | | | | | |
|---|---|---|---|---|---|---|---|---|
| ***Zfp41*** | 22701 | 0.107 | 0.0279 | 0.053 | 0.162 | 13 | 1.21E-04 | 4.23E-02 |
| ***Iqcg*** | 69707 | 0.107 | 0.0243 | 0.059 | 0.155 | 20 | 1.08E-05 | 9.39E-03 |
| ***Zfp691*** | 195522 | 0.105 | 0.0277 | 0.051 | 0.160 | 22 | 1.45E-04 | 4.52E-02 |
| ***Upk1a*** | 109637 | 0.099 | 0.0190 | 0.062 | 0.136 | 12 | 1.89E-07 | 5.29E-04 |
| ***Tlr9*** | 81897 | 0.096 | 0.0176 | 0.062 | 0.131 | 16 | 4.17E-08 | 1.72E-04 |
| ***Stxbp5*** | 78808 | 0.096 | 0.0229 | 0.051 | 0.141 | 20 | 2.74E-05 | 1.63E-02 |
| ***Aff3*** | 16764 | 0.096 | 0.0184 | 0.060 | 0.132 | 12 | 1.92E-07 | 5.29E-04 |
| ***Cdca7l*** | 217946 | 0.094 | 0.0197 | 0.056 | 0.133 | 20 | 1.68E-06 | 2.77E-03 |
| ***Zfp324*** | 243834 | 0.091 | 0.0189 | 0.054 | 0.128 | 14 | 1.43E-06 | 2.63E-03 |
| ***Slc25a45*** | 107375 | 0.086 | 0.0193 | 0.048 | 0.124 | 22 | 9.34E-06 | 8.55E-03 |
| ***Ccrl2*** | 54199 | 0.084 | 0.0189 | 0.047 | 0.122 | 21 | 8.05E-06 | 8.30E-03 |
| ***Krt82*** | 114566 | 0.084 | 0.0200 | 0.045 | 0.124 | 16 | 2.49E-05 | 1.58E-02 |
| ***Insrr*** | 23920 | 0.084 | 0.0222 | 0.041 | 0.127 | 17 | 1.48E-04 | 4.52E-02 |
| ***Tbx19*** | 83993 | 0.083 | 0.0216 | 0.040 | 0.125 | 18 | 1.30E-04 | 4.38E-02 |
| ***Gpr160*** | 71862 | 0.079 | 0.0211 | 0.038 | 0.121 | 20 | 1.75E-04 | 4.99E-02 |
| ***Zc3h15*** | 69082 | 0.079 | 0.0176 | 0.045 | 0.114 | 12 | 7.21E-06 | 8.30E-03 |
| ***Hid1*** | 217310 | 0.078 | 0.0205 | 0.038 | 0.118 | 22 | 1.44E-04 | 4.52E-02 |
| ***Dusp9*** | 75590 | 0.077 | 0.0192 | 0.040 | 0.115 | 21 | 5.42E-05 | 2.69E-02 |
| ***Zfp773*** | 76373 | 0.077 | 0.0201 | 0.038 | 0.117 | 11 | 1.15E-04 | 4.22E-02 |
| ***Hgsnat*** | 52120 | 0.074 | 0.0166 | 0.041 | 0.106 | 12 | 9.24E-06 | 8.55E-03 |
| ***Dcaf10*** | 242418 | 0.072 | 0.0180 | 0.037 | 0.108 | 22 | 5.85E-05 | 2.76E-02 |
| ***Relch*** | 227446 | 0.063 | 0.0159 | 0.032 | 0.094 | 22 | 7.42E-05 | 3.06E-02 |
| ***Cdkn2d*** | 12581 | 0.062 | 0.0143 | 0.034 | 0.090 | 12 | 1.31E-05 | 1.08E-02 |
| ***Sdhaf3*** | 71238 | 0.060 | 0.0142 | 0.032 | 0.088 | 21 | 2.24E-05 | 1.48E-02 |
| ***Fen1*** | 14156 | 0.060 | 0.0158 | 0.029 | 0.090 | 22 | 1.60E-04 | 4.80E-02 |
| ***Naa50*** | 72117 | 0.055 | 0.0144 | 0.027 | 0.083 | 21 | 1.46E-04 | 4.52E-02 |

***Table S4. Hippocampal meta-analysis reveals genes that are consistently downregulated following antidepressant treatment (DEGs FDR<0.05).*** *The rows are ordered by increasing Log2FC value (lowest to highest). Negative Log2FC value denotes that a gene's expression is higher in the experimental group compared to the control group, with increasingly negative values signifying stronger effect. Full results for all genes can be found in* ***Table S2****.*

| Gene Symbol | Mouse Entrez Gene ID | Log2FC Estimate | SE | CI_lb | CI_ub | # Contrasts | Pval | FDR |
|---|---|---|---|---|---|---|---|---|
| ***Fbxw10*** | 213980 | -0.206 | 0.0398 | -0.284 | -0.128 | 12 | 2.27E-07 | 5.35E-04 |
| ***Unc13c*** | 208898 | -0.191 | 0.0445 | -0.278 | -0.103 | 22 | 1.85E-05 | 1.31E-02 |
| ***Or13a28*** | 18362 | -0.160 | 0.0401 | -0.239 | -0.081 | 11 | 6.78E-05 | 2.94E-02 |
| ***Ccdc160*** | 434778 | -0.154 | 0.0306 | -0.214 | -0.094 | 14 | 5.01E-07 | 1.03E-03 |
| ***Mertk*** | 17289 | -0.138 | 0.0319 | -0.201 | -0.076 | 21 | 1.52E-05 | 1.19E-02 |
| ***Kcnk10*** | 72258 | -0.138 | 0.0362 | -0.209 | -0.067 | 22 | 1.42E-04 | 4.52E-02 |
| ***C1qtnf7*** | 109323 | -0.136 | 0.0215 | -0.178 | -0.094 | 19 | 2.19E-10 | 1.80E-06 |
| ***Cables1*** | 63955 | -0.116 | 0.0259 | -0.166 | -0.065 | 20 | 7.52E-06 | 8.30E-03 |
| ***Gpr182*** | 11536 | -0.116 | 0.0296 | -0.174 | -0.058 | 22 | 9.27E-05 | 3.55E-02 |
| ***Ankrd7*** | 75196 | -0.107 | 0.0279 | -0.162 | -0.053 | 20 | 1.18E-04 | 4.23E-02 |
| ***Ccl20*** | 20297 | -0.107 | 0.0235 | -0.153 | -0.061 | 15 | 5.64E-06 | 7.75E-03 |
| ***Lrrc1*** | 214345 | -0.091 | 0.0204 | -0.131 | -0.051 | 22 | 8.00E-06 | 8.30E-03 |
| ***Nrsn1*** | 22360 | -0.089 | 0.0205 | -0.129 | -0.048 | 20 | 1.59E-05 | 1.19E-02 |
| ***Pea15a*** | 18611 | -0.087 | 0.0217 | -0.129 | -0.044 | 22 | 6.55E-05 | 2.94E-02 |
| ***Hexd*** | 238023 | -0.078 | 0.0187 | -0.115 | -0.042 | 20 | 2.77E-05 | 1.63E-02 |

| | | | | | | | | |
|---|---|---|---|---|---|---|---|---|
| ***Sim2*** | 20465 | -0.077 | 0.0193 | -0.115 | -0.039 | 21 | 6.99E-05 | 2.96E-02 |
| ***Csad*** | 246277 | -0.075 | 0.0200 | -0.115 | -0.036 | 22 | 1.73E-04 | 4.99E-02 |
| ***Clcn2*** | 12724 | -0.072 | 0.0177 | -0.107 | -0.038 | 21 | 4.54E-05 | 2.50E-02 |
| ***Map3k1*** | 26401 | -0.072 | 0.0178 | -0.107 | -0.037 | 22 | 4.89E-05 | 2.60E-02 |
| ***C2cd2*** | 207781 | -0.070 | 0.0178 | -0.104 | -0.035 | 21 | 8.99E-05 | 3.53E-02 |
| ***Krt27*** | 16675 | -0.069 | 0.0179 | -0.104 | -0.034 | 18 | 1.28E-04 | 4.38E-02 |
| ***Xrcc6*** | 14375 | -0.061 | 0.0158 | -0.092 | -0.030 | 22 | 1.09E-04 | 4.08E-02 |
| ***Rpp25l*** | 69961 | -0.060 | 0.0149 | -0.089 | -0.031 | 22 | 5.23E-05 | 2.69E-02 |

***Table S5: Hippocampal non-directional fast Gene Set Enrichment Analysis (fGSEA) results (10,975 gene sets).*** *This .xlsx file includes two worksheets: 1) The worksheet "Non_directional_WholeHPC_GSEA" provides the full fGSEA results, with each row representing the results for one gene set, and each column providing the fGSEA statistical output. The results are ordered by p-value, so that the top rows in the worksheet are the gene sets with the smallest p-values. 2) The worksheet "Column Definitions" provides the definitions for the variables present in each column in "Non_directional_WholeHPC_GSEA".*

***Table S6: Hippocampal directional fast Gene Set Enrichment Analysis (fGSEA) results (10,975 gene sets).*** *This .xlsx file includes two worksheets: 1) The worksheet "Directional_Whole_HPC_GSEA" provides the full fGSEA results, with each row representing the results for one gene set, and each column providing the fGSEA statistical output. The results are ordered by p-value, so that the top rows in the worksheet are the gene sets with the smallest p-values. 2) The worksheet "Column Definitions" provides the definitions for the variables present in each column in "Directional_Whole_HPC_GSEA".*

***Table S7. 58 genes showed significant evidence of publication bias when considering the full hippocampal meta-analysis output (Egger's test FDR<0.05).*** *The Egger's regression test is used to detect funnel plot asymmetry, which is a commonly used indicator of publication bias. Rows in the table are ordered by the p-value from the Egger's regression test ("Egger Pval"), provided with the associated Z statistic ("Egger Zstat") and false discovery rate ("Egger FDR"). Other columns follow the conventions used in* ***Table S3*** *and* ***Table S4****, but with "AD" listed before each statistic to signify "Antidepressant Effect". Full results for all genes can be found in* ***Table S2****.*

| *Gene Symbol* | *Mouse Entrez Gene ID* | *AD Log2FC Estimate* | *AD SE* | *AD CI_lb* | *AD CI_ub* | *# Contrasts* | *AD Pval* | *AD FDR* | *Egger Zstat* | *Egger Pval* | *Egger FDR* |
|---|---|---|---|---|---|---|---|---|---|---|---|
| ***Cd302*** | *66205* | *-0.05* | *0.03* | *-0.11* | *0.01* | *20* | *9.01E-02* | *4.66E-01* | *-5.94* | *2.80E-09* | *4.62E-05* |
| ***Lhfpl6*** | *108927* | *0.13* | *0.09* | *-0.05* | *0.31* | *22* | *1.67E-01* | *5.71E-01* | *5.45* | *4.94E-08* | *4.08E-04* |
| ***Pnck*** | *93843* | *-0.15* | *0.07* | *-0.28* | *-0.02* | *21* | *2.69E-02* | *3.36E-01* | *-5.34* | *9.27E-08* | *5.10E-04* |
| ***Podxl2*** | *319655* | *0.11* | *0.06* | *-0.01* | *0.22* | *22* | *7.93E-02* | *4.49E-01* | *5.20* | *1.98E-07* | *8.16E-04* |
| ***Csgalnact1*** | *234356* | *-0.13* | *0.07* | *-0.26* | *0.00* | *22* | *5.74E-02* | *4.12E-01* | *-5.12* | *3.12E-07* | *1.03E-03* |
| ***Jun*** | *16476* | *-0.09* | *0.04* | *-0.18* | *0.00* | *22* | *4.41E-02* | *3.82E-01* | *-5.05* | *4.51E-07* | *1.24E-03* |
| ***Cdkn2b*** | *12579* | *0.00* | *0.03* | *-0.06* | *0.07* | *22* | *8.94E-01* | *9.71E-01* | *-4.95* | *7.59E-07* | *1.79E-03* |
| ***Rflnb*** | *76566* | *0.15* | *0.05* | *0.04* | *0.25* | *20* | *7.23E-03* | *2.25E-01* | *4.90* | *9.65E-07* | *1.99E-03* |
| ***Xpnpep2*** | *170745* | *-0.08* | *0.05* | *-0.18* | *0.02* | *21* | *1.17E-01* | *5.09E-01* | *-4.81* | *1.52E-06* | *2.59E-03* |
| ***Rps15a*** | *267019* | *-0.09* | *0.08* | *-0.25* | *0.06* | *12* | *2.42E-01* | *6.36E-01* | *-4.78* | *1.73E-06* | *2.59E-03* |
| ***Zbtb8b*** | *215627* | *-0.01* | *0.03* | *-0.06* | *0.05* | *22* | *8.01E-01* | *9.36E-01* | *-4.78* | *1.73E-06* | *2.59E-03* |
| ***Calb1*** | *12307* | *-0.17* | *0.09* | *-0.35* | *0.01* | *21* | *5.77E-02* | *4.12E-01* | *-4.77* | *1.89E-06* | *2.59E-03* |
| ***Cdk18*** | *18557* | *0.06* | *0.05* | *-0.04* | *0.16* | *21* | *2.73E-01* | *6.62E-01* | *4.67* | *3.02E-06* | *3.83E-03* |
| ***Insl5*** | *23919* | *0.00* | *0.09* | *-0.18* | *0.18* | *11* | *9.73E-01* | *9.96E-01* | *4.55* | *5.38E-06* | *6.34E-03* |

| | | | | | | | | | | | |
|---|---|---|---|---|---|---|---|---|---|---|---|
| ***Ahdc1*** | 230793 | 0.02 | 0.04 | -0.07 | 0.10 | 18 | 6.67E-01 | 8.85E-01 | 4.50 | 6.70E-06 | 7.37E-03 |
| ***Gphb5*** | 217674 | 0.15 | 0.17 | -0.18 | 0.47 | 12 | 3.81E-01 | 7.38E-01 | 4.43 | 9.38E-06 | 9.48E-03 |
| ***Chrnb3*** | 108043 | 0.05 | 0.03 | 0.00 | 0.10 | 19 | 6.09E-02 | 4.18E-01 | 4.42 | 9.78E-06 | 9.48E-03 |
| ***Tnnt1*** | 21955 | 0.00 | 0.05 | -0.10 | 0.10 | 22 | 9.70E-01 | 9.96E-01 | 4.39 | 1.14E-05 | 1.00E-02 |
| ***Egr4*** | 13656 | 0.02 | 0.04 | -0.06 | 0.10 | 22 | 5.85E-01 | 8.50E-01 | 4.39 | 1.15E-05 | 1.00E-02 |
| ***Chst7*** | 60322 | 0.06 | 0.03 | 0.01 | 0.12 | 21 | 3.15E-02 | 3.48E-01 | 4.33 | 1.52E-05 | 1.26E-02 |
| ***Cybb*** | 13058 | -0.04 | 0.05 | -0.13 | 0.05 | 20 | 4.14E-01 | 7.59E-01 | -4.30 | 1.68E-05 | 1.32E-02 |
| ***Tdo2*** | 56720 | -0.21 | 0.13 | -0.47 | 0.05 | 19 | 1.07E-01 | 4.90E-01 | -4.21 | 2.53E-05 | 1.89E-02 |
| ***Rerg*** | 232441 | -0.04 | 0.06 | -0.15 | 0.07 | 14 | 4.91E-01 | 8.04E-01 | -4.19 | 2.85E-05 | 2.04E-02 |
| ***Ccrl1l*** | 12770 | 0.05 | 0.03 | -0.01 | 0.11 | 17 | 9.41E-02 | 4.72E-01 | 4.17 | 3.00E-05 | 2.06E-02 |
| ***Slc22a7*** | 108114 | -0.02 | 0.03 | -0.08 | 0.05 | 20 | 6.09E-01 | 8.62E-01 | -4.15 | 3.26E-05 | 2.08E-02 |
| ***Cdhr2*** | 268663 | 0.05 | 0.04 | -0.03 | 0.12 | 14 | 2.16E-01 | 6.15E-01 | 4.15 | 3.34E-05 | 2.08E-02 |
| ***Tubgcp2*** | 74237 | -0.04 | 0.03 | -0.09 | 0.01 | 22 | 1.22E-01 | 5.13E-01 | -4.15 | 3.40E-05 | 2.08E-02 |
| ***Fam43b*** | 625638 | 0.18 | 0.09 | 0.01 | 0.35 | 11 | 3.76E-02 | 3.66E-01 | 4.10 | 4.10E-05 | 2.41E-02 |
| ***Npy*** | 109648 | 0.10 | 0.09 | -0.07 | 0.28 | 21 | 2.52E-01 | 6.45E-01 | 4.08 | 4.58E-05 | 2.58E-02 |
| ***Stra6l*** | 74152 | -0.02 | 0.03 | -0.08 | 0.05 | 17 | 5.60E-01 | 8.38E-01 | 4.07 | 4.73E-05 | 2.58E-02 |
| ***Clybl*** | 69634 | -0.06 | 0.03 | -0.12 | 0.00 | 21 | 4.29E-02 | 3.80E-01 | -4.06 | 4.85E-05 | 2.58E-02 |
| ***Spon1*** | 233744 | -0.17 | 0.08 | -0.32 | -0.03 | 22 | 2.20E-02 | 3.09E-01 | -4.04 | 5.31E-05 | 2.73E-02 |
| ***Adamts6*** | 108154 | 0.05 | 0.03 | 0.00 | 0.11 | 20 | 6.15E-02 | 4.19E-01 | 4.04 | 5.46E-05 | 2.73E-02 |
| ***Card6*** | 239319 | -0.10 | 0.06 | -0.21 | 0.01 | 20 | 8.24E-02 | 4.54E-01 | -4.02 | 5.74E-05 | 2.79E-02 |
| ***2610524H06Rik*** | 330173 | 0.07 | 0.04 | -0.01 | 0.15 | 11 | 7.40E-02 | 4.40E-01 | -4.01 | 6.13E-05 | 2.86E-02 |
| ***Pdcd6ip*** | 18571 | -0.06 | 0.04 | -0.14 | 0.02 | 22 | 1.18E-01 | 5.10E-01 | -4.00 | 6.37E-05 | 2.86E-02 |
| ***Ets1*** | 23871 | -0.02 | 0.02 | -0.05 | 0.01 | 22 | 1.80E-01 | 5.81E-01 | -4.00 | 6.42E-05 | 2.86E-02 |
| ***Gpr83*** | 14608 | 0.22 | 0.13 | -0.05 | 0.48 | 22 | 1.05E-01 | 4.90E-01 | 3.98 | 6.82E-05 | 2.96E-02 |
| ***Dock3*** | 208869 | 0.04 | 0.03 | -0.02 | 0.11 | 20 | 2.01E-01 | 6.03E-01 | 3.97 | 7.11E-05 | 3.01E-02 |
| ***Gdf15*** | 23886 | 0.06 | 0.02 | 0.02 | 0.10 | 19 | 1.64E-03 | 1.40E-01 | 3.96 | 7.49E-05 | 3.01E-02 |
| ***Cdh1*** | 12550 | 0.02 | 0.02 | -0.03 | 0.07 | 22 | 3.75E-01 | 7.35E-01 | -3.96 | 7.52E-05 | 3.01E-02 |
| ***Tonsl*** | 72749 | 0.00 | 0.03 | -0.06 | 0.05 | 21 | 9.33E-01 | 9.84E-01 | -3.95 | 7.76E-05 | 3.01E-02 |
| ***Adcy10*** | 271639 | 0.03 | 0.03 | -0.02 | 0.08 | 21 | 2.91E-01 | 6.73E-01 | -3.95 | 7.86E-05 | 3.01E-02 |
| ***Arl4d*** | 80981 | -0.03 | 0.03 | -0.10 | 0.03 | 22 | 3.58E-01 | 7.24E-01 | 3.93 | 8.65E-05 | 3.18E-02 |
| ***Car4*** | 12351 | -0.15 | 0.06 | -0.26 | -0.04 | 22 | 6.46E-03 | 2.13E-01 | -3.93 | 8.67E-05 | 3.18E-02 |
| ***Pcdhga2*** | 93710 | 0.03 | 0.04 | -0.05 | 0.11 | 12 | 4.96E-01 | 8.06E-01 | 3.87 | 1.11E-04 | 3.97E-02 |
| ***Grm7*** | 108073 | 0.00 | 0.02 | -0.03 | 0.03 | 22 | 9.56E-01 | 9.92E-01 | 3.86 | 1.14E-04 | 4.02E-02 |
| ***Sema3e*** | 20349 | 0.16 | 0.11 | -0.05 | 0.38 | 21 | 1.27E-01 | 5.21E-01 | 3.85 | 1.20E-04 | 4.09E-02 |
| ***Rab26*** | 328778 | -0.38 | 0.20 | -0.77 | 0.01 | 15 | 5.40E-02 | 4.07E-01 | -3.84 | 1.21E-04 | 4.09E-02 |
| ***Col27a1*** | 373864 | 0.03 | 0.02 | -0.01 | 0.07 | 22 | 1.18E-01 | 5.10E-01 | 3.82 | 1.36E-04 | 4.49E-02 |
| ***Gpc1*** | 14733 | 0.16 | 0.06 | 0.04 | 0.27 | 17 | 9.06E-03 | 2.35E-01 | 3.80 | 1.44E-04 | 4.65E-02 |
| ***Rspo2*** | 239405 | 0.04 | 0.05 | -0.06 | 0.14 | 22 | 4.22E-01 | 7.64E-01 | 3.78 | 1.59E-04 | 4.91E-02 |
| ***Slbp*** | 20492 | -0.17 | 0.13 | -0.42 | 0.08 | 17 | 1.87E-01 | 5.88E-01 | -3.77 | 1.61E-04 | 4.91E-02 |
| ***Raver1*** | 71766 | 0.05 | 0.03 | -0.01 | 0.11 | 19 | 1.11E-01 | 4.99E-01 | 3.77 | 1.62E-04 | 4.91E-02 |
| ***Ptgds*** | 19215 | -0.04 | 0.05 | -0.14 | 0.06 | 22 | 4.16E-01 | 7.61E-01 | -3.76 | 1.69E-04 | 4.91E-02 |
| ***Gkn2*** | 66284 | -0.06 | 0.04 | -0.13 | 0.01 | 18 | 7.59E-02 | 4.43E-01 | -3.76 | 1.72E-04 | 4.91E-02 |
| ***Golm1*** | 105348 | -0.19 | 0.10 | -0.38 | 0.01 | 14 | 5.86E-02 | 4.14E-01 | -3.76 | 1.72E-04 | 4.91E-02 |
| ***Fbln2*** | 14115 | 0.26 | 0.19 | -0.11 | 0.63 | 14 | 1.67E-01 | 5.70E-01 | 3.76 | 1.73E-04 | 4.91E-02 |

***Table S8. Exploratory traditional antidepressant hippocampal meta-analysis results (9,628 genes, 9,612 stable meta-analysis estimates).*** *This .xlsx file includes two worksheets: 1) The worksheet "metaOutputByPval" provides the full meta-analysis results, with each row representing the results for one gene, and each column providing either gene annotation or meta-analysis statistical output. The results are ordered by p-value, so that the top rows in the worksheet are the genes with the smallest p-values. 2) The worksheet "Column Definitions" provides the definitions for the variables present in each column in "metaOutputByPval".*

***Table S9. Exploratory non-traditional antidepressant hippocampal meta-analysis results (12,203 genes, 12,154 stable meta-analysis estimates).*** *This .xlsx file includes two worksheets: 1) The worksheet "metaOutputByPval" provides the full meta-analysis results, with each row representing the results for one gene, and each column providing either gene annotation or meta-analysis statistical output. The results are ordered by p-value, so that the top rows in the worksheet are the genes with the smallest p-values. 2) The worksheet "Column Definitions" provides the definitions for the variables present in each column in "metaOutputByPval".*

***Table S10. Exploratory hippocampal results from a meta-regression including antidepressant type (non-traditional vs. traditional) and dissection (DG vs. whole hippocampus) as co-variates (16,494 genes, 16,442 stable meta-analysis estimates).*** *This .xlsx file includes two worksheets: 1) The worksheet "metaOutputByPval" provides the full meta-analysis results, with each row representing the results for one gene, and each column providing either gene annotation or meta-analysis statistical output. The results are ordered by p-value for the overall effect of antidepressants, so that the top rows in the worksheet are the genes with the smallest p-values. 2) The worksheet "Column Definitions" provides the definitions for the variables present in each column in "metaOutputByPval".*

***Table S11. Exploratory hippocampal results from a meta-regression including antidepressant type (non-traditional vs. traditional), dissection (DG vs. whole hippocampus), and transcriptional profiling platform (microarray vs. RNA-seq) as co-variates (16,494 genes, 15,082 stable meta-analysis estimates).*** *This .xlsx file includes two worksheets: 1) The worksheet "metaOutputByPval" provides the full meta-analysis results, with each row representing the results for one gene, and each column providing either gene annotation or meta-analysis statistical output. The results are ordered by p-value for the overall effect of antidepressants, so that the top rows in the worksheet are the genes with the smallest p-values. 2) The worksheet "Column Definitions" provides the definitions for the variables present in each column in "metaOutputByPval".*

***Table S12. Exploratory hippocampal results from a meta-regression including antidepressant type (non-traditional vs. traditional), dissection (DG vs. whole hippocampus), and inclusion of a depression model (vs. control-only) as co-variates (16,494 genes, 16,148 stable meta-analysis estimates).*** *This .xlsx file includes two worksheets: 1) The worksheet "metaOutputByPval" provides the full meta-analysis results, with each row representing the results for one gene, and each column providing either gene annotation or meta-analysis statistical output. The results are ordered by p-value for the overall effect of antidepressants, so that the top rows in the worksheet are the genes with the smallest p-values. 2) The worksheet "Column Definitions" provides the definitions for the variables present in each column in "metaOutputByPval".*

***Table S13: A comparison of the FLX-related hippocampal gene expression identified by the meta-analyses in Ibrahim et al. 2022 and our antidepressant meta-analysis results.*** *Ibrahim et al. (2022) examined the effects of the FLX on the hippocampus using meta-analyses of transcriptional profiling results from either stressed rodents (7 studies, n=75) or stress naive rodents (7 studies, n=56). Due to differing inclusion/exclusion criteria, only 66% of the samples included in their meta-analyses were in our own meta-analysis (n=87 of 131), contributing 28% to our final sample size (n=87 of 313). Therefore, their results can provide insight that is partially independent from our own. Their publication* *provided results for the top DEGs identified using two methods of integrating findings across studies (integration method and portrait method) and consensus scores indicating whether those DEGs were predominantly upregulated or downregulated for the stressed animals (Table 2 & 3) and stress-naive animals (Table 6 & Table 7). For comparison with our results, consensus scores from these tables were converted to a categorical variable indicating "Up" or "Down" direction of effect. For each gene, antidepressant Log2FC and nominal P-values are provided from each of our hippocampal meta-analyses (full hippocampal ("HPC") meta-analysis, traditional ("Trad") antidepressant-only, and non-traditional ("NonTrad") antidepressant-only. For ease of visual comparison, down-regulation following antidepressant treatment (negative Log2FC or Consensus Score) are colored blue, and upregulation following antidepressant treatment (positive Log2FC or Consensus Score) are colored pink. P-values surviving a traditional nominal threshold for significance (p<0.05) are in bold.*

| Mouse Symbol | Mouse Entrez Gene ID | Rat Entrez Gene ID | Ibrahim Direction | Ibrahim Meta-Analysis | Full HPC Meta: Log2FC | Full HPC Meta: Pval | Trad HPC Meta: Log2FC | Trad HPC Meta: Pval | NonTrad HPC Meta: Log2FC | NonTrad HPC Meta: Pval |
|---|---|---|---|---|---|---|---|---|---|---|
| Arhgef28 | 110596 | 361882 | Down | FLX_NoStress_Integration | -0.05 | 0.25089 | -0.12 | **0.02846** | 0.03 | 0.50521 |
| Arrb2 | 216869 | 25388 | Down | FLX_NoStress_Integration | -0.01 | 0.61192 | -0.01 | 0.73926 | -0.03 | 0.22127 |
| Cdon | 57810 | 50938 | Down | FLX_NoStress_Integration | -0.04 | 0.23978 | -0.04 | 0.44509 | -0.04 | 0.46428 |
| Chgb | 12653 | 24259 | Down | FLX_NoStress_Integration | 0.07 | 0.07000 | 0.10 | 0.11180 | 0.04 | 0.06566 |
| Doc2b | 13447 | 81820 | Down | FLX_NoStress_Integration | -0.07 | 0.62544 | -0.21 | 0.40088 | 0.06 | 0.36133 |
| Fat4 | 329628 | 310341 | Down | FLX_NoStress_Integration | -0.26 | **0.01903** | -0.42 | **0.02682** | -0.05 | 0.27329 |
| Gpr12 | 14738 | 80840 | Down | FLX_NoStress_Integration | -0.10 | 0.34206 | -0.24 | 0.13946 | -0.05 | 0.17188 |
| Ints10 | 70885 | 290679 | Down | FLX_NoStress_Integration | -0.02 | 0.12347 | NA | NA | -0.01 | 0.62361 |
| Isoc1 | 66307 | 364879 | Down | FLX_NoStress_Integration | -0.10 | **0.04802** | NA | NA | -0.01 | 0.56603 |
| Itga4 | 16401 | 311144 | Down | FLX_NoStress_Integration | -0.07 | 0.19594 | -0.12 | 0.16717 | -0.02 | 0.57864 |
| Itsn1 | 16443 | 29491 | Down | FLX_NoStress_Integration | -0.02 | 0.27391 | -0.03 | 0.25697 | -0.01 | 0.71699 |
| Kirrel3 | 67703 | 315546 | Down | FLX_NoStress_Integration | -0.05 | 0.12167 | -0.05 | 0.30870 | -0.03 | 0.40183 |
| Map1a | 17754 | 25152 | Down | FLX_NoStress_Integration | 0.01 | 0.45048 | NA | NA | NA | NA |
| Mpdz | 17475 | 29365 | Down | FLX_NoStress_Integration | -0.05 | 0.13256 | -0.09 | 0.07397 | -0.01 | 0.61429 |
| Negr1 | 320840 | 59318 | Down | FLX_NoStress_Integration | -0.10 | **0.00820** | -0.15 | **0.01739** | -0.03 | 0.22340 |
| Nhsl2 | 100042480 | NA | Down | FLX_NoStress_Integration | NA | NA | NA | NA | NA | NA |
| Ntrk3 | 18213 | 29613 | Down | FLX_NoStress_Integration | -0.01 | 0.58092 | -0.01 | 0.64792 | -0.02 | 0.46957 |
| Pcdh19 | 279653 | 317183 | Down | FLX_NoStress_Integration | -0.16 | 0.06381 | NA | NA | -0.04 | 0.16806 |
| Pde7b | 29863 | 140929 | Down | FLX_NoStress_Integration | -0.11 | **0.03791** | -0.15 | 0.05635 | -0.05 | 0.16187 |
| Pdlim5 | 56376 | 64353 | Down | FLX_NoStress_Integration | -0.04 | **0.04426** | -0.05 | 0.12682 | -0.04 | 0.12474 |
| Rab27a | 11891 | 50645 | Down | FLX_NoStress_Integration | -0.16 | **0.03682** | -0.23 | 0.05709 | -0.04 | 0.45440 |
| Rasgrf1 | 19417 | 192213 | Down | FLX_NoStress_Integration | -0.09 | 0.06941 | -0.14 | 0.06499 | 0.00 | 0.87570 |
| Scn3b | 235281 | 245956 | Down | FLX_NoStress_Integration | -0.02 | 0.59571 | -0.09 | 0.17028 | 0.01 | 0.76576 |
| Tnxb | 81877 | NA | Down | FLX_NoStress_Integration | -0.33 | **0.01455** | NA | NA | NA | NA |
| Zfhx2 | 239102 | 305888 | Down | FLX_NoStress_Integration | 0.03 | 0.32086 | NA | NA | -0.02 | 0.40066 |
| Zfp316 | 54201 | 304293 | Down | FLX_NoStress_Integration | -0.03 | 0.13655 | -0.03 | 0.44689 | -0.04 | 0.08244 |
| Cd68 | 12514 | 287435 | Up | FLX_NoStress_Integration | 0.14 | 0.05264 | 0.22 | **0.03405** | 0.02 | 0.77618 |
| Cfh | 12628 | NA | Up | FLX_NoStress_Integration | 0.05 | 0.24816 | NA | NA | NA | NA |
| Ddr1 | 12305 | 25678 | Up | FLX_NoStress_Integration | 0.04 | 0.36515 | 0.08 | 0.21715 | -0.01 | 0.75520 |
| Gsn | 227753 | 296654 | Up | FLX_NoStress_Integration | 0.10 | 0.09665 | 0.13 | 0.17172 | 0.06 | 0.16378 |
| H2-D1 | NA | NA | Up | FLX_NoStress_Integration | NA | NA | NA | NA | NA | NA |
| Homer1 | 26556 | 29546 | Up | FLX_NoStress_Integration | 0.07 | 0.26035 | NA | NA | NA | NA |
| Htr1b | 15551 | 25075 | Up | FLX_NoStress_Integration | -0.02 | 0.78822 | -0.08 | 0.43408 | 0.04 | 0.47219 |
| Htr5b | 15564 | 79247 | Up | FLX_NoStress_Integration | 0.12 | **0.00935** | 0.18 | **0.01168** | -0.01 | 0.70570 |
| Igfbp6 | 16012 | 25641 | Up | FLX_NoStress_Integration | 0.33 | 0.08497 | 0.62 | **0.03730** | 0.04 | 0.38266 |
| Knstrn | 51944 | 311325 | Up | FLX_NoStress_Integration | 0.07 | **0.00032** | 0.06 | **0.00186** | 0.09 | **0.02808** |
| Mat2a | 232087 | 171347 | Up | FLX_NoStress_Integration | -0.03 | 0.06010 | -0.02 | 0.43513 | -0.05 | **0.01549** |
| Mylk | 107589 | 288057 | Up | FLX_NoStress_Integration | -0.04 | 0.52346 | -0.07 | 0.44220 | 0.02 | 0.42075 |
| Myo1e | 71602 | 25484 | Up | FLX_NoStress_Integration | 0.00 | 0.83349 | NA | NA | 0.01 | 0.79355 |
| Pcdh7 | 54216 | 360942 | Up | FLX_NoStress_Integration | 0.10 | 0.14176 | 0.19 | 0.07931 | 0.02 | 0.58592 |
| Pde4b | 18578 | 24626 | Up | FLX_NoStress_Integration | 0.08 | 0.20242 | 0.14 | 0.16253 | -0.01 | 0.90279 |

| | | | | | | | | | | |
|---|---|---|---|---|---|---|---|---|---|---|
| Rassf5 | 54354 | 54355 | Up | FLX_NoStress_Integration | 0.00 | 0.99532 | 0.07 | 0.24228 | -0.02 | 0.57907 |
| S100a6 | 20200 | 85247 | Up | FLX_NoStress_Integration | 0.17 | 0.07341 | 0.33 | **0.02781** | -0.01 | 0.73898 |
| Sel1l3 | 231238 | 360945 | Up | FLX_NoStress_Integration | 0.14 | 0.05900 | NA | NA | NA | NA |
| Sema3a | 20346 | 29751 | Up | FLX_NoStress_Integration | 0.18 | 0.12547 | 0.24 | 0.23988 | 0.06 | 0.26520 |
| Sorcs1 | 58178 | 309533 | Up | FLX_NoStress_Integration | 0.14 | 0.11163 | 0.22 | 0.14544 | 0.00 | 0.88144 |
| Tfrc | 22042 | 64678 | Up | FLX_NoStress_Integration | 0.04 | 0.06712 | 0.05 | 0.14854 | 0.03 | 0.23622 |
| Trpm3 | 226025 | 309407 | Down | FLX_NoStress_Portrait | -0.05 | 0.24509 | -0.11 | 0.05519 | 0.04 | 0.42082 |
| Jun | 16476 | 24516 | Down | FLX_NoStress_Portrait | -0.09 | **0.04409** | -0.10 | 0.16682 | -0.04 | 0.06615 |
| Efnb3 | 13643 | 360546 | Down | FLX_NoStress_Portrait | -0.10 | 0.09045 | -0.13 | 0.16145 | -0.05 | 0.29331 |
| Lct | 226413 | 116569 | Down | FLX_NoStress_Portrait | -0.24 | 0.30446 | NA | NA | 0.02 | 0.52309 |
| Pdia6 | 71853 | 286906 | Down | FLX_NoStress_Portrait | -0.02 | 0.60428 | -0.05 | 0.18990 | NA | NA |
| Kcnq3 | 110862 | 29682 | Down | FLX_NoStress_Portrait | -0.04 | 0.11339 | -0.07 | **0.00648** | 0.02 | 0.41573 |
| Mcm6 | 17219 | 29685 | Down | FLX_NoStress_Portrait | -0.06 | 0.25481 | -0.10 | 0.19020 | 0.02 | 0.57793 |
| Cacna1d | 12289 | 29716 | Down | FLX_NoStress_Portrait | -0.01 | 0.61939 | 0.00 | 0.90449 | -0.04 | 0.21315 |
| Nfia | 18027 | 25492 | Down | FLX_NoStress_Portrait | 0.00 | 0.97362 | -0.06 | 0.22653 | 0.02 | 0.50891 |
| Nedd4l | 83814 | 291553 | Down | FLX_NoStress_Portrait | -0.02 | 0.33074 | -0.04 | 0.31071 | -0.03 | 0.12825 |
| Fndc1 | 68655 | 308099 | Down | FLX_NoStress_Portrait | -0.15 | **0.00805** | NA | NA | -0.04 | 0.47828 |
| Ntf3 | 18205 | 81737 | Down | FLX_NoStress_Portrait | -0.34 | 0.09909 | -0.61 | 0.06081 | -0.06 | 0.17750 |
| Itga4 | 16401 | 311144 | Down | FLX_NoStress_Portrait | -0.07 | 0.19594 | -0.12 | 0.16717 | -0.02 | 0.57864 |
| Kbtbd11 | 74901 | 306617 | Down | FLX_NoStress_Portrait | -0.02 | 0.75619 | NA | NA | -0.01 | 0.72339 |
| Kctd4 | 67516 | 691835 | Down | FLX_NoStress_Portrait | -0.03 | 0.37873 | -0.04 | 0.47371 | -0.04 | 0.27450 |
| Tnxb | 81877 | NA | Down | FLX_NoStress_Portrait | -0.33 | **0.01455** | NA | NA | NA | NA |
| Scn3b | 235281 | 245956 | Down | FLX_NoStress_Portrait | -0.02 | 0.59571 | -0.09 | 0.17028 | 0.01 | 0.76576 |
| Foxo1 | 56458 | 84482 | Down | FLX_NoStress_Portrait | -0.08 | 0.14842 | -0.13 | 0.13530 | -0.03 | 0.42102 |
| Rasgrf1 | 19417 | 192213 | Down | FLX_NoStress_Portrait | -0.09 | 0.06941 | -0.14 | 0.06499 | 0.00 | 0.87570 |
| Auts2 | 319974 | NA | Down | FLX_NoStress_Portrait | -0.06 | 0.17803 | NA | NA | NA | NA |
| Doc2b | 13447 | 81820 | Down | FLX_NoStress_Portrait | -0.07 | 0.62544 | -0.21 | 0.40088 | 0.06 | 0.36133 |
| Dsp | 109620 | 306871 | Down | FLX_NoStress_Portrait | -0.37 | 0.20205 | NA | NA | NA | NA |
| Slc4a4 | 54403 | 84484 | Down | FLX_NoStress_Portrait | -0.08 | **0.03769** | NA | NA | -0.02 | 0.57718 |
| Pcdh19 | 279653 | 317183 | Down | FLX_NoStress_Portrait | -0.16 | 0.06381 | NA | NA | -0.04 | 0.16806 |
| Fat4 | 329628 | 310341 | Down | FLX_NoStress_Portrait | -0.26 | **0.01903** | -0.42 | **0.02682** | -0.05 | 0.27329 |
| Akt3 | 23797 | 29414 | Down | FLX_NoStress_Portrait | 0.00 | 0.86977 | -0.01 | 0.74583 | 0.02 | 0.42391 |
| Sipa1l2 | 244668 | 361442 | Down | FLX_NoStress_Portrait | -0.07 | 0.11910 | NA | NA | -0.01 | 0.78764 |
| Zfp316 | 54201 | 304293 | Down | FLX_NoStress_Portrait | -0.03 | 0.13655 | -0.03 | 0.44689 | -0.04 | 0.08244 |
| Slit1 | 20562 | 65047 | Down | FLX_NoStress_Portrait | -0.02 | 0.62631 | -0.02 | 0.76075 | -0.04 | 0.40829 |
| Itsn1 | 16443 | 29491 | Down | FLX_NoStress_Portrait | -0.02 | 0.27391 | -0.03 | 0.25697 | -0.01 | 0.71699 |
| Actr10 | 56444 | 299121 | NA | FLX_NoStress_Portrait | -0.01 | 0.52894 | -0.02 | 0.32522 | 0.00 | 0.84589 |
| Vip | 22353 | 117064 | Up | FLX_NoStress_Portrait | 0.05 | 0.63564 | 0.12 | 0.42447 | 0.01 | 0.88805 |
| Gsn | 227753 | 296654 | Up | FLX_NoStress_Portrait | 0.10 | 0.09665 | 0.13 | 0.17172 | 0.06 | 0.16378 |
| Cd9 | 12527 | 24936 | Up | FLX_NoStress_Portrait | 0.05 | 0.36336 | 0.03 | 0.72934 | 0.07 | 0.20389 |
| S100a10 | 20194 | 81778 | Up | FLX_NoStress_Portrait | 0.11 | 0.06303 | 0.17 | 0.07187 | 0.02 | 0.53796 |
| Fcgr2b | 14130 | 289211 | Up | FLX_NoStress_Portrait | 0.05 | 0.24743 | 0.09 | 0.07967 | 0.01 | 0.80871 |
| Vsnl1 | 26950 | 24877 | Up | FLX_NoStress_Portrait | 0.01 | 0.79471 | 0.02 | 0.51012 | 0.00 | 0.87376 |
| Bgn | 12111 | 25181 | Up | FLX_NoStress_Portrait | 0.11 | 0.12191 | 0.19 | 0.16177 | 0.00 | 0.94691 |
| Sel1l3 | 231238 | 360945 | Up | FLX_NoStress_Portrait | 0.14 | 0.05900 | NA | NA | NA | NA |
| Ppp2r5c | 26931 | 691318 | Up | FLX_NoStress_Portrait | 0.07 | **0.04984** | 0.12 | 0.05824 | -0.01 | 0.48188 |
| Ddr1 | 12305 | 25678 | Up | FLX_NoStress_Portrait | 0.04 | 0.36515 | 0.08 | 0.21715 | -0.01 | 0.75520 |
| Anxa5 | 11747 | 25673 | Up | FLX_NoStress_Portrait | 0.02 | 0.51054 | 0.06 | 0.23923 | -0.01 | 0.70033 |
| Spock3 | 72902 | 306404 | Up | FLX_NoStress_Portrait | 0.14 | **0.01625** | 0.13 | 0.18561 | 0.09 | **0.00766** |
| Icam1 | 15894 | 25464 | Up | FLX_NoStress_Portrait | 0.04 | 0.17262 | 0.06 | 0.10934 | -0.01 | 0.82726 |
| Rassf8 | 71323 | 312846 | Up | FLX_NoStress_Portrait | 0.02 | 0.69643 | NA | NA | -0.03 | 0.35432 |
| Tspan5 | 56224 | 362048 | Up | FLX_NoStress_Portrait | 0.08 | **0.04457** | 0.13 | 0.08118 | 0.02 | 0.34080 |
| Kif5b | 16573 | 117550 | Up | FLX_NoStress_Portrait | 0.01 | 0.78147 | -0.01 | 0.75502 | 0.02 | 0.29693 |
| Cfh | 12628 | NA | Up | FLX_NoStress_Portrait | 0.05 | 0.24816 | NA | NA | NA | NA |
| Homer1 | 26556 | 29546 | Up | FLX_NoStress_Portrait | 0.07 | 0.26035 | NA | NA | NA | NA |
| Tmem47 | 192216 | 501569 | Up | FLX_NoStress_Portrait | 0.06 | 0.21331 | 0.08 | 0.26519 | 0.01 | 0.70893 |
| Tmem98 | 103743 | 303356 | Up | FLX_NoStress_Portrait | 0.02 | 0.66307 | 0.03 | 0.64104 | 0.06 | 0.41600 |
| Knstrn | 51944 | 311325 | Up | FLX_NoStress_Portrait | 0.07 | **0.00032** | 0.06 | **0.00186** | 0.09 | **0.02808** |
| Sema3a | 20346 | 29751 | Up | FLX_NoStress_Portrait | 0.18 | 0.12547 | 0.24 | 0.23988 | 0.06 | 0.26520 |
| C1qb | 12260 | 29687 | Up | FLX_NoStress_Portrait | 0.03 | 0.57979 | 0.07 | 0.36050 | -0.04 | 0.53511 |
| Dpp4 | 13482 | 25253 | Up | FLX_NoStress_Portrait | 0.07 | 0.36156 | 0.07 | 0.57164 | 0.07 | 0.19000 |
| Igfbp6 | 16012 | 25641 | Up | FLX_NoStress_Portrait | 0.33 | 0.08497 | 0.62 | **0.03730** | 0.04 | 0.38266 |
| Adk | 11534 | 25368 | Up | FLX_NoStress_Portrait | -0.04 | **0.03422** | -0.07 | **0.02544** | -0.01 | 0.65074 |
| Drd1 | 13488 | 24316 | Up | FLX_NoStress_Portrait | 0.14 | 0.43428 | 0.36 | 0.16392 | 0.00 | 0.99000 |

| | | | | | | | | | | |
|---|---|---|---|---|---|---|---|---|---|---|
| ***Tyro3*** | 22174 | 25232 | Up | FLX_NoStress_Portrait | 0.02 | 0.64027 | 0.07 | 0.35860 | 0.01 | 0.69646 |
| ***Sox11*** | 20666 | 84046 | Up | FLX_NoStress_Portrait | 0.16 | 0.10243 | NA | NA | 0.08 | 0.15365 |
| ***Adprm*** | 66358 | 287406 | Down | FLX_Stress_Integration | -0.03 | **0.04326** | -0.04 | **0.01980** | 0.01 | 0.77183 |
| ***Hmgcs1*** | 208715 | 29637 | Down | FLX_Stress_Integration | -0.03 | 0.43587 | -0.03 | 0.57411 | 0.01 | 0.70271 |
| ***Klhl5*** | 71778 | 305351 | Down | FLX_Stress_Integration | -0.04 | 0.08157 | -0.07 | **0.03281** | 0.00 | 0.99199 |
| ***Nfib*** | 18028 | 29227 | Down | FLX_Stress_Integration | 0.03 | 0.41464 | 0.01 | 0.91418 | 0.03 | 0.25973 |
| ***Ppara*** | 19013 | 25747 | Down | FLX_Stress_Integration | -0.06 | **0.00106** | -0.08 | **0.00641** | -0.04 | 0.17488 |
| ***Rps10*** | 67097 | NA | Down | FLX_Stress_Integration | NA | NA | NA | NA | NA | NA |
| ***Wnk1*** | 232341 | 116477 | Down | FLX_Stress_Integration | -0.01 | 0.73246 | -0.04 | **0.03477** | 0.04 | 0.17537 |
| ***Wtap*** | 60532 | 499020 | Down | FLX_Stress_Integration | -0.02 | 0.10285 | -0.02 | 0.22358 | -0.02 | 0.21875 |
| ***Zfhx2*** | 239102 | 305888 | Down | FLX_Stress_Integration | 0.03 | 0.32086 | NA | NA | -0.02 | 0.40066 |
| ***Nr4a1*** | 15370 | 79240 | NA | FLX_Stress_Integration | 0.04 | 0.73323 | 0.10 | 0.63727 | -0.03 | 0.48082 |
| ***Arc*** | 11838 | 54323 | Up | FLX_Stress_Integration | 0.00 | 0.98543 | 0.09 | 0.66816 | -0.05 | 0.14209 |
| ***Ddah1*** | 69219 | 64157 | Up | FLX_Stress_Integration | 0.01 | 0.56370 | 0.03 | 0.42231 | 0.00 | 0.89503 |
| ***Egr1*** | 13653 | 24330 | Up | FLX_Stress_Integration | 0.00 | 0.98450 | -0.04 | 0.67043 | 0.06 | 0.34102 |
| ***Ephb6*** | 13848 | 312275 | Up | FLX_Stress_Integration | 0.13 | 0.11573 | 0.24 | 0.08249 | -0.01 | 0.65132 |
| ***Kcng2*** | 240444 | 307234 | Up | FLX_Stress_Integration | -0.02 | 0.71944 | NA | NA | -0.06 | 0.36321 |
| ***Lzts1*** | 211134 | 266711 | Up | FLX_Stress_Integration | -0.04 | 0.48480 | 0.02 | 0.78728 | -0.02 | 0.55439 |
| ***Mef2d*** | 17261 | 81518 | Up | FLX_Stress_Integration | 0.06 | 0.07054 | 0.10 | **0.01213** | 0.00 | 0.97508 |
| ***Oxtr*** | 18430 | 25342 | Up | FLX_Stress_Integration | -0.03 | 0.76265 | NA | NA | -0.02 | 0.62824 |
| ***Prkar1b*** | 19085 | 25521 | Up | FLX_Stress_Integration | 0.10 | 0.09865 | 0.21 | **0.02617** | -0.01 | 0.44480 |
| ***Rimbp2*** | 231760 | 266780 | Up | FLX_Stress_Integration | 0.08 | 0.15915 | NA | NA | 0.01 | 0.83386 |
| ***Sgsm2*** | 97761 | 303304 | Up | FLX_Stress_Integration | 0.01 | 0.81974 | 0.04 | 0.29981 | -0.04 | 0.39418 |
| ***Ttbk1*** | 106763 | 316229 | Up | FLX_Stress_Integration | 0.01 | 0.61770 | NA | NA | -0.01 | 0.81139 |
| ***Ttyh3*** | 78339 | 304315 | Up | FLX_Stress_Integration | 0.03 | 0.34018 | 0.08 | 0.11828 | -0.04 | 0.30143 |
| ***Hmgcs1*** | 208715 | 29637 | Down | FLX_Stress_Portrait | -0.03 | 0.43587 | -0.03 | 0.57411 | 0.01 | 0.70271 |
| ***Rspo3*** | 72780 | 498997 | Down | FLX_Stress_Portrait | -0.14 | **0.01948** | -0.19 | 0.07585 | -0.09 | 0.08610 |
| ***Cnn3*** | 71994 | 54321 | Down | FLX_Stress_Portrait | -0.02 | 0.63357 | -0.07 | 0.31647 | 0.04 | 0.07512 |
| ***Sh3d19*** | 27059 | 295171 | Down | FLX_Stress_Portrait | -0.09 | 0.20813 | NA | NA | NA | NA |
| ***Ier5*** | 15939 | 498256 | Up | FLX_Stress_Portrait | 0.24 | 0.07591 | 0.39 | 0.08793 | 0.01 | 0.82208 |
| ***Mapk4*** | 225724 | 54268 | Up | FLX_Stress_Portrait | 0.09 | 0.09812 | 0.14 | 0.12880 | 0.02 | 0.37855 |
| ***Diras2*** | 68203 | 291006 | Up | FLX_Stress_Portrait | 0.02 | 0.55915 | 0.07 | 0.20962 | -0.04 | 0.06079 |
| ***Kcnh3*** | 16512 | 27150 | Up | FLX_Stress_Portrait | 0.07 | 0.29369 | 0.15 | 0.14055 | -0.04 | 0.35516 |
| ***Sema7a*** | 20361 | 315711 | Up | FLX_Stress_Portrait | 0.16 | **0.02263** | 0.24 | **0.01560** | 0.00 | 0.94162 |
| ***Mapk9*** | 26420 | 50658 | Up | FLX_Stress_Portrait | 0.05 | **0.02866** | 0.08 | **0.02426** | -0.01 | 0.63351 |
| ***Nptx2*** | 53324 | 288475 | Up | FLX_Stress_Portrait | 0.23 | **0.01316** | 0.39 | **0.01568** | 0.04 | 0.22731 |
| ***Prkar1b*** | 19085 | 25521 | Up | FLX_Stress_Portrait | 0.10 | 0.09865 | 0.21 | **0.02617** | -0.01 | 0.44480 |

***Table S14: Hippocampal antidepressant meta-analysis: Enrichment in gene sets derived from snRNA-seq.*** *This .xlsx file includes two worksheets: 1) The worksheet "GSEA_Results_snRNASeq" provides the full fGSEA results derived from the full hippocampal antidepressant meta-analysis, the exploratory traditional antidepressant meta-analysis, and the exploratory non-traditional antidepressant meta-analysis. Each row represents the results for one gene set, and each column provides the fGSEA statistical output. The results are ordered by p-value, so that the top rows in the worksheet are the gene sets with the smallest p-values. 2) The worksheet "Column Definitions" provides the definitions for the variables present in each column in "GSEA_Results_snRNASeq".*

***Table S15. The full cortical meta-analysis results (15,583 genes, 15,454 stable meta-analysis estimates).*** *This .xlsx file includes two worksheets: 1) The worksheet "metaOutputByPval" provides the full meta-analysis results, with each row representing the results for one gene, and each column providing either gene annotation or meta-analysis statistical output. The results are ordered by p-value, so that the top rows in the worksheet are the genes with the smallest p-values. 2) The worksheet "Column Definitions" provides the definitions for the variables present in each column in "metaOutputByPval".*

***Table S16: Cortical directional fast Gene Set Enrichment Analysis (fGSEA) results (10,579 gene sets).***

*This .xlsx file includes two worksheets: 1) The worksheet "CTX_Directional_GSEA" provides the full fGSEA results, with each row representing the results for one gene set, and each column providing the fGSEA statistical output. The results are ordered by p-value, so that the top rows in the worksheet are the gene sets with the smallest p-values. 2) The worksheet "Column Definitions" provides the definitions for the variables present in each column in "CTX_Directional_GSEA".*

***Table S17: Cortical non-directional fast Gene Set Enrichment Analysis (fGSEA) results (10,579 gene sets).*** *This .xlsx file includes two worksheets: 1) The worksheet "CTX_NonDirectional_GSEA" provides the full fGSEA results, with each row representing the results for one gene set, and each column providing the fGSEA statistical output. The results are ordered by p-value, so that the top rows in the worksheet are the gene sets with the smallest p-values. 2) The worksheet "Column Definitions" provides the definitions for the variables present in each column in "CTX_NonDirectional_GSEA".*

***Table S18. Seven genes showed significant evidence of publication bias when considering the full cortical meta-analysis output (Egger's test FDR<0.05).*** *The Egger's regression test is used to detect funnel plot asymmetry, which is a commonly used indicator of publication bias. Table follows the conventions of **Table S7**. Results for all genes can be found in **Table S15.***

| Gene Symbol | Mouse Entrez Gene ID | AD Log2FC Estimate | AD SE | AD CI_lb | AD CI_ub | # Contrasts | AD Pval | AD FDR | Egger Zstat | Egger Pval | Egger FDR |
|---|---|---|---|---|---|---|---|---|---|---|---|
| Pim1 | 18712 | -0.17 | 0.06 | -0.28 | -0.05 | 14 | 4.16E-03 | 4.51E-01 | -5.04 | 4.62E-07 | 5.28E-03 |
| Cartpt | 27220 | 0.08 | 0.07 | -0.05 | 0.21 | 16 | 2.17E-01 | 8.53E-01 | 4.97 | 6.77E-07 | 5.28E-03 |
| Hspa1a | 193740 | -0.21 | 0.16 | -0.52 | 0.11 | 13 | 2.00E-01 | 8.51E-01 | -4.80 | 1.59E-06 | 8.24E-03 |
| Gpr149 | 229357 | 0.11 | 0.06 | 0.00 | 0.22 | 16 | 6.12E-02 | 7.25E-01 | 4.35 | 1.37E-05 | 4.08E-02 |
| Scg2 | 20254 | 0.01 | 0.05 | -0.09 | 0.12 | 16 | 7.86E-01 | 9.77E-01 | 4.34 | 1.45E-05 | 4.08E-02 |
| Sox1 | 20664 | 0.03 | 0.08 | -0.12 | 0.19 | 13 | 6.71E-01 | 9.51E-01 | 4.32 | 1.57E-05 | 4.08E-02 |
| Vgll2 | 215031 | 0.38 | 0.25 | -0.11 | 0.87 | 11 | 1.27E-01 | 8.14E-01 | 4.27 | 1.99E-05 | 4.44E-02 |

***Table S19. Exploratory cortical results from a meta-regression including antidepressant type (non-traditional vs. traditional) and dissection (PFC vs. other cortex) as co-variates (15,583 genes, 15,406 stable meta-analysis estimates).*** *This .xlsx file includes two worksheets: 1) The worksheet "metaOutputByPval" provides the full meta-analysis results, with each row representing the results for one gene, and each column providing either gene annotation or meta-analysis statistical output. The results are ordered by p-value for the overall effect of antidepressants, so that the top rows in the worksheet are the genes with the smallest p-values. 2) The worksheet "Column Definitions" provides the definitions for the variables present in each column in "metaOutputByPval".*

***Table S20. Exploratory cortical results from a meta-regression including antidepressant type (non-traditional vs. traditional) and dissection (ACg vs. other cortex) as co-variates (15,583 genes, 15,464 stable meta-analysis estimates).*** *This .xlsx file includes two worksheets: 1) The worksheet "metaOutputByPval" provides the full meta-analysis results, with each row representing the results for one gene, and each column providing either gene annotation or meta-analysis statistical output. The results are ordered by p-value for the overall effect of antidepressants, so that the top rows in the worksheet are the genes with the smallest p-values. 2) The worksheet "Column Definitions" provides the definitions for the variables present in each column in "metaOutputByPval".*

***Table S21. Exploratory results from a meta-regression encompassing both cortical and hippocampal datasets, with brain region included as a co-variate (12,857 genes, 12,835 stable meta-analysis estimates).*** *This .xlsx file includes two worksheets: 1) The worksheet "metaOutputByPval" provides the full meta-analysis results, with each row representing the results for one gene, and each column providing either gene annotation or meta-analysis statistical output. The results are ordered by p-value for the overall effect of antidepressants, so that the top rows in the worksheet are the genes with the smallest p-values. 2) The worksheet "Column Definitions" provides the definitions for the variables present in each column in "metaOutputByPval".*

### Supplemental Figures

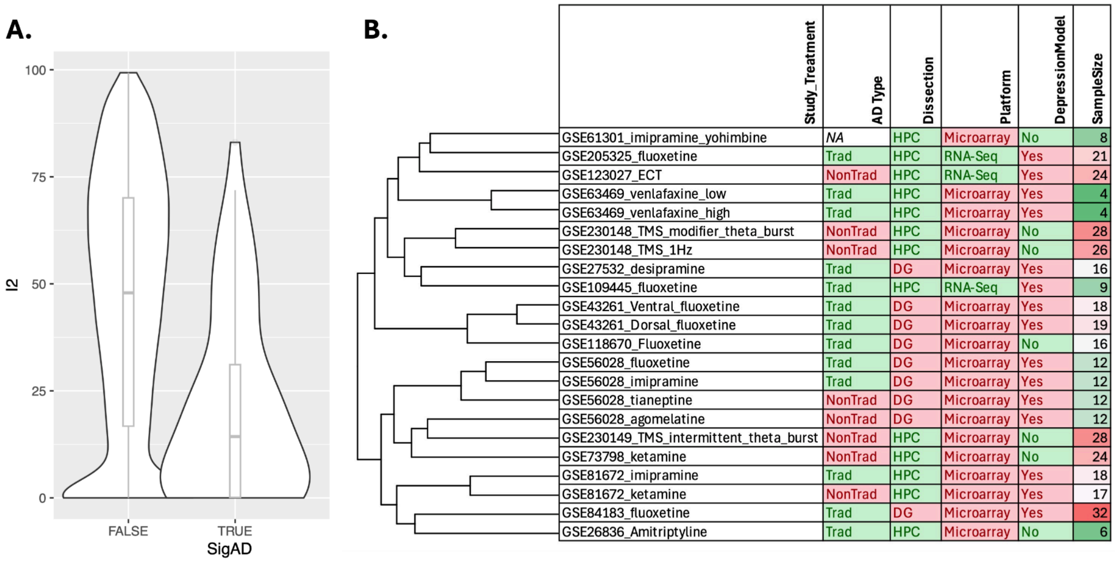


***Figure S1. Exploring heterogeneity in the antidepressant effects across hippocampal studies and contrasts. A.*** *There is evidence of significant heterogeneity within the antidepressant effects calculated across hippocampal studies and contrasts. A violin plot illustrates the distribution of the $I^2$ statistic for genes that were significant antidepressant DEGs in our meta-analysis (58 genes FDR<0.05: "True") or not ("False"). $I^2$ estimates (in percent) how much of the total variability in the observed antidepressant effect sizes (Log2FCs) can be attributed to heterogeneity among the true effects. $I^2$ is smaller for genes that were DEGs in our antidepressant meta-analysis, but still shows wide variation. Full results can be found in* ***Table S2***. ***B.*** *Hierarchical clustering of the antidepressant vs. control contrasts from each of the studies, as performed using the spearman rank correlations of the antidepressant effect sizes (Log2FCs) for all genes that are shared between each pair of studies and that were included in the meta-analysis. The clustering does not easily map on to any of the suspected variables, including antidepressant type (traditional (Trad) vs. non-traditional (NonTrad)), dissection type (dentate gyrus (DG) vs. whole hippocampus (HPC), transcriptional profiling platform, whether the study included any form of animal depression model, or the sample size associated with the contrast (n=antidepressant + control).*

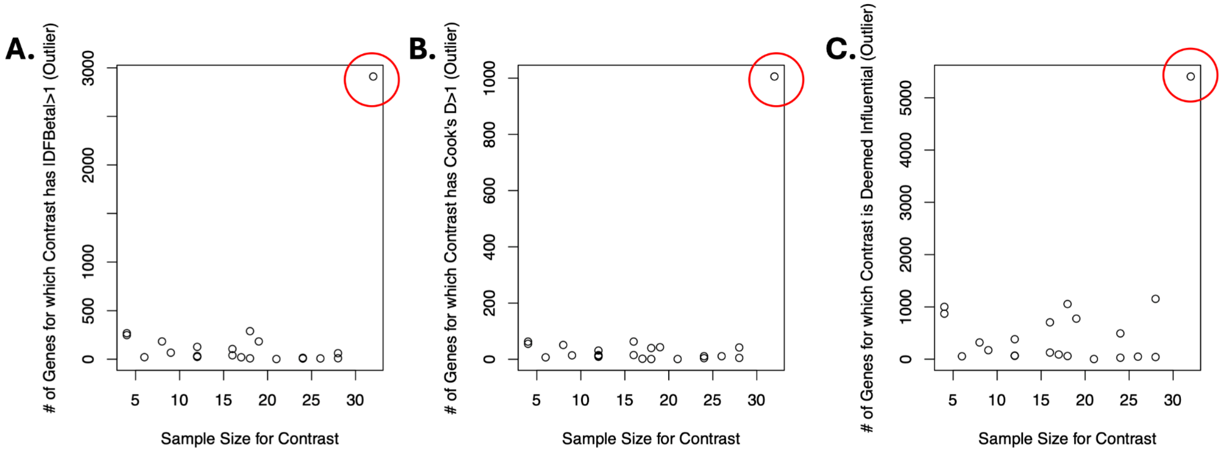


***Figure S2. Small sample size studies do not appear to be adding disproportionately to the noise in our hippocampal meta-analysis.*** *When examining the hippocampal meta-analysis results for any particular gene, the study that seemed to be the most likely to be flagged as having an antidepressant effect (Log2FC) that was distinctly different from the group estimate, causing disproportionate influence (i.e., outlier status), was actually the study with the largest sample size (GSE84183_Fluoxetine). However, this study did not stand out as an extreme outlier in our hierarchical clustering analyses, and it does not have any features that would suggest it should be removed from the analysis.* ***A.*** *A scatterplot showing the number of genes that had a particular antidepressant vs. control contrast flagged as an outlier using one common definition of influence (Difference in Betas (DFBetas)>1) in relationship to the sample size for that contrast (n=antidepressant+control).* ***B.*** *A scatterplot showing the number of genes that had a particular antidepressant vs. control contrast flagged as an outlier using one common definition of influence (Cook's difference (Cook's d)>1) in relationship to the sample size for the contrast (n=antidepressant+control).* ***C.*** *A scatterplot showing the number of genes that had a particular antidepressant vs. control contrast flagged as an outlier using the influence() function provided by the metafor package in relationship to the sample size for the contrast (n=antidepressant+control).*

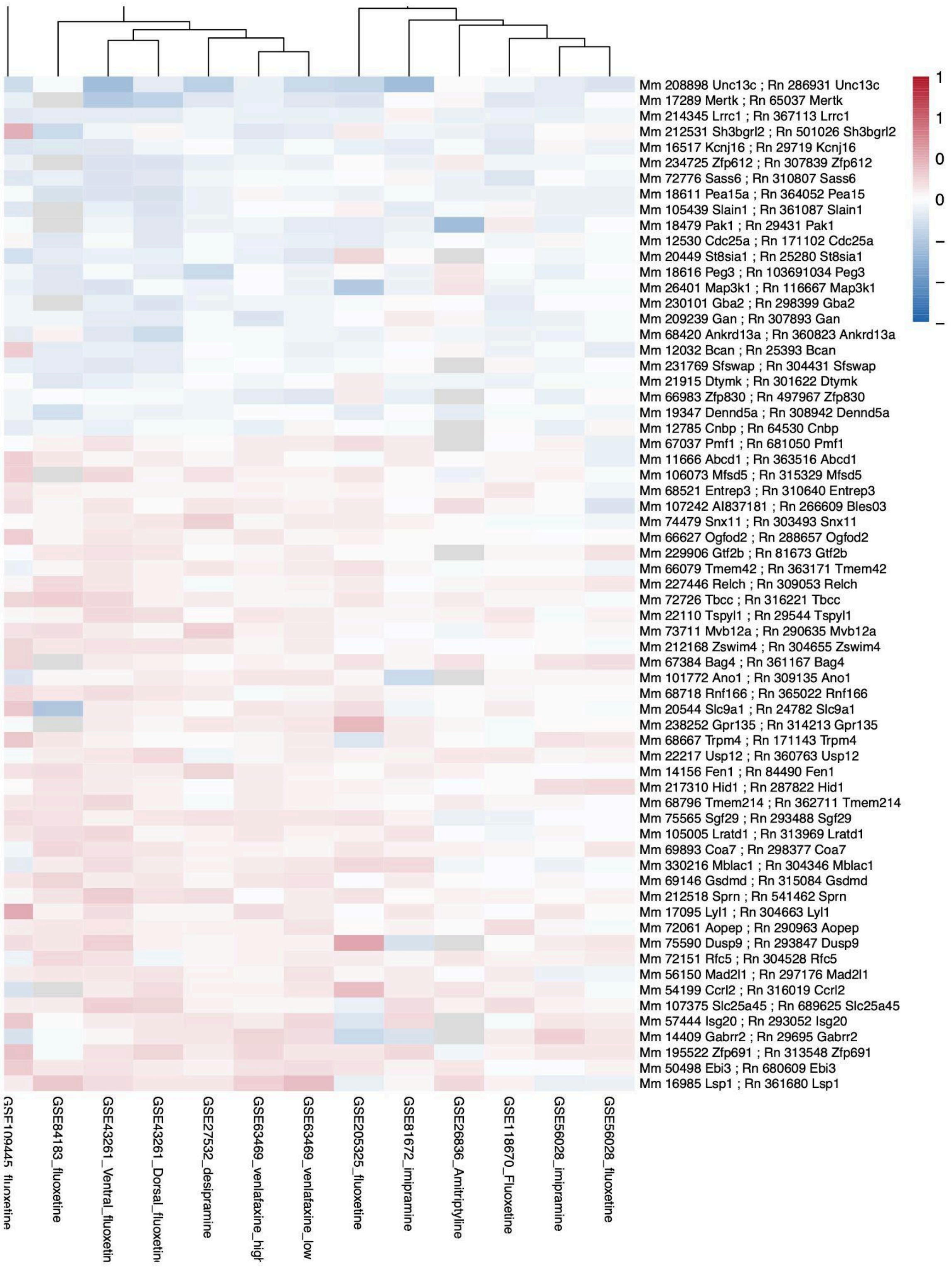

Mm 208898 Unc13c ; Rn 286931 Unc13c
Mm 17289 Mertk ; Rn 65037 Mertk
Mm 214345 Lrrc1 ; Rn 367113 Lrrc1
Mm 212531 Sh3bgrl2 ; Rn 501026 Sh3bgrl2
Mm 16517 Kcnj16 ; Rn 29719 Kcnj16
Mm 234725 Zfp612 ; Rn 307839 Zfp612
Mm 72776 Sass6 ; Rn 310807 Sass6
Mm 18611 Pea15a ; Rn 364052 Pea15
Mm 105439 Slain1 ; Rn 361087 Slain1
Mm 18479 Pak1 ; Rn 29431 Pak1
Mm 12530 Cdc25a ; Rn 171102 Cdc25a
Mm 20449 St8sia1 ; Rn 25280 St8sia1
Mm 18616 Peg3 ; Rn 103691034 Peg3
Mm 26401 Map3k1 ; Rn 116667 Map3k1
Mm 230101 Gba2 ; Rn 298399 Gba2
Mm 209239 Gan ; Rn 307893 Gan
Mm 68420 Ankrd13a ; Rn 360823 Ankrd13a
Mm 12032 Bcan ; Rn 25393 Bcan
Mm 231769 Sfswap ; Rn 304431 Sfswap
Mm 21915 Dtymk ; Rn 301622 Dtymk
Mm 66983 Zfp830 ; Rn 497967 Zfp830
Mm 19347 Dennd5a ; Rn 308942 Dennd5a
Mm 12785 Cnbp ; Rn 64530 Cnbp
Mm 67037 Pmf1 ; Rn 681050 Pmf1
Mm 11666 Abcd1 ; Rn 363516 Abcd1
Mm 106073 Mfsd5 ; Rn 315329 Mfsd5
Mm 68521 Entrep3 ; Rn 310640 Entrep3
Mm 107242 AI837181 ; Rn 266609 Bles03
Mm 74479 Snx11 ; Rn 303493 Snx11
Mm 66627 Ogfod2 ; Rn 288657 Ogfod2
Mm 229906 Gtf2b ; Rn 81673 Gtf2b
Mm 66079 Tmem42 ; Rn 363171 Tmem42
Mm 227446 Relch ; Rn 309053 Relch
Mm 72726 Tbcc ; Rn 316221 Tbcc
Mm 22110 Tspyl1 ; Rn 29544 Tspyl1
Mm 73711 Mvb12a ; Rn 290635 Mvb12a
Mm 212168 Zswim4 ; Rn 304655 Zswim4
Mm 67384 Bag4 ; Rn 361167 Bag4
Mm 101772 Ano1 ; Rn 309135 Ano1
Mm 68718 Rnf166 ; Rn 365022 Rnf166
Mm 20544 Slc9a1 ; Rn 24782 Slc9a1
Mm 238252 Gpr135 ; Rn 314213 Gpr135
Mm 68667 Trpm4 ; Rn 171143 Trpm4
Mm 22217 Usp12 ; Rn 360763 Usp12
Mm 14156 Fen1 ; Rn 84490 Fen1
Mm 217310 Hid1 ; Rn 287822 Hid1
Mm 68796 Tmem214 ; Rn 362711 Tmem214
Mm 75565 Sgf29 ; Rn 293488 Sgf29
Mm 105005 Lratd1 ; Rn 313969 Lratd1
Mm 69893 Coa7 ; Rn 298377 Coa7
Mm 330216 Mblac1 ; Rn 304346 Mblac1
Mm 69146 Gsdmd ; Rn 315084 Gsdmd
Mm 212518 Sprn ; Rn 541462 Sprn
Mm 17095 Lyl1 ; Rn 304663 Lyl1
Mm 72061 Aopep ; Rn 290963 Aopep
Mm 75590 Dusp9 ; Rn 293847 Dusp9
Mm 72151 Rfc5 ; Rn 304528 Rfc5
Mm 56150 Mad2l1 ; Rn 297176 Mad2l1
Mm 54199 Ccrl2 ; Rn 316019 Ccrl2
Mm 107375 Slc25a45 ; Rn 689625 Slc25a45
Mm 57444 Isg20 ; Rn 293052 Isg20
Mm 14409 Gabrr2 ; Rn 29695 Gabrr2
Mm 195522 Zfp691 ; Rn 313548 Zfp691
Mm 50498 Ebi3 ; Rn 680609 Ebi3
Mm 16985 Lsp1 ; Rn 361680 Lsp1
GSE109445_fluoxetine
GSE84183_fluoxetine
GSE43261_Ventral_fluoxetin
GSE43261_Dorsal_fluoxetin
GSE27532_desipramine
GSE63469_venlafaxine_high
GSE63469_venlafaxine_low
GSE205325_fluoxetine
GSE81672_imipramine
GSE26836_Amitriptyline
GSE118670_Fluoxetine
GSE56028_imipramine
GSE56028_fluoxetine
1
1
0
0

***Figure S3. Heatmap of the Top 50 Traditional Antidepressant Hippocampal Meta-Analysis Genes Across Datasets.*** *Each column represents an individual dataset and each row represents one gene. The color scale indicates antidepressant vs. control effect size ($log_2$ fold change), with red denoting upregulation and blue denoting downregulation relative to control samples. Heatmaps allow for visual comparison of the expression patterns of the top 50 meta-analysis genes across individual datasets. The dendrogram at the top groups datasets by similarity in their transcriptional profiles, with shorter connecting lines indicating greater similarity in gene expression changes across datasets. The full results for the hippocampal non-traditional antidepressant meta-analysis can be found in* ***Table S8****.*

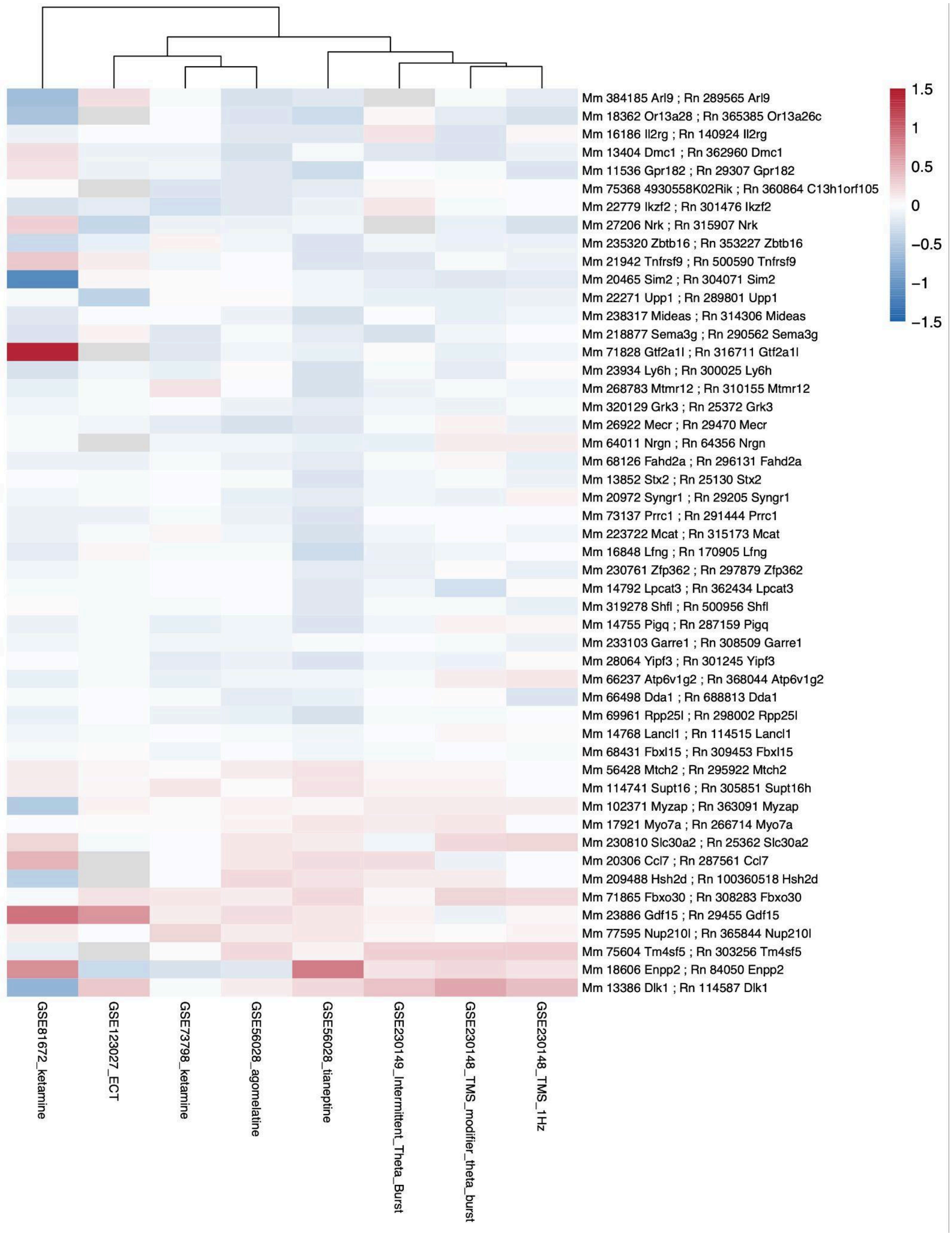

1.5
1
0.5
0
−0.5
−1
−1.5
Mm 384185 Arl9 ; Rn 289565 Arl9
Mm 18362 Or13a28 ; Rn 365385 Or13a26c
Mm 16186 Il2rg ; Rn 140924 Il2rg
Mm 13404 Dmc1 ; Rn 362960 Dmc1
Mm 11536 Gpr182 ; Rn 29307 Gpr182
Mm 75368 4930558K02Rik ; Rn 360864 C13h1orf105
Mm 22779 Ikzf2 ; Rn 301476 Ikzf2
Mm 27206 Nrk ; Rn 315907 Nrk
Mm 235320 Zbtb16 ; Rn 353227 Zbtb16
Mm 21942 Tnfrsf9 ; Rn 500590 Tnfrsf9
Mm 20465 Sim2 ; Rn 304071 Sim2
Mm 22271 Upp1 ; Rn 289801 Upp1
Mm 238317 Mideas ; Rn 314306 Mideas
Mm 218877 Sema3g ; Rn 290562 Sema3g
Mm 71828 Gtf2a1l ; Rn 316711 Gtf2a1l
Mm 23934 Ly6h ; Rn 300025 Ly6h
Mm 268783 Mtmr12 ; Rn 310155 Mtmr12
Mm 320129 Grk3 ; Rn 25372 Grk3
Mm 26922 Mecr ; Rn 29470 Mecr
Mm 64011 Nrgn ; Rn 64356 Nrgn
Mm 68126 Fahd2a ; Rn 296131 Fahd2a
Mm 13852 Stx2 ; Rn 25130 Stx2
Mm 20972 Syngr1 ; Rn 29205 Syngr1
Mm 73137 Prrc1 ; Rn 291444 Prrc1
Mm 223722 Mcat ; Rn 315173 Mcat
Mm 16848 Lfng ; Rn 170905 Lfng
Mm 230761 Zfp362 ; Rn 297879 Zfp362
Mm 14792 Lpcat3 ; Rn 362434 Lpcat3
Mm 319278 Shfl ; Rn 500956 Shfl
Mm 14755 Pigq ; Rn 287159 Pigq
Mm 233103 Garre1 ; Rn 308509 Garre1
Mm 28064 Yipf3 ; Rn 301245 Yipf3
Mm 66237 Atp6v1g2 ; Rn 368044 Atp6v1g2
Mm 66498 Dda1 ; Rn 688813 Dda1
Mm 69961 Rpp25l ; Rn 298002 Rpp25l
Mm 14768 Lancl1 ; Rn 114515 Lancl1
Mm 68431 Fbxl15 ; Rn 309453 Fbxl15
Mm 56428 Mtch2 ; Rn 295922 Mtch2
Mm 114741 Supt16 ; Rn 305851 Supt16h
Mm 102371 Myzap ; Rn 363091 Myzap
Mm 17921 Myo7a ; Rn 266714 Myo7a
Mm 230810 Slc30a2 ; Rn 25362 Slc30a2
Mm 20306 Ccl7 ; Rn 287561 Ccl7
Mm 209488 Hsh2d ; Rn 100360518 Hsh2d
Mm 71865 Fbxo30 ; Rn 308283 Fbxo30
Mm 23886 Gdf15 ; Rn 29455 Gdf15
Mm 77595 Nup210l ; Rn 365844 Nup210l
Mm 75604 Tm4sf5 ; Rn 303256 Tm4sf5
Mm 18606 Enpp2 ; Rn 84050 Enpp2
Mm 13386 Dlk1 ; Rn 114587 Dlk1
GSE81672_ketamine
GSE123027_ECT
GSE73798_ketamine
GSE56028_agomelatine
GSE56028_tianeptine
GSE230149_Intermittent_Theta_Burst
GSE230148_TMS_modifier_theta_burst
GSE230148_TMS_1Hz

***Figure S4. Heatmap of the Top 50 Non-traditional Antidepressant Hippocampal Meta-Analysis Genes Across Datasets.*** *Each column represents an individual dataset and each row represents one gene. The color scale indicates antidepressant vs. control effect size (log$_2$ fold change), with red denoting upregulation and blue denoting downregulation relative to control samples. Heatmaps allow for visual comparison of the expression patterns of the top 50 meta-analysis genes across individual datasets. The dendrogram at the top groups datasets by similarity in their transcriptional profiles, with shorter connecting lines indicating greater similarity in gene expression changes across datasets. The full results for the hippocampal non-traditional antidepressant meta-analysis can be found in* ***Table S9****.*

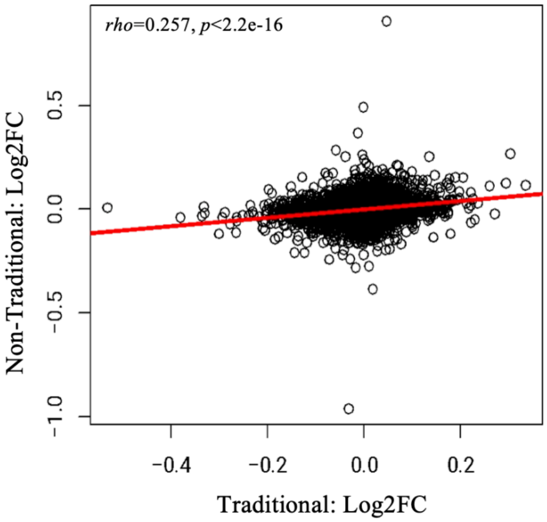


***Figure S5. Exploratory analysis: Traditional and non-traditional antidepressants have overlapping effects on the hippocampus.*** *Scatterplot comparing antidepressant vs. control effects (log$_2$ fold changes) derived from exploratory meta-analyses examining the effects of traditional (n=177, 13 contrasts) and non-traditional (n=148, 8 contrasts) antidepressants on the hippocampus. The two types of antidepressants show a moderate positive correlation in their effects when examining all genes included in both meta-analyses (df=12,614, rho=0.257, p<2.2e-16 ).*

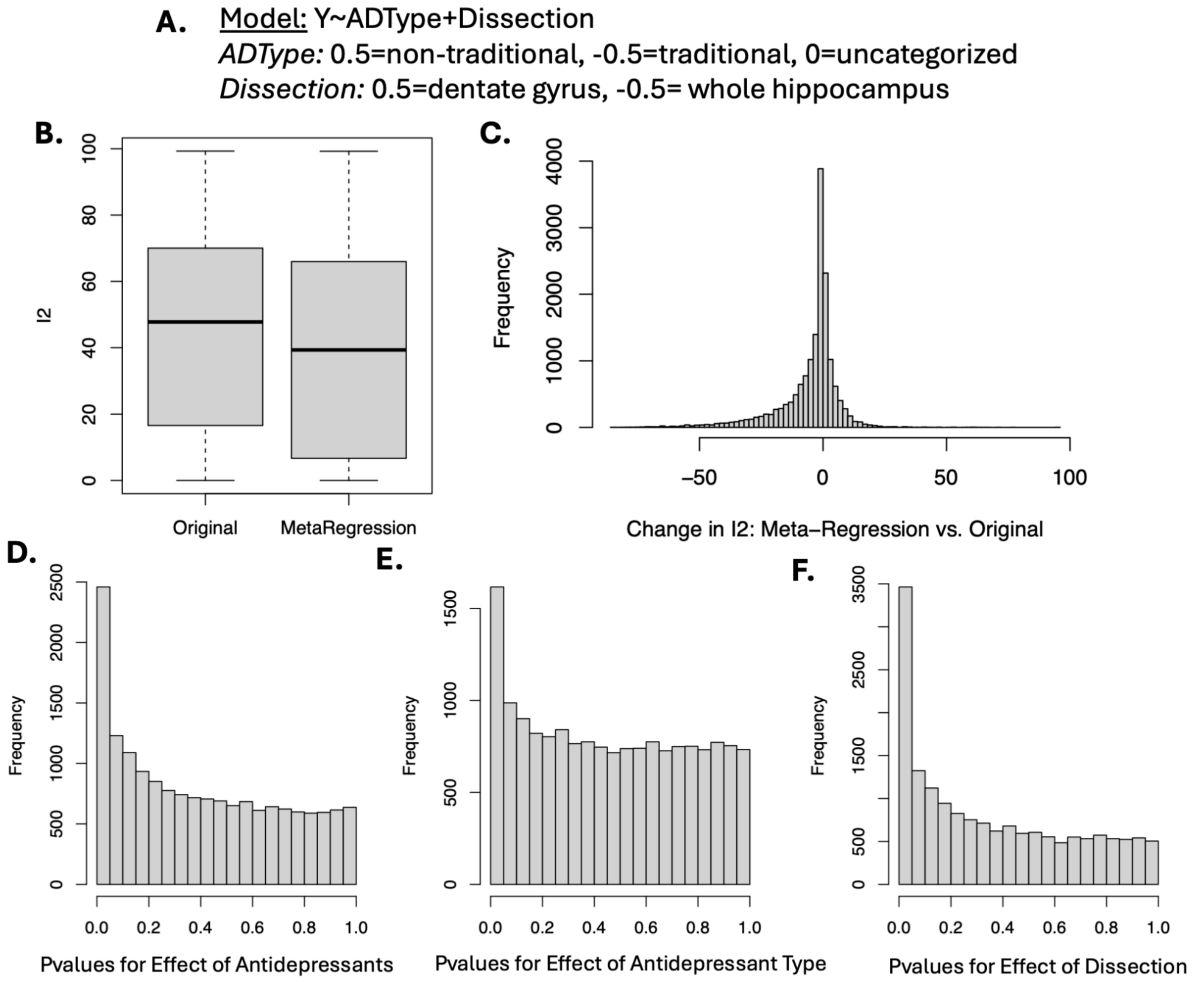


***Figure S6. A meta-regression including antidepressant type (non-traditional vs. traditional) and dissection (dentate gyrus (DG) vs. whole hippocampus) as co-variates provided insight into the heterogeneity in antidepressant effects observed in the hippocampal data. A.*** *The meta-regression model.* ***B-C.*** *Meta-regression decreased the residual heterogeneity present in the effects, as illustrated by a boxplot showing the $I^2$ values for all genes included in the original meta-analysis or meta-regression (**B**) or by a histogram illustrating the change in $I^2$ between the original meta-analysis and meta-regression (**C**).* ***D-F.*** *Histograms showing that the p-values for each of the variables in the meta-regression are enriched in favor of significant findings:* ***D.*** *Overall effect of antidepressants,* ***E.*** *Effect of antidepressant type (non-traditional vs. traditional),* ***F.*** *Effect of dissection (dentate gyrus (DG) vs. whole hippocampus)). The full results can be found in* ***Table S10.***

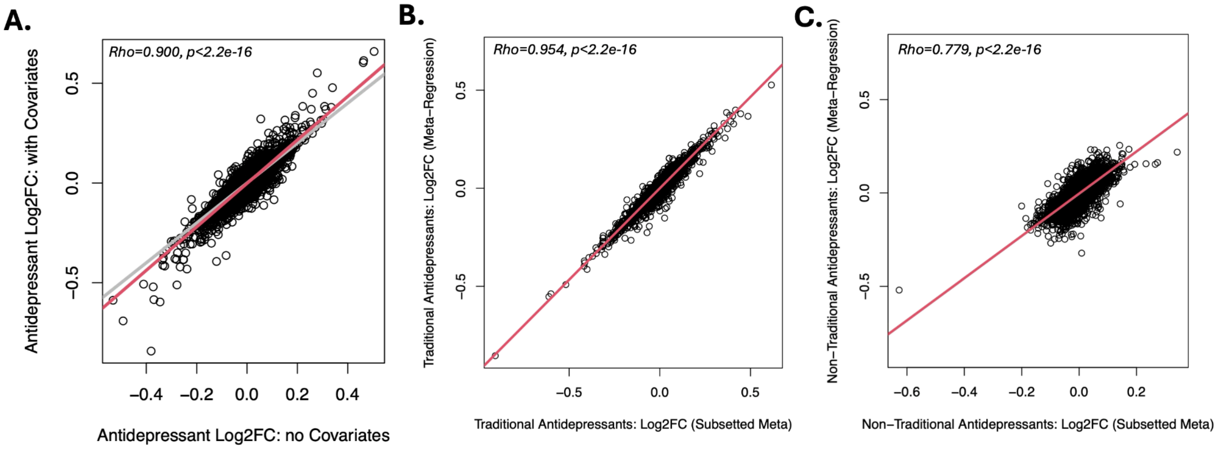


***Figure S7. A meta-regression including antidepressant type (non-traditional vs. traditional) and dissection (dentate gyrus (DG) vs. whole hippocampus) as co-variates produced estimates of antidepressant effects that closely resembled the estimates from our original meta-analyses. A.*** *A scatterplot shows that the effects of antidepressants (Log2FCs) derived from the original meta-analysis (no covariates) closely correlates with the effects of antidepressants (Log2FCs) derived from our meta-regression (covariates: antidepressant type, dissection) (Rho=0.900, p<2.2e-16), with a slope (red) closely approximating 1 (grey).* ***B.*** *A scatterplot shows that the effects of antidepressants (Log2FCs) derived from our subgroup meta-analysis focused on traditional antidepressants (no covariates) closely correlates with the predicted effects of traditional antidepressants (Log2FCs) derived from our meta-regression (-0.5* Log2FC for antidepressant type+Log2FC for overall effect of antidepressants) (Rho=0.954, p<2.2e-16).* ***C.*** *A scatterplot shows that the effects of antidepressants (Log2FCs) derived from our subgroup meta-analysis focused on non-traditional antidepressants (no covariates) correlates with the predicted effects of non-traditional antidepressants (Log2FCs) derived from our meta-regression (0.5* Log2FC for antidepressant type+Log2FC for overall effect of antidepressants) (Rho=0.779, p<2.2e-16).*

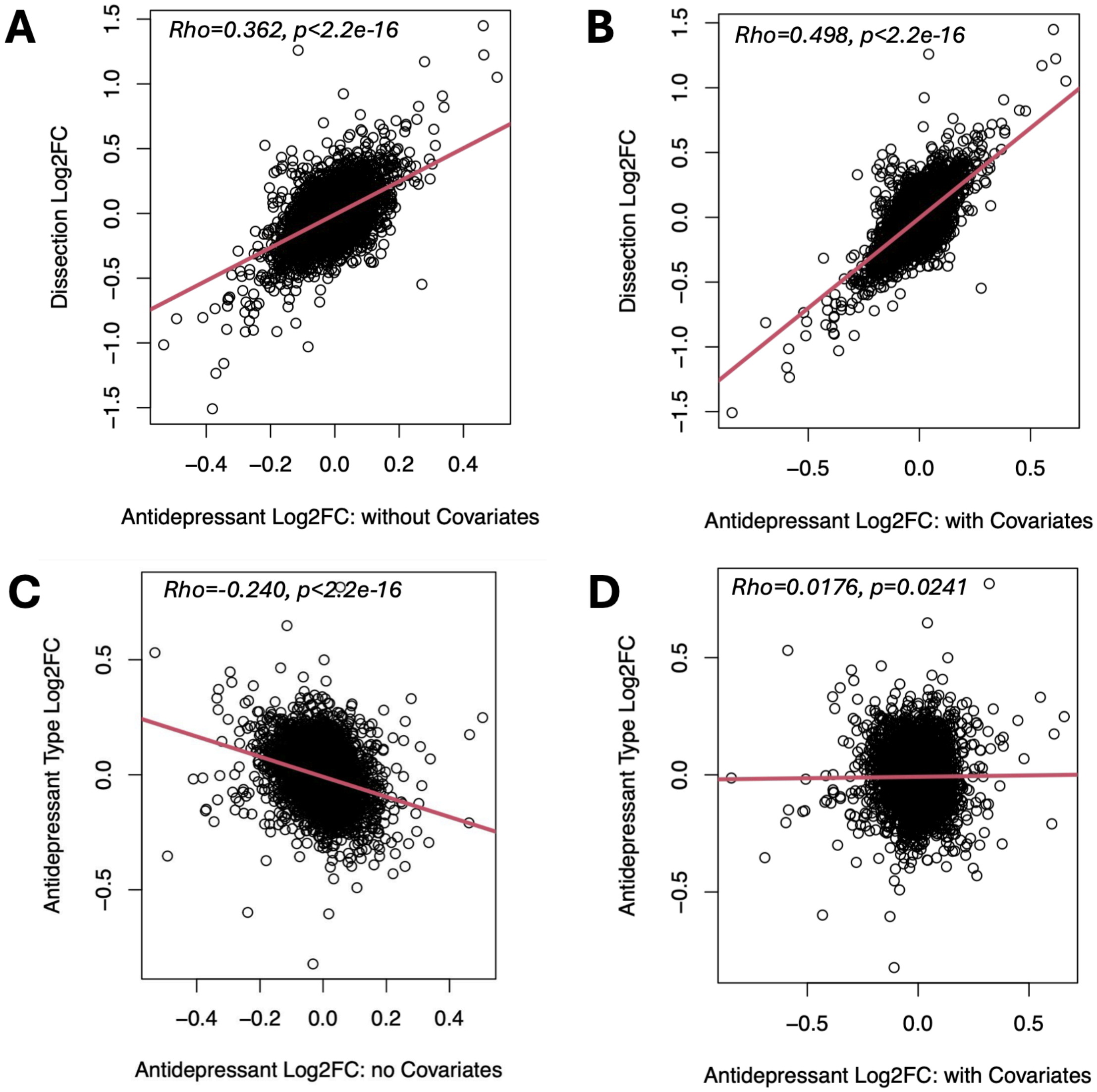


***Figure S8. A meta-regression including antidepressant type (non-traditional vs. traditional) and dissection (dentate gyrus (DG) vs. whole hippocampus) as co-variates suggests that antidepressant effects are larger in the dentate gyrus. A.*** *A scatterplot shows that the effects of dissection (Log2FCs) derived from our meta-regression (dentate gyrus (DG) vs. whole hippocampus) correlates positively with the effects of antidepressants (Log2FCs) derived from the original meta-analysis (no covariates) (Rho=0.362, p<2.2e-16).* ***B.*** *A scatterplot shows that the effects of dissection (Log2FCs) derived from our meta-regression (dentate gyrus (DG) vs. whole hippocampus) correlates positively with the overall effects of antidepressants (Log2FCs) derived from the meta-regression (Rho=0.498, p<2.2e-16).* ***C.*** *A scatterplot shows that the effects of antidepressant type (Log2FCs) derived from our meta-regression (non-traditional vs. traditional) correlates negatively with the effects of antidepressants (Log2FCs) derived from the original meta-analysis (no covariates) (Rho=-0.240, p<2.2e-16), implying that the original meta-analysis was slightly skewed in favor of traditional antidepressants.* ***D.*** *A scatterplot shows that the effects of antidepressant type (Log2FCs) derived from our meta-regression (non-traditional vs.*

*traditional) shows extremely minimal correlation with the overall effects of antidepressants (Log2FCs) derived from the meta-regression (Rho=0.0176, p=0.0241).*

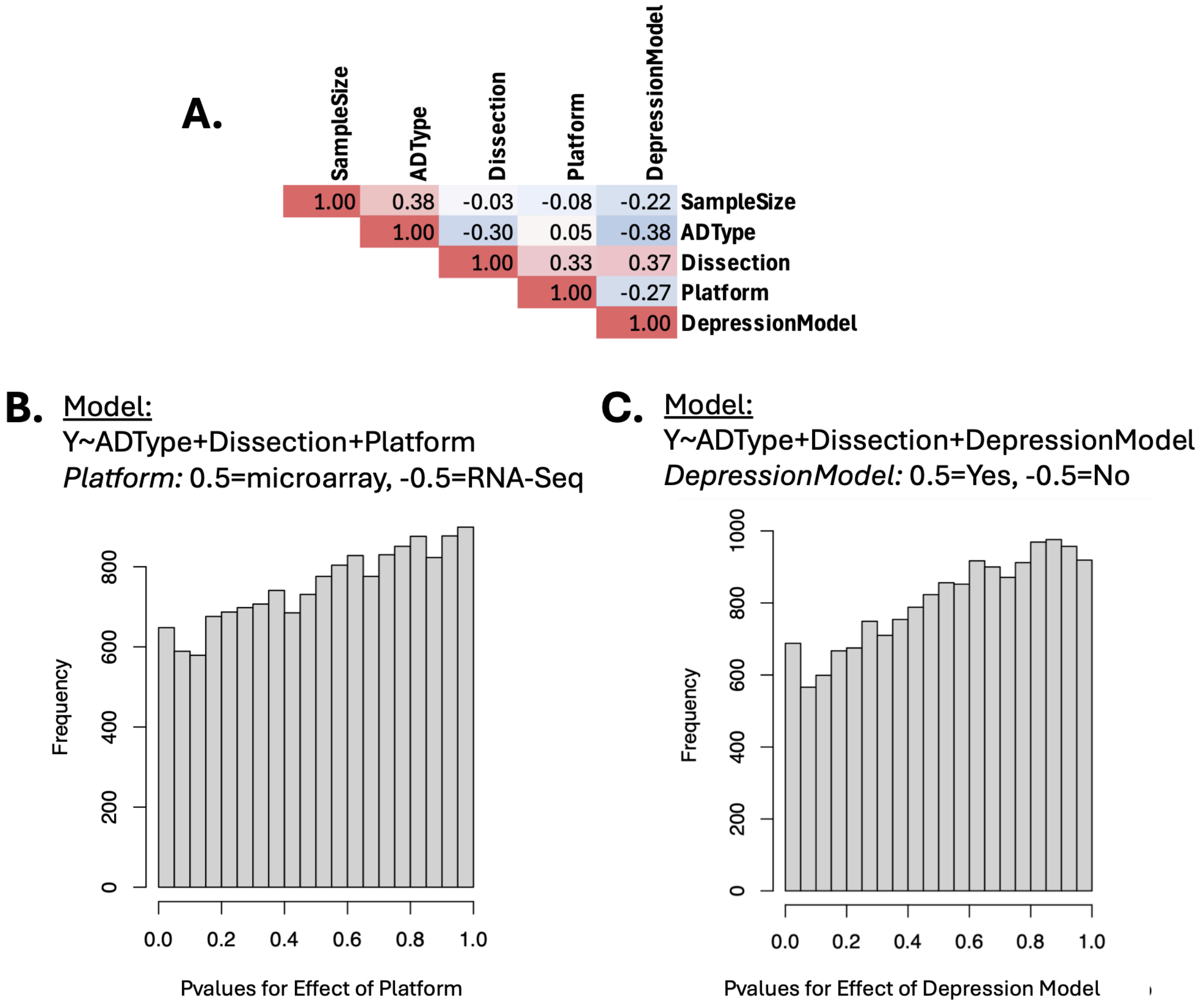


***Figure S9. Other potential hippocampal meta-regression models produced less insight into the heterogeneity in antidepressant effects observed in the hippocampal data. A.*** *When constructing the meta-regression model, it was important to consider the fact that many of the potential co-variates of interest were correlated with each other, as illustrated here by a spearman's rank correlation matrix. The co-variates that were considered are sample size for the contrast (n=antidepressant+control), antidepressant type (ADType: non-traditional vs. traditional), dissection (dentate gyrus (DG) vs. whole hippocampus), platform (microarray vs. RNA-seq), and whether the study included a depression model (vs. controls only).* ***B.*** *Transcriptional profiling platform (microarray vs. RNA-Seq) was a potential variable of interest for strong theoretical reasons, but it was not well represented in the data (only 3 RNA-Seq study contrasts total, and potentially less for any particular gene). When included in an exploratory meta-regression, it showed no significant modulating effect, potentially due to being underpowered, and its inclusion in the model led to the exclusion of 1,413 genes from the analysis which were only represented on one platform. A histogram shows that the p-values for Platform in the meta-regression showed no enrichment in favor of significant findings.* ***C.*** *Whether a study included a*

*depression model (vs. only control subjects) was a potential variable of interest for strong theoretical reasons, but our study was not designed well to quantify it - we extracted antidepressant effects for all subjects in a study that received the same treatment as a single input for the meta-analysis model (Log2FC & accompanying sampling variance). Therefore, it was not surprising that when this variable was included in an exploratory meta-regression, it showed only one gene with a significant modulating effect ($FDR<0.05$: Zpbp2). A histogram shows that the p-values for depression model in the meta-regression showed no enrichment in favor of significant findings. For a meta-analysis study that is better designed to tackle this question, we refer the reader to Ibrahim et al. 2022.*

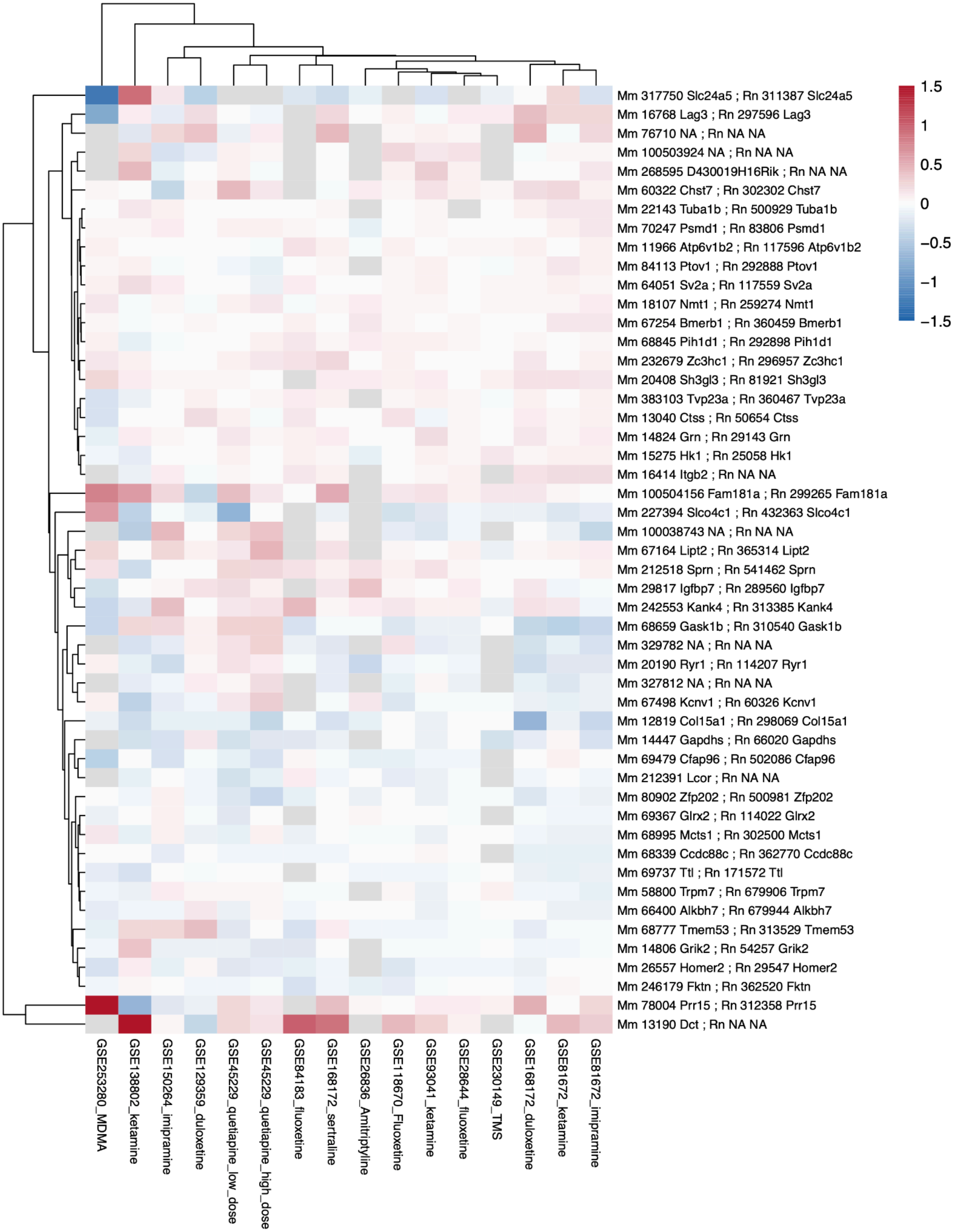


***Figure S10. Heatmap of the Top 50 Cortical Meta-Analysis Genes Across Antidepressant Datasets.*** *Each column represents an individual dataset and each row represents one gene. The color scale*

*indicates the antidepressant vs. control effect size (log₂ fold change), with red denoting upregulation and blue denoting downregulation relative to control samples. Heatmaps allow for visual comparison of the expression patterns of the top 50 meta-analysis genes across individual datasets. The dendrogram at the top groups datasets by similarity in their transcriptional profiles, with shorter connecting lines indicating greater similarity in gene expression changes across datasets. The full results for the cortical antidepressant meta-analysis can be found in* ***Table S15****.*

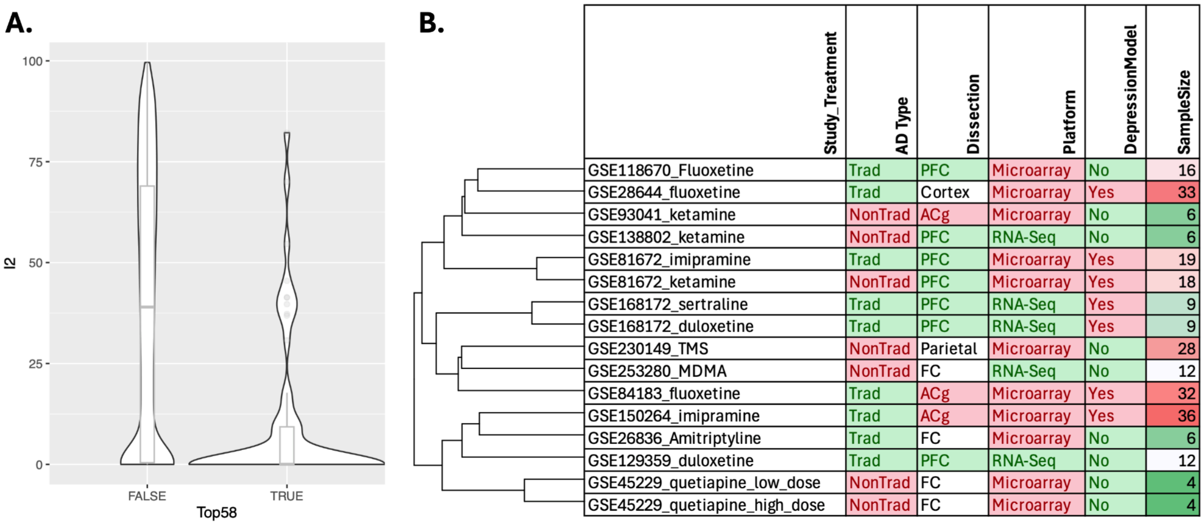


| Study_Treatment | AD Type | Dissection | Platform | DepressionModel | SampleSize |
|---|---|---|---|---|---|
| GSE118670_Fluoxetine | Trad | PFC | Microarray | No | 16 |
| GSE28644_fluoxetine | Trad | Cortex | Microarray | Yes | 33 |
| GSE93041_ketamine | NonTrad | ACg | Microarray | No | 6 |
| GSE138802_ketamine | NonTrad | PFC | RNA-Seq | No | 6 |
| GSE81672_imipramine | Trad | PFC | Microarray | Yes | 19 |
| GSE81672_ketamine | NonTrad | PFC | Microarray | Yes | 18 |
| GSE168172_sertraline | Trad | PFC | RNA-Seq | Yes | 9 |
| GSE168172_duloxetine | Trad | PFC | RNA-Seq | Yes | 9 |
| GSE230149_TMS | NonTrad | Parietal | Microarray | No | 28 |
| GSE253280_MDMA | NonTrad | FC | RNA-Seq | No | 12 |
| GSE84183_fluoxetine | Trad | ACg | Microarray | Yes | 32 |
| GSE150264_imipramine | Trad | ACg | Microarray | Yes | 36 |
| GSE26836_Amitriptyline | Trad | FC | Microarray | No | 6 |
| GSE129359_duloxetine | Trad | PFC | RNA-Seq | No | 12 |
| GSE45229_quetiapine_low_dose | NonTrad | FC | Microarray | No | 4 |
| GSE45229_quetiapine_high_dose | NonTrad | FC | Microarray | No | 4 |

***Figure S11. Exploring heterogeneity in the antidepressant effects across cortical studies and contrasts. A.*** *There is evidence of significant heterogeneity within the antidepressant effects calculated across hippocampal studies and contrasts. A violin plot illustrates the distribution of the $I^2$ statistic for genes that had the highest ranking antidepressant p-values in our cortical meta-analysis (58 genes, to parallel the hippocampal figure: "True") or not ("False"). $I^2$ estimates (in percent) how much of the total variability in the observed antidepressant effect sizes (Log2FCs) can be attributed to heterogeneity among the true effects. $I^2$ is smaller for genes that had high ranking antidepressant p-values in our meta-analysis, but still shows wide variation. The full results can be found in* ***Table S15. B.*** *Hierarchical clustering of the antidepressant vs. control contrasts from each of the studies, as performed based on the spearman rank correlations for the antidepressant vs. control effect sizes (Log2FCs) using all genes shared between each pair of studies that were included in the meta-analysis. The clustering does not easily map on to any of the suspected variables, including antidepressant type (traditional (Trad) vs. non-traditional (NonTrad)), dissection type (anterior cingulate (ACg), prefrontal cortex (PFC), frontal cortex (FC), parietal cortex or cerebral cortex not otherwise defined), transcriptional profiling platform, whether the study included any form of animal depression model (vs. controls only), or the sample size associated with the contrast (n=antidepressant + control).*

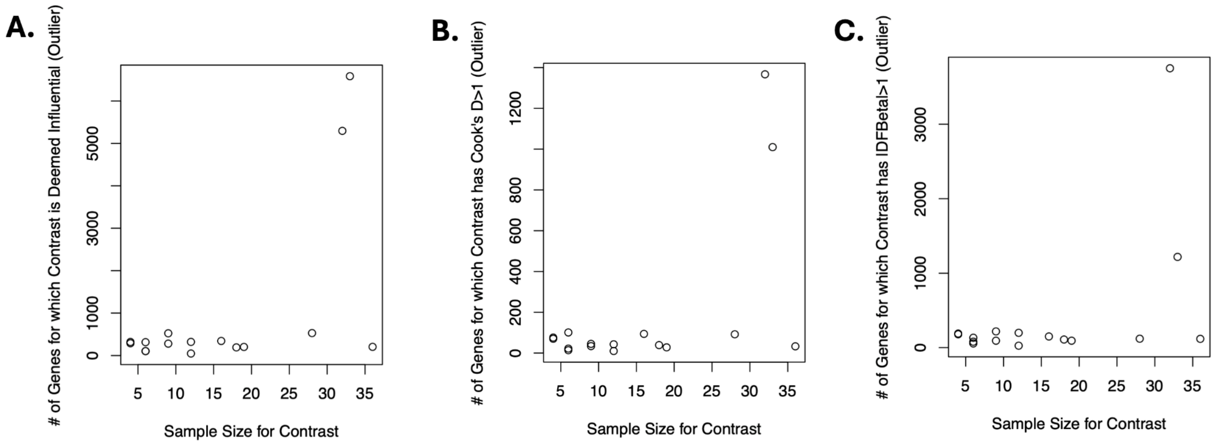


***Figure S12. Small sample size studies do not appear to be adding disproportionately to the noise in our cortical meta-analysis.*** *When examining the meta-analysis results, the two studies that seemed to be the most likely to be flagged as having an antidepressant effect (Log2FC) that was distinctly different from the group estimate, causing disproportionate influence (i.e., outlier status), were actually two of the studies with the largest sample sizes.* ***A.*** *A scatterplot showing the number of genes that had a particular antidepressant vs. control contrast flagged as an outlier using one common definition of influence (Difference in Betas (DFBetas)>1) in relationship to the sample size for that contrast (n=antidepressant+control).* ***B.*** *A scatterplot showing the number of genes that had a particular antidepressant vs. control contrast flagged as an outlier using one common definition of influence (Cook's difference (Cook's d)>1) in relationship to the sample size for the contrast (n=antidepressant+control).* ***C.*** *A scatterplot showing the number of genes that had a particular antidepressant vs. control contrast flagged as an outlier using the influence() function provided by the metafor package versus the sample size for the contrast (n=antidepressant+control).*

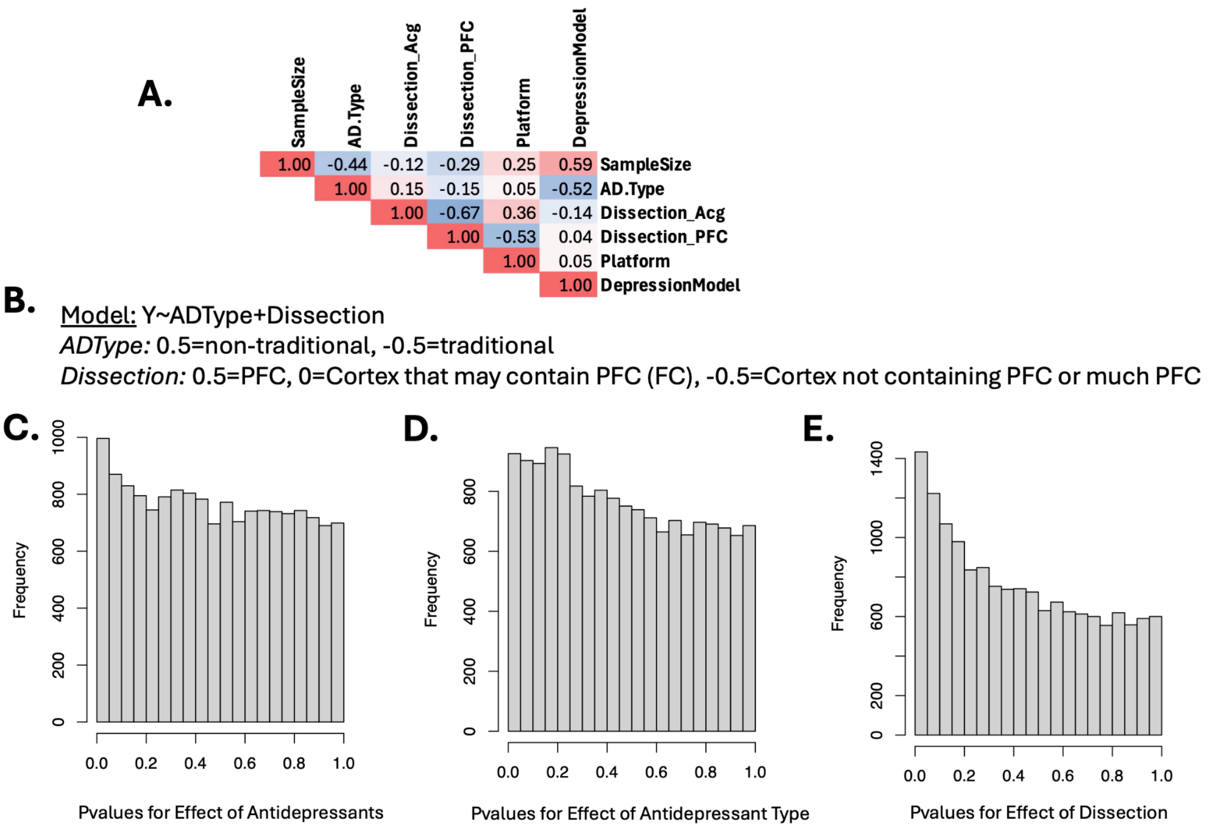


***Figure S13. A cortical meta-regression model produced minimal insight into the heterogeneity in antidepressant effects observed in the cortical data. A.*** *When constructing the meta-regression model, it was important to consider the fact that many of the potential co-variates of interest were correlated with each other, as illustrated here by a spearman's rank correlation matrix. The co-variates that were considered are sample size for the contrast (n=antidepressant+control), antidepressant type (ADType: non-traditional vs. traditional), dissection (either defined as 1: anterior cingulate (ACg) vs. cortex that might contain ACg (FC: frontal cortex) vs. other cortex, or 2: prefrontal cortex (PFC), cortex that might contain prefrontal cortex (FC: frontal cortex) vs. other cortex), platform (microarray vs. RNA-seq), and whether the study included a depression model (vs. controls only).* ***B.*** *Another potential way of summarizing the heterogeneity in dissection in the cortical data was prefrontal cortex (PFC) vs. cortex that might contain PFC (FC: frontal cortex) vs. other cortex.* ***C-D.*** *When this definition was used in a meta-regression model (vs. the definition based on anterior cingulate (ACg) vs. other cortex in* ***Figure S13****), the results for all variables were generally weaker.* ***C.*** *A histogram shows that the p-values for the overall effect of antidepressants in the meta-regression are more weakly enriched in favor of significant findings than in* ***Figure S13****. No genes reached significance for effects (FDR<0.05).* ***D.*** *A histogram shows that the p-values for the effects of antidepressant type in the meta-regression are more weakly enriched in favor of significant findings than in* ***Figure S13****. Seven genes reached significance for effects (FDR<0.05).* ***E.*** *A histogram shows that the p-values for the effect of dissection in the meta-regression are more weakly enriched in favor of significant findings than in* ***Figure S13****. 24 genes reached significance for effects. The full results can be found in* ***Table S19.***

**A.** Model: Y~ADType+Dissection
*ADType:* 0.5=non-traditional, -0.5=traditional
*Dissection:* 0.5=ACg, 0=Cortex that may contain Acg (FC), -0.5=Cortex not containing Acg or much ACg

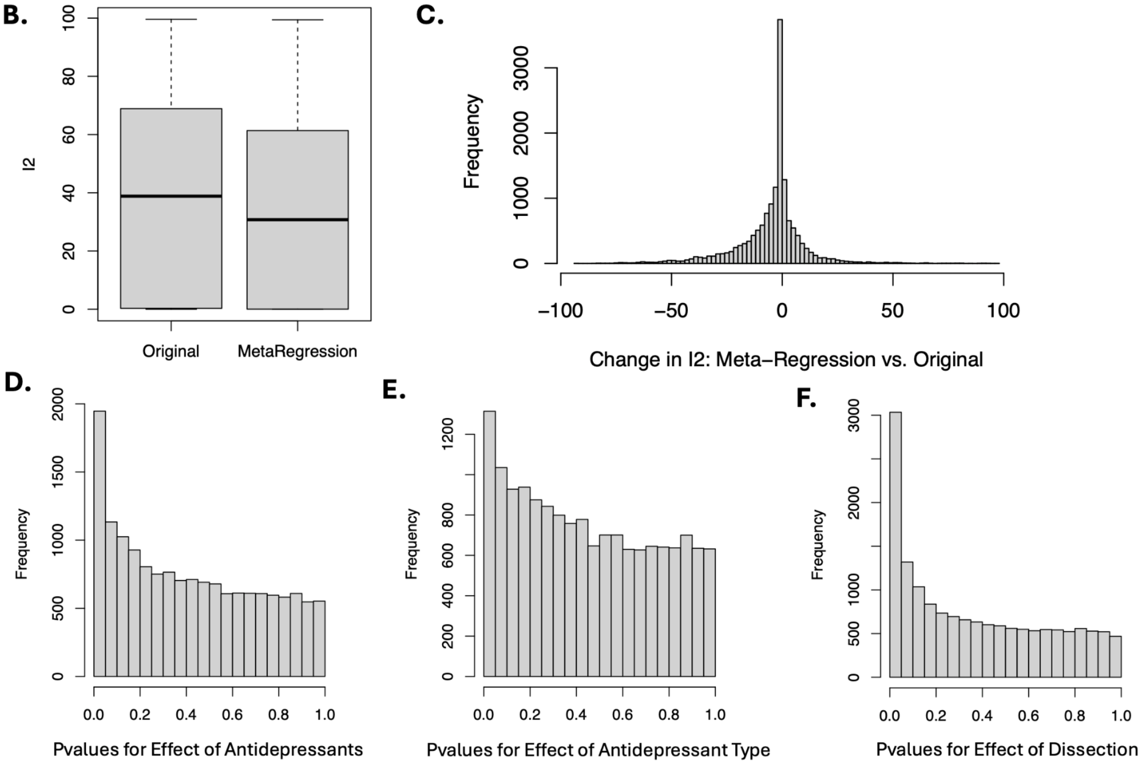


***Figure S14. A meta-regression including antidepressant type (non-traditional vs. traditional) and dissection (anterior cingulate (ACg) vs. other cortex) as co-variates provides extremely tentative insight into the heterogeneity in antidepressant effects observed in the cortical data. A.*** *The definitions used for the meta-regression model.* ***B-C.*** *Meta-regression decreased the residual heterogeneity present in the effects, as illustrated by a boxplot showing the $I^2$ values for all genes included in the original meta-analysis or meta-regression (**B**) or by a histogram illustrating the change in $I^2$ between the original meta-analysis and meta-regression (**C**).* ***D-F.*** *Histograms showing that the p-values for each of the variables in the meta-regression is enriched in favor of significant findings:* ***D.*** *Overall effect of antidepressants,* ***E.*** *Effect of antidepressant type (non-traditional vs. traditional),* ***F.*** *Effect of dissection (anterior cingulate (ACg) vs. cortex that may contain ACg (FC: frontal cortex) vs. other cortex)). The full results can be found in* ***Table S20.*** <u>*These results should be taken extremely tentatively due to the small number of antidepressant vs. control contrasts derived from ACg tissue (3 at most, less for some genes).*</u>

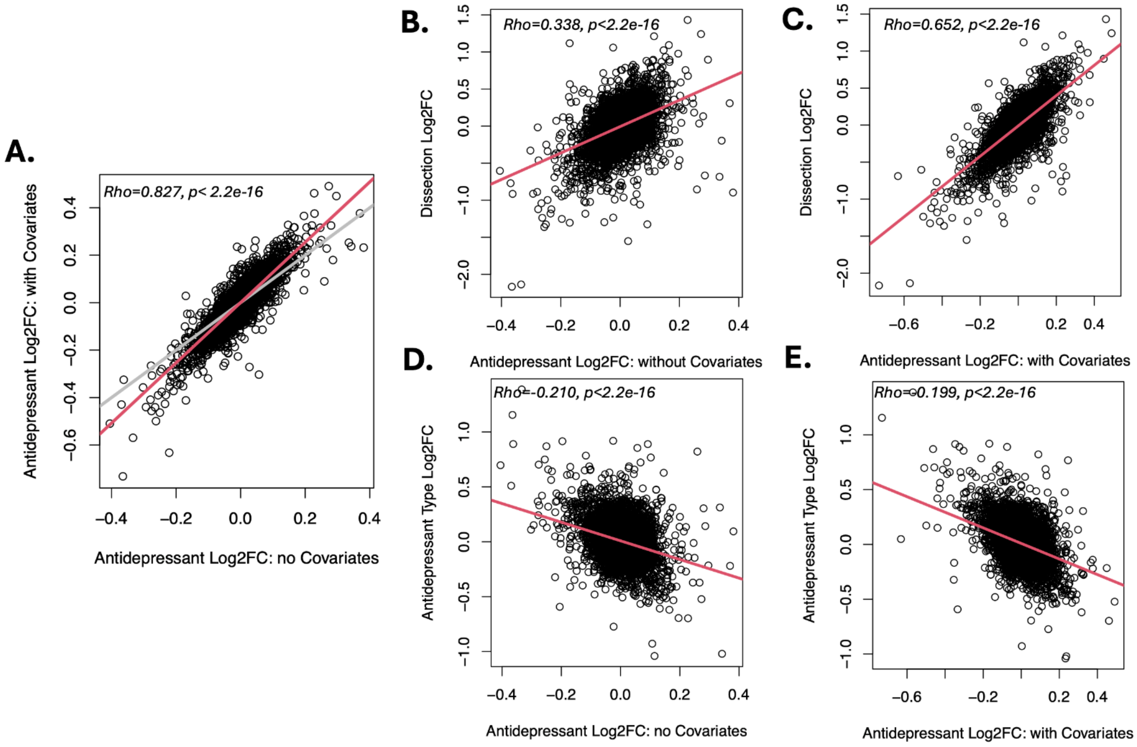


***Figure S15. A meta-regression including antidepressant type (non-traditional vs. traditional) and dissection (anterior cingulate (ACg) vs. other cortex) as co-variates suggests that antidepressant effects are larger in the anterior cingulate. A.*** *The antidepressant effects detected within the meta-regression with co-variates (Model:* ***Figure S13A****) tended to be larger than the antidepressant effects detected in our original meta-analysis. A scatterplot shows that the overall effect of antidepressants (Log2FCs) derived from our meta-regression (with covariates) correlates strongly with the effects of antidepressants (Log2FCs) derived from the original cortical meta-analysis (no covariates) (Rho=0.827, p<2.2e-16), showing a slope (red) that exceeds 1 (grey).* ***B.*** *A scatterplot shows that the effects of dissection (Log2FCs) derived from our meta-regression (ACg vs. other cortex) correlates positively with the effects of antidepressants (Log2FCs) derived from the original cortical meta-analysis (no covariates) (Rho=0.338, p<2.2e-16).* ***C.*** *A scatterplot shows that the effects of dissection (Log2FCs) derived from our meta-regression (ACg vs. other cortex) correlates positively with the overall effects of antidepressants (Log2FCs) derived from the meta-regression (Rho=0.652, p<2.2e-16).* ***D.*** *A scatterplot shows that the effects of antidepressant type (Log2FCs) derived from our meta-regression (non-traditional vs. traditional) correlates negatively with the effects of antidepressants (Log2FCs) derived from the original cortical meta-analysis (no covariates) (Rho=-0.210, p<2.2e-16), implying that the original meta-analysis was slightly skewed in favor of traditional antidepressants.* ***E.*** *A scatterplot shows that the effects of antidepressant type (Log2FCs) derived from our meta-regression (non-traditional vs. traditional) continue to correlate negatively with the overall effects of antidepressants (Log2FCs) derived from the meta-regression (Rho=-0.199, p<2.2e-16). These results should be taken extremely tentatively due to the small number of antidepressant vs. control contrasts derived from ACg tissue (3 at most, less for some genes).*

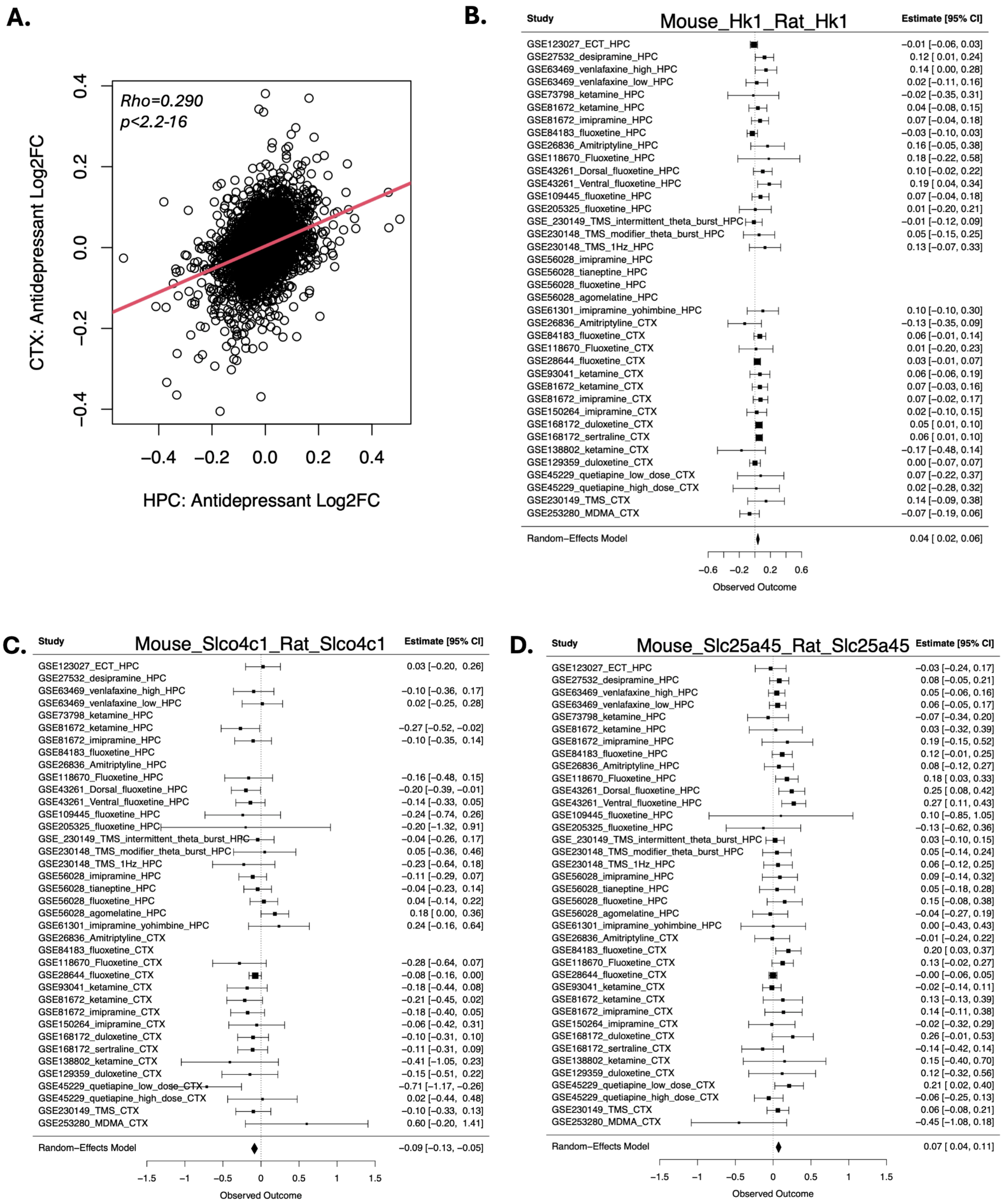


***Figure S16. A meta-regression including both the hippocampal and cortical data, with brain region included as a co-variate, identifies additional antidepressant-related gene expression. A.*** *A scatterplot showing that the effect of antidepressants (Log2FCs) derived from our original cortical meta-analysis (no*

*covariates) correlates with the effects of antidepressants (Log2FCs) derived from the original hippocampal meta-analysis (no covariates) when considering all genes included in both analyses (Rho=0.290, p<2.2-16).* ***B-D.*** *Twenty nine genes showed a significant overall effect of antidepressants (FDR<0.05) in a meta-regression including both the hippocampal and cortical data, with brain region included as a co-variate. Example forest plots illustrate the effects of antidepressants in both the hippocampus and cortex for:* ***B.*** *Hexokinase 1 (Hk1), an enzyme which catalyzes the first rate-limiting step of glucose metabolism (glycolysis). Hk1 is consistently upregulated by antidepressants.* ***C.*** *Solute Carrier Organic Anion Transporter Family Member 4C1 (Slco4c1), an organic anion transporter which is involved in the membrane transport of many molecules, including thyroid hormones, peptides, and many drugs.* ***D.*** *Solute Carrier Family 25, Member 45 (Slc25a45), a transmembrane transporter predicted to be located in the mitochondrial inner membrane. The full meta-analysis results can be found in* ***Table S21****.*
